# PROXINAUT: An Automation Platform for Rapid Hit-to-Degrader Discovery

**DOI:** 10.64898/2026.09.22.753586

**Authors:** Sean M. McKenna, Monika T. Gnatzy, Edith van der Nol, Femke J. Boxman, Chenlu Zhang, Xiaokang Jin, Jingming Liu, Jeroen Kortekaas, Xiaoyu Zhang, Sebastian J. Pomplun

## Abstract

Massive library-based platforms enable rapid early hit discovery, yet subsequent hit resynthesis and validation remain major bottlenecks in drug discovery. This delay is amplified when optimizing hits, or elaborating structures towards more advanced modalities, such as chemical inducers of proximity (CIPs). Here, we report PROXINAUT, an end-to-end workflow integrating self-encoded library-based hit discovery with an automated parallel solid-phase synthesis platform to streamline hit discovery, validation, optimization, and degrader development. As a demonstration, screening a 176k-member benzimidazole library against BRD4 yielded six hit candidates, which were obtained via parallel automated synthesis and validated as submicromolar binders. The automation workflow empowered a rapid structure activity relationship exploration around the whole scaffold and resulted in a variant with six-fold improved binding affinity. Next, we aimed to automatize the development of bifunctional degraders. Solid phase synthesis typically leaves a C-terminal amide as a synthetic artifact that can compromise druglike properties. We developed a strategy to repurpose this resin-attachment site, converting it into functional degrons featuring either FBXO31-targeting C-terminal amides or cereblon-binding cyclic imides. Operating without human intervention, our automated platform executes up to 16 synthetic steps across multiple parallel structures, enabling true *de novo* synthesis of full bifunctional scaffolds, where ligand, linker and E3 recruiter variations can all be explored within the same workflow. This resulted in cell-active BRD4 degraders with subnanomolar to nanomolar potency. Overall, we show how library selections can merge with multi-step parallel automation to convert hits directly into validated leads and advanced degrader modalities.

## Introduction

Conventional drug discovery campaigns rely on disjointed, resource-intensive technologies for hit identification, validation, and structure-activity relationship (SAR) optimization. Affinity-selection platforms, such as DNA-encoded libraries (DELs)^1–5^ and self-encoded libraries (SELs),^6–9^ substantially simplify and accelerate initial hit discovery, by enabling the screening of libraries of millions or billions of compounds all at once. However, subsequent resynthesis of individual compounds introduces a bottleneck in hit validation. This limitation is further compounded during the iterative design-make-test-analyze (DMTA) cycles required for downstream optimization.

Beyond mere ligand optimization, modern drug discovery increasingly involves the generation of more complex modalities to tackle challenging drug targets. Chemical inducers of proximity (CIPs) have emerged as one of the most transformative therapeutic modalities in drug discovery over the past decade. The development of proteolysis-targeting chimeras (PROTACs) established targeted protein degradation (TPD) as a strategy for selective depletion of disease-associated proteins and sparked widespread interest in proximity-driven pharmacology.^10–13^ More recently, the principles of induced proximity have been applied to engineer ternary complexes that direct diverse post-translational modifications of neosubstrates,^13–17^ or to induce complex-dependent functional effects.^18^

The generation of such functional CIPs requires systematic optimization of all components: the ligand for the protein of interest and it’s exit vector, the linker^19^ and the recruiter of the effector protein, e.g., an E3 ligase. An emerging strategy which helps to streamline this complex process is direct-to-biology (D2B) screening, where parallelized microscale conjugation chemistries have been used to combine ligands for proteins of interest with different linkers and E3 ligases.^20–22^ In bypassing compound purification, a major bottleneck in the DMTA cycle, D2B enables the rapid exploration of many CIP component combinations. Yet, this strategy is usually dependent on substantial synthetic investment in the generation of fully functionalized ligands. Furthermore, D2B limits screening libraries to one or two conjugation reactions using biocompatible reagent and solvent combinations.

In solid-phase synthesis (SPS), excess reactants, reagents, and soluble byproducts are removed by simple washing steps following each transformation, enabling efficient multistep synthesis without intermediate purification. Automated SPS has recently been demonstrated as a powerful source of small molecule libraries for screening.^23^ However, SPS, has been scarcely applied as a resource for CIP libraries,^24,25^ and to our knowledge, automated SPS has never been applied for the *de novo* generation of CIPs from readily available building blocks.

To establish PROXINAUT as a platform for rapid translation of encoded library hits into functional <u>prox</u>imity <u>in</u>ducers using <u>aut</u>omated synthesis, we ran an end-to-end workflow using the BET bromodomain-containing protein 4 (BRD4) as a demonstration case.^11,14,26^ A SEL screen of a 176k-member benzimidazole library resulted in the *de novo* identification of six hits which were rapidly prepared for validation by automated synthesis. Target binding was optimized through automated synthesis and SAR analysis of an additional 26 analogues, after which 33 CIPs were designed and synthesized bridging our high-affinity ligands through a variety of linker types to E3 ligase recruiters for F-Box protein 31 (FBXO31) and cereblon (CRBN), enabling discovery of subnanomolar cell-active degraders (Figure 1).

**Figure 1:**
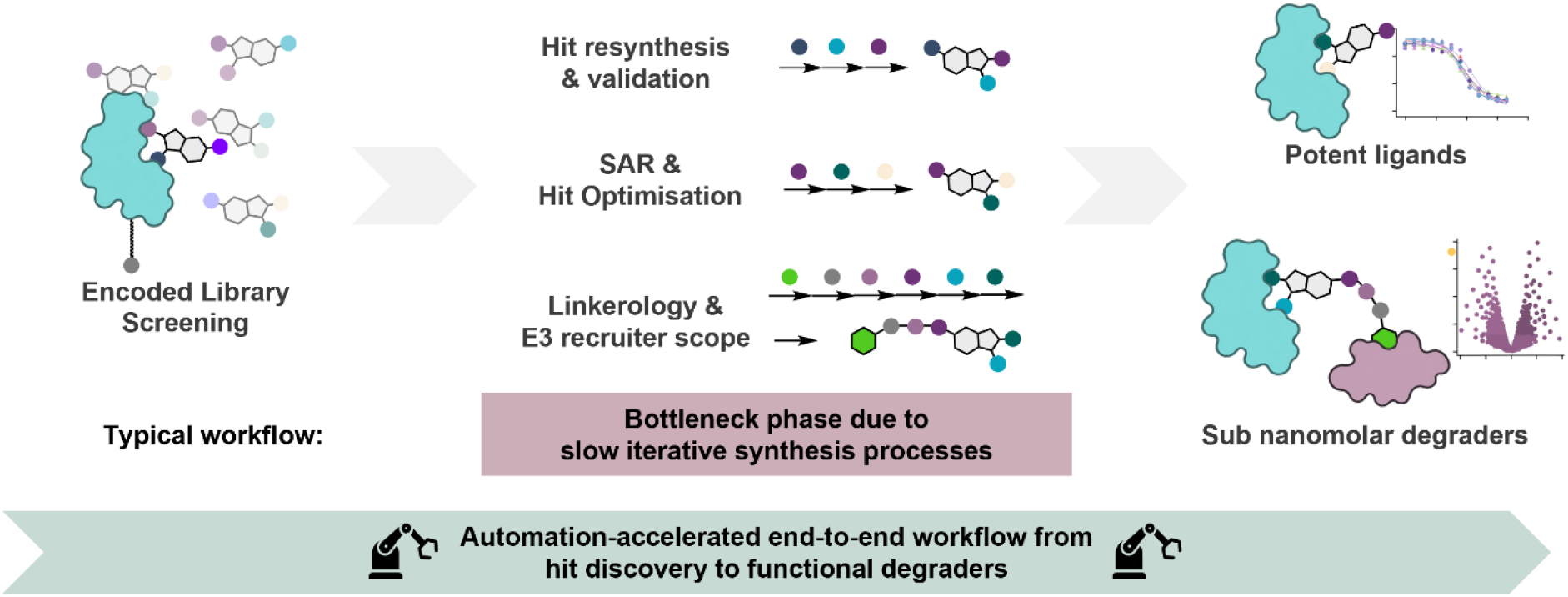
Harnessing automated synthesis as a tool for encoded library screening hit validation, optimization and elaboration to fully functionalized CIPs. In this study we developed an automated synthesis platform capable of performing multiple DMTA cycles to streamline hit-to-degrader workflow.

## Results & Discussion

### A SEL screen with a 176k-membered library identifies six putative binders of BRD4

To showcase our end-to-end workflow, from identification of novel binders, to validation, SAR and CIP development, we began by screening a combinatorial 176,384-member benzimidazole library (Figure 2a) in an affinity selection against biotinylated BRD4-BD1. The biotinylated target protein was immobilized on streptavidin-coated magnetic beads and incubated with the library (∼100 fmol/member)(Table S1-2). After removing non-binders, protein-bound hits were eluted under denaturing conditions and analyzed using our nanoLC-MS/MS SIRIUS-COMET workflow (Figure 2b).^8^ To account for non-specific binders, parallel selections were performed against carbonic anhydrase IX (CAIX) and WD-repeat containing protein 5 (WDR5). Compounds were classified as hits if they exhibited at least a three-fold enrichment in the BRD4 samples compared to the control proteins, and if MS2 fragmentation spectra could be matched in at least two out of three replicate samples. This SEL selection ultimately resulted in the identification of benzimidazole hits **1**-**6** (Figure 2c, Figure S1). Common motifs were found with saturated lipophilic ring systems at the N-1 position and dimethylphenol at the C-2 position, while greater variability was found at the C-5 carboxamide position. This hit family showed close resemblance to previously described benzimidazole based ligands for BRD4.^27,28^

**Figure 2:**
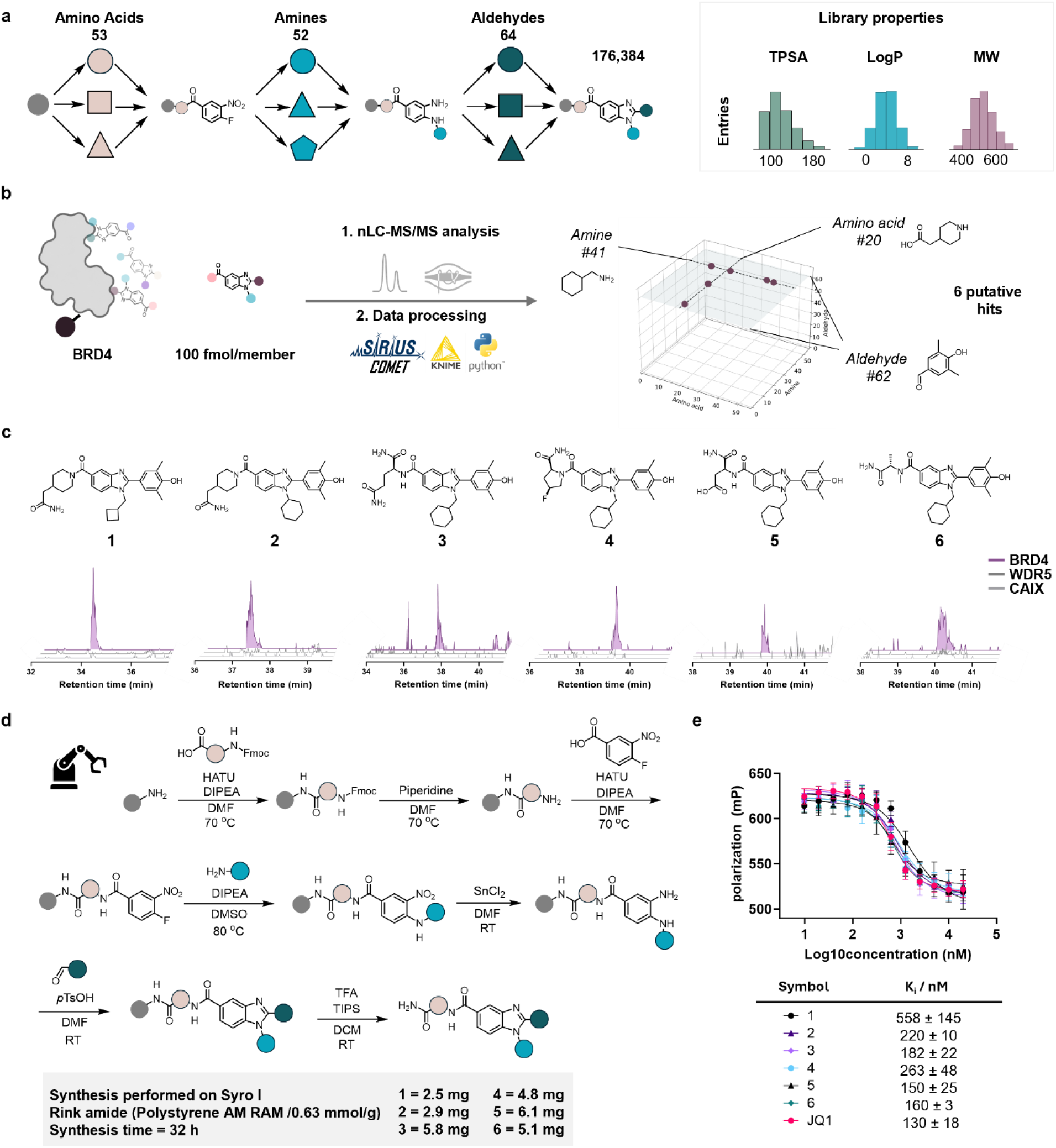
SEL-based BRD4 hit identification and automation-enabled resynthesis and validation. a) A 176k-member library was prepared by split-and-pool combinatorial chemistry using 53 amino acids, 52 primary amines and 64 aldehydes, b) An affinity selection was performed on BRD4. WDR5 and CAIX were used as negative controls. Hits were identified by nanoLC-MS/MS and decoded using SIRIUS COMET resulting in identification of six hits, with notable enrichment of building blocks, c) Benzimidazole hit structures and corresponding extracted ion counts using a 5 ppm accuracy of their corresponding masses in the BRD4 sample and WDR5 and CAIX controls. d) Summary of automated synthesisworkflow to generate compounds 1-6 for binding validation studies, e) Fluorescence polarization assay results for compounds 1-6 displacing JQ1-FITC binding to BRD4.

### Automated synthesis for streamlined SEL hit validation

With six putative BRD4 binders identified, we initiated the automated resynthesis of benzimidazole hits **1**-**6** in parallel for binding validation studies. In our automated synthesis workflow, Fmoc-protected amino acids were immobilized on polystyrene Rink-amide resin, deprotected, then coupled to the benzimidazole scaffold precursor 4-fluoro-3-nitrobenzoic acid (Figure 2d). Nucleophilic aromatic substitution with primary amine building blocks generated secondary anilines, and reduction of the 3-nitro group was achieved using tin(II) chloride. Finally, the diamine was treated with 4-hydroxy-3,5-dimethylbenzaldehyde in the presence of *p*-toluene sulfonic acid to initiate heterocyclization to the desired benzimidazole. Upon completion of the 32-hour automated synthesis procedure, compounds were cleaved from resin under acidic conditions and purified by automated reverse phase column chromatography, resulting in isolation of 2-6 mg of target compounds with overall yields of 8-23%.

To validate compounds **1**-**6** as true BRD4 binders, we measured displacement of a FITC-labeled JQ1 tracer in a fluorescence polarization (FP) assay along with JQ1 as a positive control. Dissociation constants were calculated from dose-response curves (Figure 2e), confirming that SEL hits **1**-**6** were all competitive submicromolar binders of BRD4, with compounds **3, 5** and **6** almost equaling the K^i^ of JQ1.

### Accelerating SAR analysis through automated generation of analogues

With an automation platform readily capable of generating multiple benzimidazoles in parallel (Figure 3a), we were interested in exploring the chemical space around our validated hits to better understand underlying structure-activity relationships, identifying opportunities to increase the potency of our series. Therefore, we initiated the automated synthesis of **7**-**32** (Table S3), systematically modifying each variable position around the benzimidazole core and evaluated their binding affinities by FP assay (Figure 3b, Figure S2).

**Figure 3:**
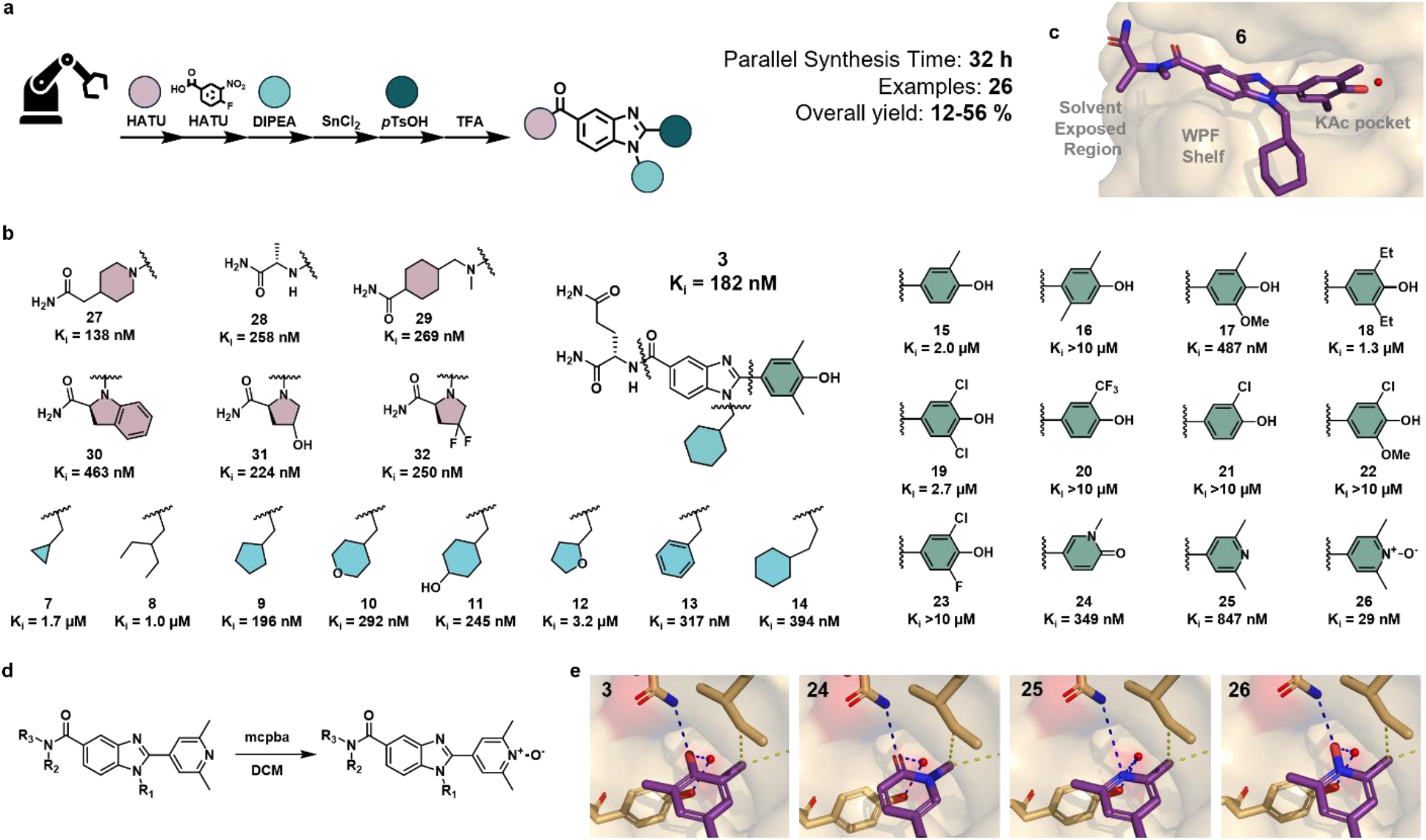
Structure-activity relationship exploration empowered by automated synthesis. (a) Parallel synthesis was used to generate 26 analogues of SEL hit **3** varying amino acid, amine and aldehyde building blocks, (b) Summary of analogues **7**-**32** as derivatives of SEL hit **3**, with their corresponding binding affinities determined by FP assay, (c)SEL hits were in the chemical space of a previously reported series of BRD4 ligands. Using previously published crystal structures (PDB: 6ZCI, 6TPX) we docked SEL hit **6** to rationalize protein-ligand interactions and design analogues to probe interactions with the KAc pocket, WPF shelf and the solvent-exposed region, (d) Bespoke pyridine N-oxidation for preparation of **26**, (e) A pharmacophore hybrid approach identified the novel dimethylpyridine N-oxide motif of **26** as a group capable of fulfilling essential interactions in the KAc pocket.^29^

In position N-1, contraction of aliphatic ring size from cyclohexylmethylamine **3** to cyclopropylmethylamine **7** resulted in decreased binding affinity, consistent with reduced engagement with a lipophilic patch formed from the neighboring residues Trp81, Pro82 and Phe83, commonly known as the ‘WPF shelf’, adjacent to the BRD4 pocket (Figure 3c). Acyclic **8** gave five-fold lower binding affinity than the corresponding cyclopentane **9**. Inclusion of an endocyclic or exocyclic heteroatom in pyran **10** and cyclohexanol **11** were both moderately well tolerated, while tetrahydrofuran **12** exhibited substantial loss in binding affinity. Finally, desaturating the ring system (**13**) or extending the linker (**14**) resulted in only modest losses in binding affinity.

Next, we evaluated modifications at the C-2 position. We found that removal of one methyl substituent (**15**) resulted in greater than ten-fold loss in binding affinity, while methyl migration (**16**) abolished protein binding altogether. While very minor modifications such as mono-methoxylation (**17**) were tolerated, more considerable changes to arene substitution strongly decreased (**18**-**19**) or completely abolished binding (**20**-**23**). Previously described N-methylated pyridone^21,27,28^ **24** exhibited a K^i^ of 349 nM, which was notable given that the corresponding building block had been included in the combinatorial synthesis of the SEL but not identified as a hit. This result highlights that ionization and fragmentation characteristics may bias hit identification during SEL decoding.

Motivated to identify novel motifs with H-bond acceptors capable of engaging Asn140 in the KAc pocket, we proposed dimethylpyridine N-oxide as a pharmacophore hybrid which could be accessed via late-stage oxidation of dimethylpyridine **25** (Figure 3d). Pleasingly, **26** gave a binding affinity of 29 nM, corresponding with a six-fold binding affinity improvement above parent hit **3**, nearly thirty-fold greater potency than precursor **25**, and four-fold greater affinity than JQ1. To our knowledge, this is the first application of a pyridine N-oxide ligand being used to engage the KAc pocket of BRD4 (Figure 3e).

Finally, we sampled a smaller set of amino acids at the C-5 carboxamide, as this variable position was understood to least significantly influence binding affinity. We first prepared consensus compound **27**, containing the most commonly enriched building blocks from our SEL screen, and were rewarded with a slight improvement in binding affinity (K^i^ = 138 nM). However, the incorporation of alternative amino acids into **28**-**32** resulted in mild losses in binding affinity.

At the conclusion of our SAR analysis, we observed that the overwhelming majority of benzimidazole analogues gave weaker binding affinities than our original SEL hits. Notably, only compound **26**, which required a rational drug design strategy and an additional bespoke synthetic transformation, and **27**, offered improved BRD4 binding affinity. Given that many of the individually tested building blocks were part of the initial combinatorial library design, these findings underscore the good, though not complete, coverage of the best building block combinations in our SEL workflow.

### Translating small molecule hits-to-degraders

While heterobifunctional molecules such as PROTACs hold immense therapeutic potential, their physicochemical optimization remains a formidable challenge. Combining two distinct ligands connected by a linker routinely pushes these structures well beyond traditional rule-of-5 guidelines, making the minimization of molecular weight, polar surface area, and hydrogen bond donors critical for cell permeability.^30,31^ SPS introduces an inherent structural constraint: resin cleavage via standard amine- or hydroxyl-based linkers (such as the Rink amide used in our workflow) leaves behind a vestigial C-terminal amide or carboxylic acid “scar”. In a conventional SPS workflow for bifunctional discovery, these appendage groups are structurally unnecessary liabilities that add polarity, hydrogen bond donors and acceptors, and molecular weight. Although engineered cleavable linkers can circumvent this,^24,25,32,33^ they are often bespoke and can be poorly compatible with long, multi-step automated synthetic sequences. Rather than treating the resin attachment point as a synthetic limitation, we sought to directly co-opt this cleavage site as a component of the functional E3 ligase recognition motif.

We implemented two distinct design strategies to leverage this resin-attachment point for E3 ligase recruitment, both compatible with our automated parallel synthesis workflow. First, we capitalized on recent discoveries identifying C-terminal amides as physiological degron motifs recognized by the F-box protein FBXO31 (Figure 4a), allowing us to seamlessly integrate E3 engagement directly into our resin design without auxiliary functional groups. Second, inspired by recent insights into cyclic imide-mediated cereblon (CRBN) recruitment (Figure 4b), we engineered a strategy to harness intramolecular aspartimide and glutarimide formation. By incorporating terminal aspartate or glutamate residues into our automated parallel workflow, we can directly transform the resin-derived terminal position into a cyclic imide degron for CRBN. Both of these strategies and their implementation within the PROXINAUT platform are outlined in detail in the following sections.

**Figure 4.**
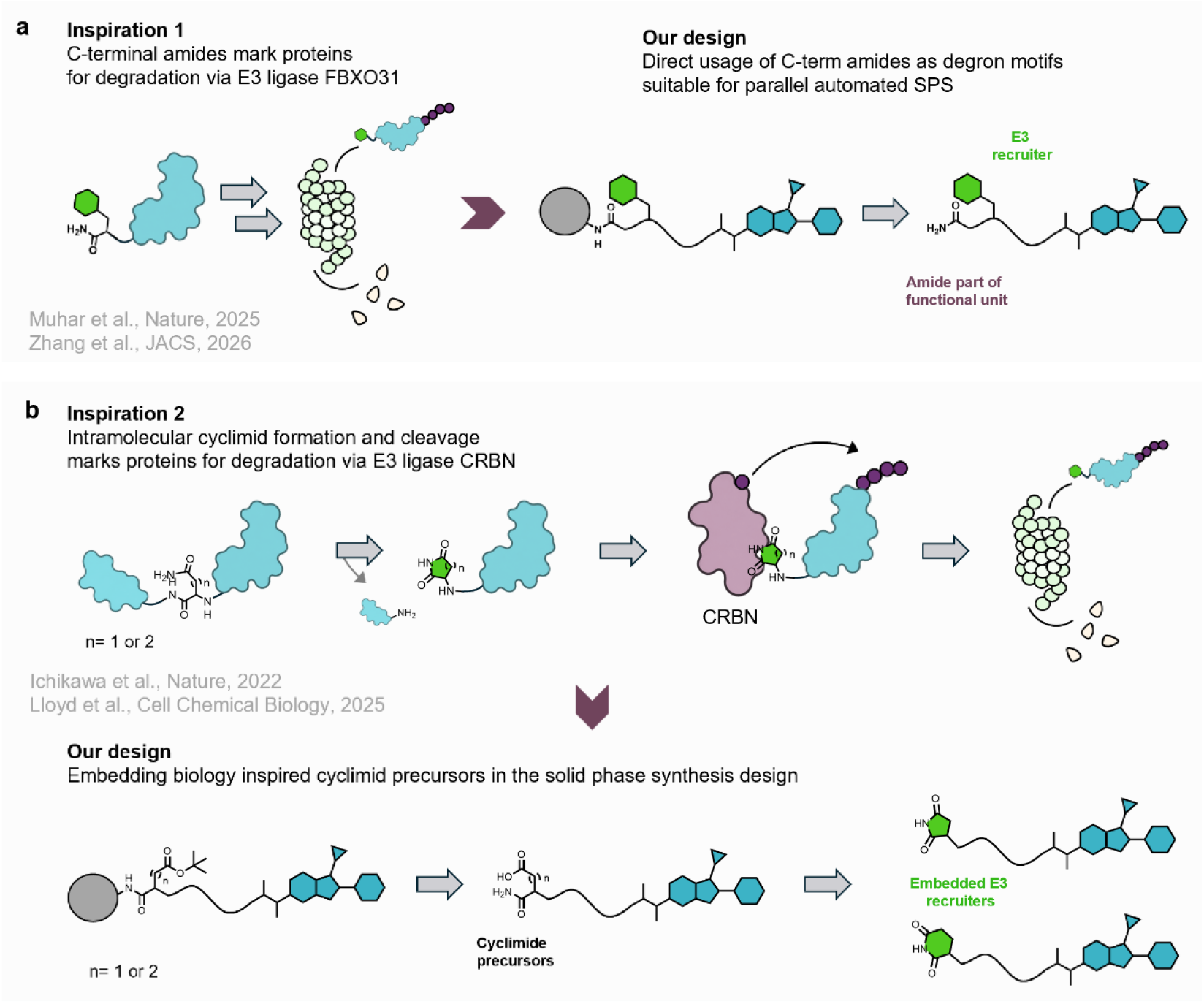
Bioinspired degron motifs enable practical E3 recruiter embedding via automated solid phase synthesis. Common solid phase synthesis leads to an additional functional group as a remnant of the solid phase attachment. Avoiding such a “scar” becomes especially important for the design of modalities like PROTACs which often do not conform to rule-of-5 criteria. (a) C-terminal amides generated by SPS were used as part of the E3 ligase recruiting motif inspired by the natural recognition motif of the E3 ligase FBXO31, (b) Cyclic imide formation as a marker for protein degradation via the E3 ligase CRBN served as an inspiration for the synthesis of CIPs by cyclisation of C-terminal Asp or Glu.

### Hit-to-degrader development using FBXO31 recruiters

Recently, Muhar *et al*. reported that terminal amides are a degron recognition motif for F-Box protein 31 (FBXO31), a substrate receptor of the ubiquitin ligase complex SKP1-CUL1-F-box.^34^ In a follow-up study, Zhang *et al*. identified C-terminal amide-phenylalanine-alanine (Phe-Ala) as an optimal recruiter, applying this dipeptide motif in PROTACs for FBXO31-dependent degradation of FKBP and BRD4.^35^ C-terminal installation of Phe-Ala appeared to be a readily tractable strategy for accessing bifunctional degrader libraries using our automated SPS platform (Figure 5a).

**Figure 5:**
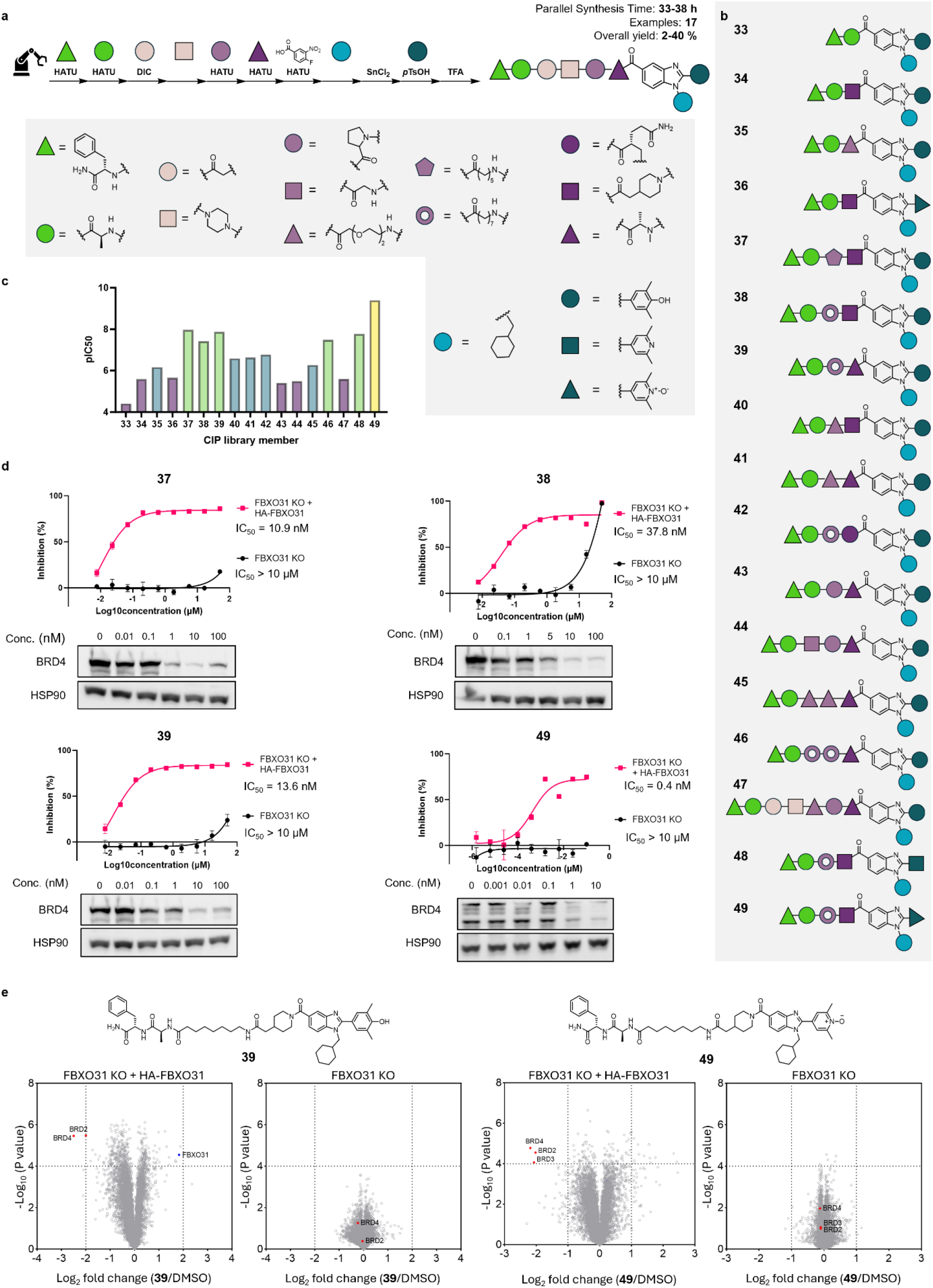
Hit-to-degrader discovery for FBXO31-recruiting CIPs. (a) Automated synthesis workflow for preparation of CIPs **33**-**49**, (b) Schematic representation of FBXO31-recruiting CIPs **33**-**49**, (C) pIC_50_ cell viability values determined for CIPs **33**-**49** in FBXO31 KO HEK293 cells re-expressing FBXO31, (d) Cell viability IC_50_ values determined for CIPs **37, 38, 39** and **49** in FBXO31 KO HEK293T cells and FBXO31 KO cells re-expressing FBXO31,and dose-dependent BRD4 degradation determined by Western blotting, (e) Global proteomic analysis of CIPs **39** and **49** in FBXO31 KO HEK293 cells re-expressing FBXO31. No enrichment or stabilization of BRD4/2 or FBXO31 can be seen in FBXO31 KO HEK293T cells.

Therefore, we designed a library of FBXO31-recruiting CIPs featuring our highest affinity BRD4 binders and sampling a range of linker lengths and chemistries (Figure 5b, Table S4). Due to building block accessibility, we retained the dimethylphenol group at the C-2 position in the majority of cases, while still preparing select dimethylpyridine and dimethylpyridine N-oxide-containing examples for matched-pair analysis. Compounds **33**-**47** were prepared by automated synthesis. Up to 16 steps were performed via full automation, in parallel on multiple variants and upon flash purification we obtained all compounds in high purities and in sufficient quantity for further bioassays (1mg – 20mg).

We screened for functional activity by examining dose-dependent effects on cell viability in FBXO31 knockout (KO) HEK293T cells and FBXO31 KO cells re-expressing FBXO31 (Figure 5c, Figure S3). Because BRD4 degradation has a more drastic effect on cell viability than simple competitive binding to the bromodomain pocket, a differential response in this assay served not only as an indicator of overall cellular activity, but also provides a direct indication of whether the compound acts via targeted protein degradation. C5 linked **37**, and C7 linked **38** and **39** gave a strong FBXO31-dependent dose-response. Meanwhile, constrained linker compounds **43** and **44** demonstrated lower degradation activity, revealing that flexible linker CIPs gave more effective degradation. Based on the most promising linker identity, we prepared variants **48** and **49** featuring dimethylpyridine and dimethylpyridine N-oxide C-2 substituents respectively. Both series exhibited strong differential activity, with dimethylpyridine N-oxide **49** emerging as the most potent variant of the series in accordance with our highest affinity BRD4 binder, **26**.

With the most promising variants **37, 38, 39** and **49**, we proceeded to measure the BRD4 degradation by Western blotting in FBXO31 re-expressing cells incubated for 24 hours (Figure 5d). Dose-dependent depletion in BRD4 was observed and degradation activity was calculated in the subnanomolar to low nanomolar range. Global proteomic analysis in the same cell line found BRD4, along with its paralogue BRD2, underwent FBXO31-dependent depletion in the presence of **39** and **49** (Figure 5e, Figure S4), with concomitant stabilization of FBXO31, as previously reported.^35^

### An embedded cyclic imide precursor for cereblon-recruiting CIPs

Having verified PROXINAUT could function as a platform for rapid hit-to-degrader discovery, we aimed to develop a synthetic strategy enabling us to embed an alternative degron recognition motif for one of the most well-characterised E3 ligases in the TPD field: cereblon (CRBN).

Pioneering studies from Ichikawa *et al*. established that intramolecular cyclisation of asparagine or glutamine residues on peptide backbones generates C-terminal cyclic imides, the degron recognition motif for CRBN.^36,37^ We were inspired to mimic this transformation while inverting the reaction partners, installing activatable aspartic or glutamic acid residues which could undergo nucleophilic attack from the C-terminal amide of the resin attachment point (Figure 6a).

**Figure 6:**
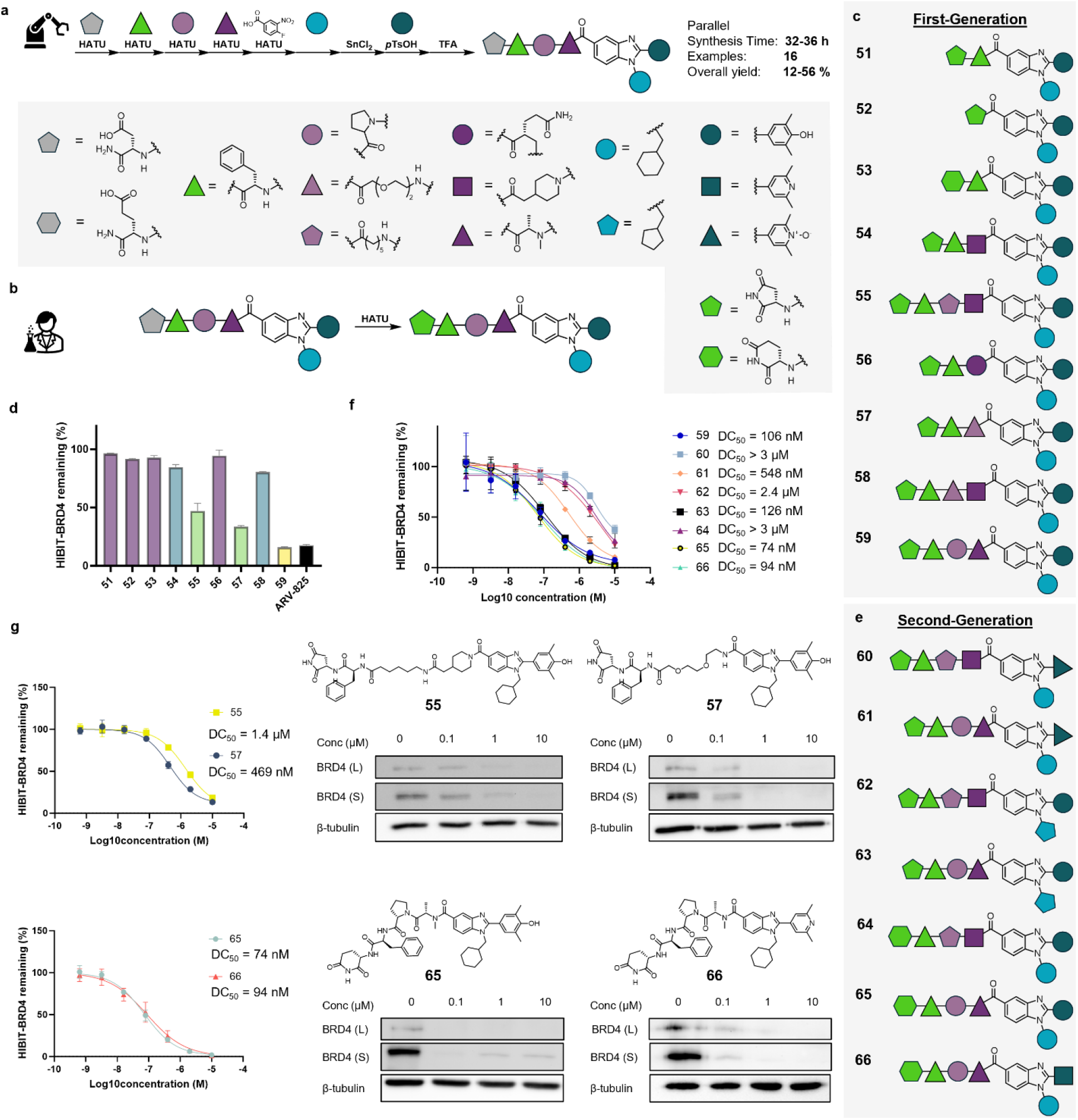
Hit-to-degrader discovery for CRBN-recruiting CIPs, (a) Workflow for preparation of CIPs **51**-**66** from corresponding building block synthons, with a single manual step for cyclic imide formation (b) A model system for cyclic imide formation from C-terminal glutamic acid, (C) Schematic representation of first-generation CRBN-recruiting CIPs **51**-**59**, (d)HiBiT-BRD4 luminescence for first-generation CIPs **51**-**59** screened at 1µM, (e) Schematic representation of second-generation CRBN-recruiting CIPs **60**-**66**, (f) HiBiT-BRD4 luminescence for second-generation CIPs **51**-**59** screened in full concentration curves, (g) BRD4 DC_50_ values for CIPs **55, 57, 65** and **66** determined by HiBiT assay and anti-BRD4 Western blotting.

To explore the feasibility of this strategy, we initiated the automated synthesis of model benzimidazole **50** (Figure 6b, Figure S5), featuring aspartic acid-phenylalanine (Asp-Phe) at the C-terminus as a precursor to a previously described peptidic cereblon recognition motif.^36^ We cleaved the precursor compounds and tested cyclisation using CDI and HATU to generate activated ester electrophiles which could undergo spontaneous C-terminal cyclisation.^38^ While the desired cyclic imide **51** was observed to form in both test reactions, HATU was found to give cleaner conversion (Figure S6), and hence was predominantly favored method for cyclisation.

Using automated synthesis followed by a single step solution-phase ring-closure reaction, we synthesized cyclic imides **51–59** (Figure 6c, Table S5). As aspartimides are prone to epimerization, we endeavored to resolve the resultant diastereomers as major (putative *S*-aspartimide) and minor (putative *R*-aspartimide) products. We tested all compounds at 100 nM and 1 µM in HEK293 cells expressing HiBiT-BRD4, measuring degradation using the luminescence signal after 24 hours of incubation (Figure 6d), confirming that our putative *S*-aspartimides were more potent degraders in all instances, consistent with previous studies (Figure S7).^39^

Anti-BRD4 Western blotting of compounds **51**-**59** revealed good agreement with results obtained using the HiBiT-BRD4 (Figure S8). Minimalist variants **51**-**53** demonstrated little to no depletion of BRD4, presumably due to linker lengths being too short for ternary complex formation. **54**-**55** showed clear dose-depletion in luminescent signal, consistent with BRD4 degradation. Once again, a glutamide-bearing variant, **56**, demonstrated very low activity likely due to excessive hydrophilicity. CIPs **57** and **58** retained reasonable degradation activity, but PEG-linker motifs appeared to generally correspond with lower cellular activity. Meanwhile, conformationally constrained CIP **59** showed exceptional degradation activity. This result demonstrated a clear difference in linker preference between CRBN-recruiting **59** and FBXO31-recruiting **49**.

Harnessing the ability of our automated synthesis workflow to enable modification of each component of the bifunctional degraders in the same workflow, **60**-**66** were proposed as a second-generation of CRBN-recruiting CIPs (Figure 6e). We iterated upon flexible **55** and constrained **59**, but varied the cyclic imide recruiter to include both 5-membered and 6-membered examples. We also integrated examples of dimethylpyridine and dimethylpyridine N-oxide at the C-2 position, as well as examples of cyclohexylmethanamine and cyclopentylmethanamine at the N-1 position (Table S5).

We tested these second-generation compounds along with the most promising CIPs from our first-generation in full concentration curves in the HiBit-BRD4 assay (Figure 6f) along with Western blotting (Figure S9). To our surprise, dimethylpyridine N-oxides **60** (DC_50_ > 2.0 µM) and **61** (DC_50_ = 548 nM) underperformed phenolic matched-pairs **55** (DC_50_ = 1.4 µM) and **59** (DC_50_ = 106 nM), highlighting a potential cell-permeability issue for the N-oxide bearing examples in this series. Cyclopentylmethanamines **62** (DC_50_ = 2.4 µM) and **63** (DC_50_ = 126 nM) gave slightly reduced degradation activity, correlating with their reduced BRD4 binding in our FP assay. Most intriguingly, we discovered that inclusion of a 6-membered CRBN recruiting cyclic imides bestowed opposite results on compounds **64** and **65**. While flexible linker-containing **64** gave reduced degradation activity (DC_50_ > 2.0 µM), inclusion of the same E3-ligase recruiter conferred higher degradation activity on constrained linker-containing **65** (DC_50_ = 74 nM). This example demonstrates the cooperative effects of E3 ligase ligands and linkers on functional ternary complex formation, and highlights the value of having rapidly accessible screening libraries to identify non-additive SAR. Finally, we identified that, mirroring trends in our FBXO31-recruiting degraders, dimethylpyridine-containing **66** (DC_50_ = 94 nM) exhibited high cellular activity comparable with dimethylphenol **65** (DC_50_ = 74 nM) (Figure 6g), despite the five-fold binding affinity difference between parent ligands **3** and **25**. Notably, through exclusion of the phenolic H-bond acceptor, **66** does not present the vulnerabilities to phase II metabolism as **65**, further highlighting the potential of this KAc pocket-binding chemotype for lead optimization.

To explore whether BRD4 degradation was the result of our designed CRBN-recruiting cyclic imide, we proceeded to screen the acyclic precursors of **57, 59, 65** and **66**, some of our highest activity first-generation and second-generation degraders. **67**-**70** exhibited no measurable degradation activity (Table S6, Figure S10), supporting CRBN-dependent degradation. Taken together, our solid phase based automated synthesis combined with late-stage cyclic imide formation enabled practical library synthesis and the development of low nanomolar degraders.

## Discussion and conclusion

In this work, we presented a practical end-to-end workflow spanning from rapid hit discovery to automation-based compound optimization and elaboration to bifunctional degrader modalities with potent cell activity. As a proof-of-concept for this PROXINAUT platform, we identified six small molecule binders for the transcriptional regulator BRD4 through SEL selections from a library with 176k benzimidazoles. Our automated parallel synthesis workflow enabled rapid hit validation, SAR interrogation and elaboration to bifunctional degrader modalities recruiting FBXO31 and CRBN. Overall, we prepared and tested over 70 variants, including automated synthesis procedures with 16 steps without any manual intervention.

Our study bridges the disconnect between rapid ultrahigh-throughput hit discovery platforms, such as encoded libraries, and the downstream bottlenecks of hit validation and elaboration. Because combinatorial libraries are inherently built from modular building blocks, they provide a natural foundation for automated synthetic workflows. We anticipate that this strategy, coupled with the expanding chemical space accessible via automated parallel synthesis,^23^ will serve as a powerful resource for both academic and industrial drug discovery communities utilizing library-based screening. Beyond SELs, we expect that our described procedures will be directly applicable for many DEL projects. Building on these advantages, our laboratory has already begun routinely deploying these automated workflows for early hit validation across multiple projects, achieving rapid access to validated compounds with minimal manual workload.

While our study was focused on the accelerated translation of SEL hits to functional degraders, we see vast potential for PROXINAUT-inspired workflows for the discovery of other proximity-inducing modalities. Among others, RIPTACs, DUBTACs, DD-TACs and PhosTACs all share the bifunctional architecture of PROTACs and with that face the same need for optimizing multiple molecular components.^13–18^ Automated SPS can support the advancement of these CIP modalities.

Solid phase synthesis by its nature needs an attachment point for compounds, typically leaving a ‘scar’ in the final product. Our study featured Rink amide as a commonly utilized acid labile linker for SPS, resulting in a terminal amide in each final compound. To overcome the potentially negative effect on druglikeness of our screening compounds, we engineered two strategies to directly embed or elaborate the cleavage adducts into degron motifs. While Rink amide linkers are highly versatile and enable high-quality synthesis under a broad range of conditions, we are intrigued to see the adoption of alternative linker types for the preparation of screening libraries, and to observe how immobilization and release strategies are applied to embed functional components of CIPs.

Head-to-tail *de novo* synthesis of CIPs empowers complete flexibility in building block incorporation, enabling holistic sampling of variations of all components of complex bifunctional modalities. As seen in this study, combining variations in E3 recruiter identity, BRD4 ligand and linker can lead to unexpected non additive synergies. PROXINAUT’s ability to bring along analogues with variations in multiple positions during our hit optimization cycle enabled dimethylpyridine **66**, whose parent ligand gave only modest results in our FP assay, to be identified as a highly potent, cell-active BRD4 degrader. This highlights a key point of differentiation between PROXINAUT and conventional PROTAC synthesis, where protein-of-interest and E3 ligase ligands are commonly fixed prior to linkerology interrogation.^20–22,24,31,32^

In this study, parallel automated synthesis was followed by individual purification of each compound. Given our scale, this approach was feasible and did not impose a substantial workflow bottleneck. However, as the field of direct-to-biology (D2B) screening rapidly advances to evaluate crude reaction mixtures in cells,^40–48^ we anticipate that our workflow can be adapted to a full D2B setup, obviating the need for final column chromatography. Unlike conventional D2B workflows, where reaction byproducts accumulate and restrict procedures to one or two synthetic steps using biocompatible reagents, solid-phase synthesis inherently removes all soluble byproducts during intermediate washing steps. Consequently, a multistep D2B workflow built on this platform would only require recovering the product from the final resin-cleavage mixture, which can be streamlined, e.g., via parallel solid-phase extraction. While similar procedures are already established for peptidic D2B screens,^49,50^ our study demonstrates how this scope can be successfully extended to diverse, non-peptidic molecular scaffolds. We expect that bead-to-biology platforms will find more and more applications in hit validation, direct screening campaigns and the exploration and development of CIP libraries.

## Supporting information

Supporting Information: PROXINAUT

## Supporting Information

Supporting Information has been provided for this manuscript, composed of:

Supplementary figures; supplementary tables; abbreviations; experimental; building block libraries; synthetic sequences; analytical data (HPLC−MS and NMR); biological activity data; references

## Acknowledgements

Research reported in the publication was supported by Oncode Accelerator, a Dutch National Growth Fund project under grant number NGFPO2201. SJP and MTG acknowledge funding via the ERC Starting Grant (101039354). The Pomplun Lab gratefully acknowledges financial support from Mr. H. J. M. Roels through a donation to the Oncode Institute and KWF’s financial support of the Oncode Institute. The HEK293 HiBiT BRD4 cell line was a kind gift from the lab of Professor Alessio Ciulli of the University of Dundee.

## Notes

### Competing Interest Statement

The authors have declared no competing interest.

