## Supporting Information: PROXINAUT for "PROXINAUT: An Automation Platform for Rapid Hit-to-Degrader Discovery"

**Table of Contents**

1. Supplementary Figures 3
2. Supplementary Tables 12
3. Abbreviations 17
4. Experimental 19
    a. Reagents and Supplies 19
    b. Instrumentation 19
    c. Software 20
    d. Protein Preparation 20
    e. Fluorescence Polarization 22
    f. Cell Lines and Cell Culture 23
    g. Western Blot 24
    h. Global Proteomics 25
    i. Library Enumeration 27
    j. Library Synthesis 28
    k. Affinity Selection Procedure 29
    l. Data Processing 29
    m. Automated Parallel Synthesis 31
    n. Manual Transformations 32
    o. Pharmacophore Hybrid Modelling 33
5. Building Block Libraries 34
    a. Amino acids 34
    b. Primary amines 35
    c. Aldehydes 36
6. Synthetic Sequences 37
    a. Synthetic Sequence for Compounds **1-32** 37
    b. Synthetic Sequence for Compounds **33-49** 38
    c. Synthetic Sequence for Compounds **51-66** 39
    d. Synthetic Sequence for Compounds **50**, **67-70** 39
7. Analytical Data 40
    a. HPLC-MS Data 40
    b. $^{1}$H-NMR Data 77
8. Biological Data 82
    a. Uncropped Western blots for compounds **37- 39**, **48** and **49** 82
    b. Uncropped Western blots for compounds **51- 59** 83
    c. Uncropped Western blots for compounds **60- 70** 84
9. References 85

### Supplementary Figures

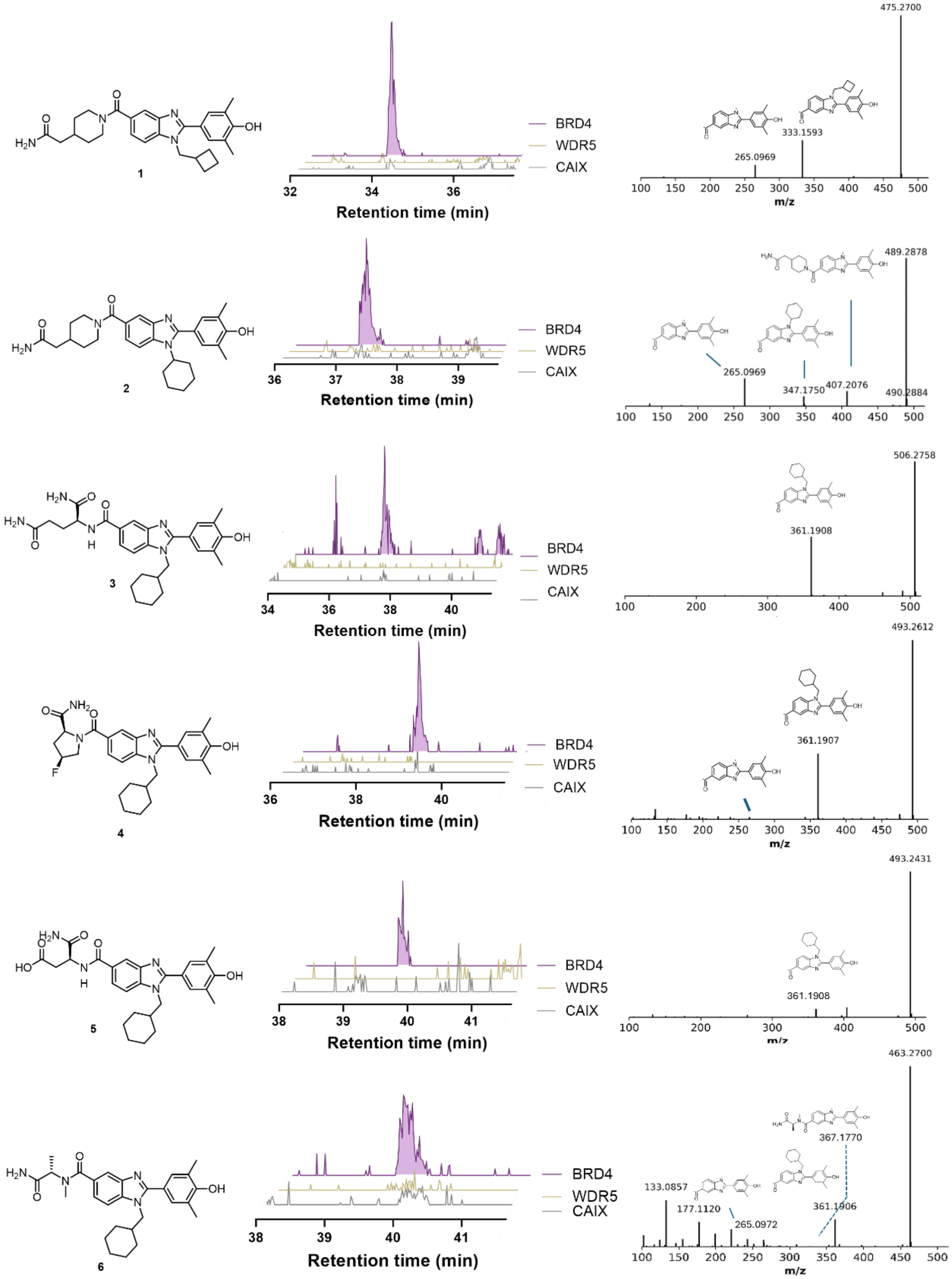

**Figure S1:** Chemical structures of hits **1**-**6** identified using COMET, with the extracted ion chromatogram (EIC) and LC-MS/MS chromatogram from the affinity selection of SEL against BRD4.

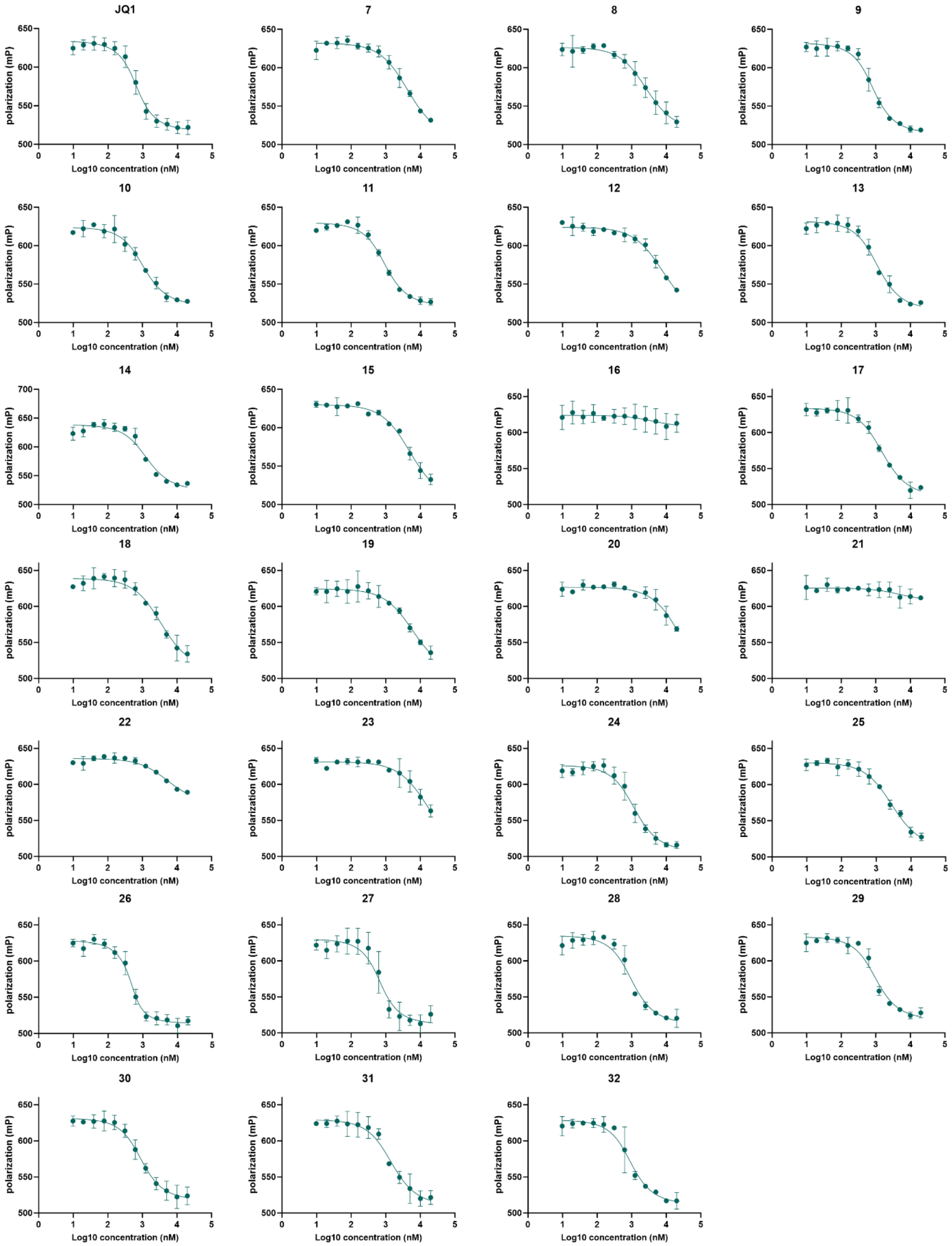

**Figure S2:** Fluorescence polarization assay results for **JQ1** and benzimidazole hit derivatives **7**-**32**.

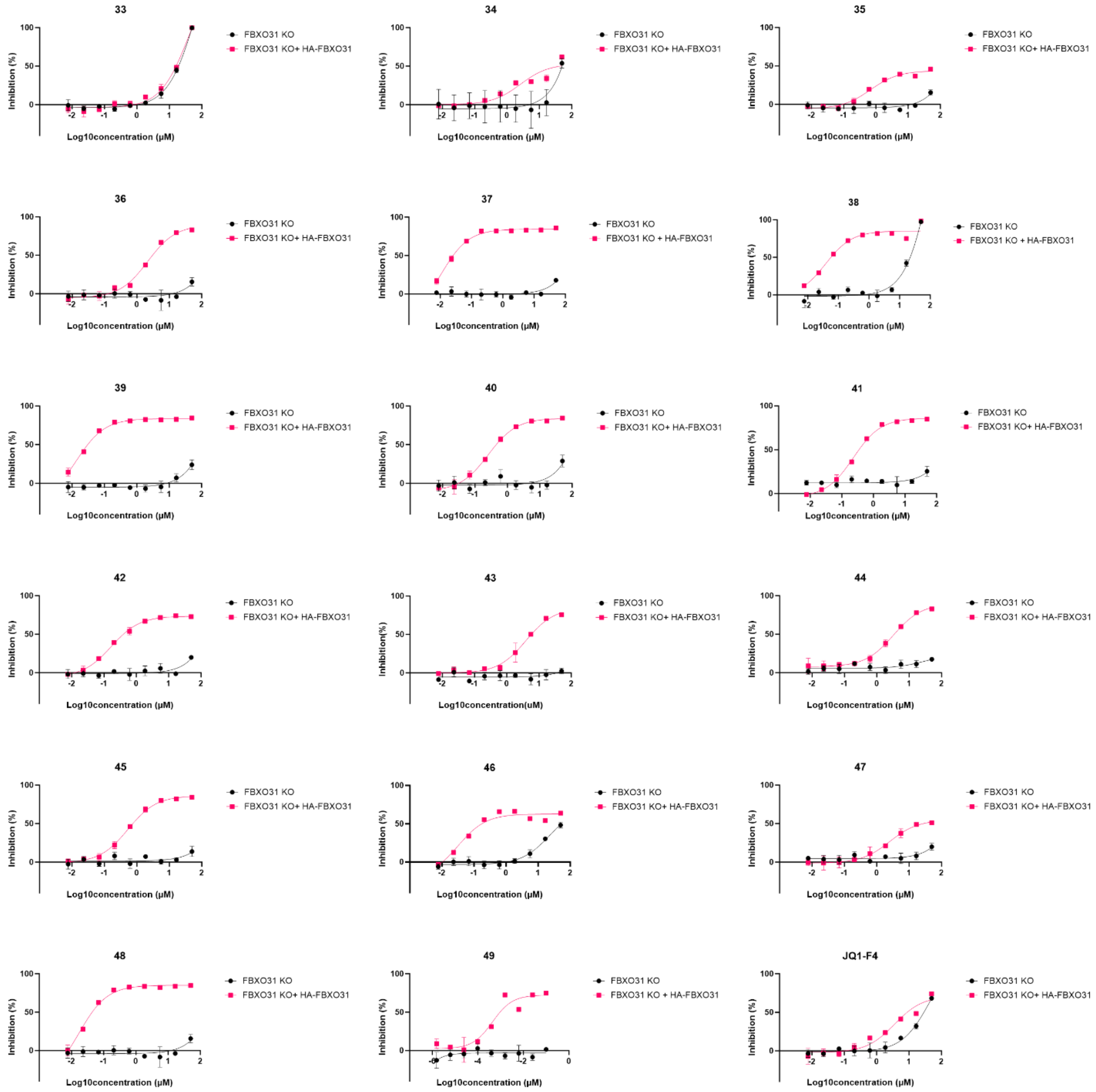

**Figure S3:** Cell viability analysis for CIPs **33**-**49** and JQ1 in FBXO31 KO HEK293T cells and FBXO31 KO cells re-expressing FBXO31 after treatment for 72h. Data represent mean values (n=3 replicates).

**Compound 39**

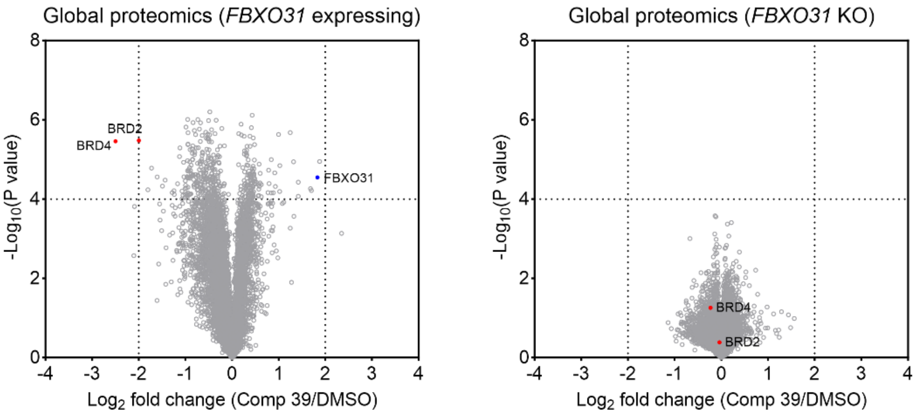

**Compound 48**

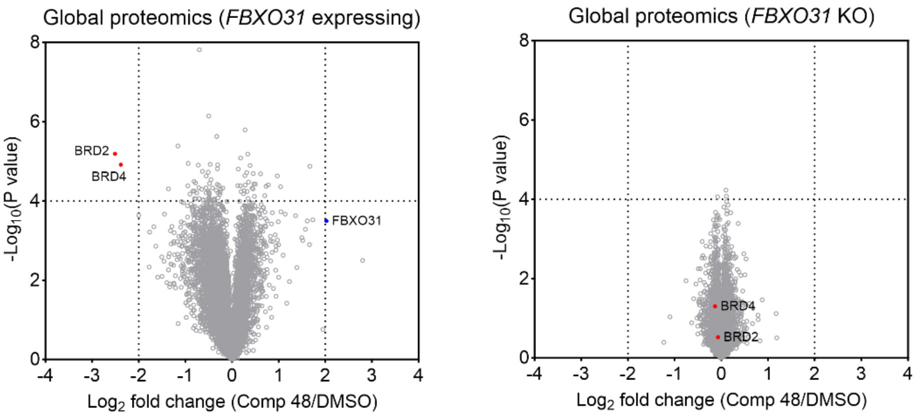

**Compound 49**

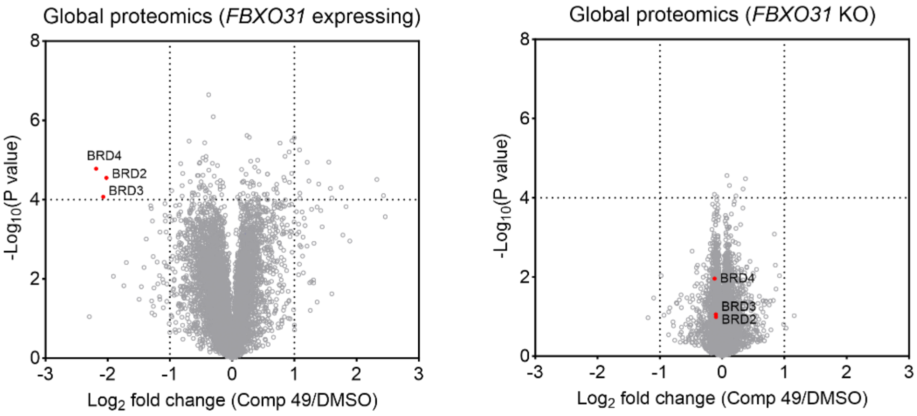

**Figure S4:** Proteomic profiling of **39**, **48** and **49**-induced target degradation in HEK293T cells. Volcano plots showing global proteome changes upon compound treatment in FBXO31 KO HEK293T cells and FBXO31 KO cells re-expressing FBXO31 (n = 3 biologically independent samples for degraders; n = 2 biologically independent samples for DMSO). *P* values were calculated using a two-sided t-test and adjusted for multiple comparisons by the Benjamini-Hochberg method.

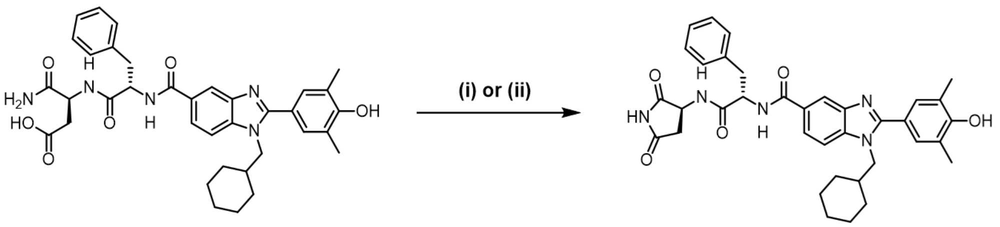

**50** **51**

(i): CDI (3.0 equiv.), DMAP (3.0 equiv.), THF, DMF
(ii): HATU (3.0 equiv.), DMAP (3.0 equiv.), THF, DMF

**Figure S5:** General reaction scheme for the formation of an example CRBN-recruiting cyclic imide. CDI and HATU were compared as carboxylic acid activating reagents for the conversion of C-terminal aspartate **50** to aspartimide **51**.

**50** + CDI + DMAP

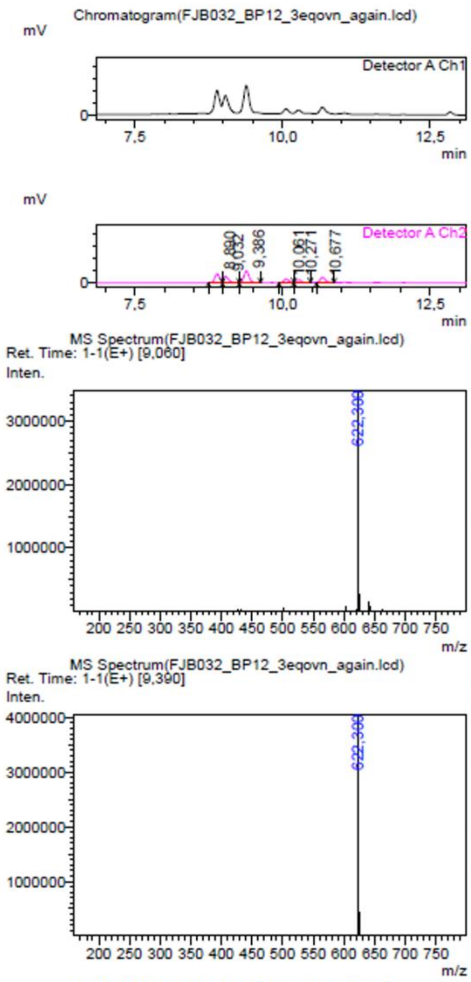

Peak Table(FJB032_BP12_3eqovn_again.lcd)
Detector A Channel 2 254nm

| Peak# | Ret. Time | Area | Height |
|---|---|---|---|
| 1 | 8,890 | 2254041 | 370888 |
| 2 | 9,032 | 1709311 | 273208 |
| 3 | 9,386 | 2898632 | 481496 |
| 4 | 10,061 | 822445 | 147266 |
| 5 | 10,271 | 533789 | 101965 |
| 6 | 10,677 | 1354635 | 216531 |
| Total | | 9572853 | 1591354 |

| Mark | Conc. | Area% |
|---|---|---|
| M | 23,546 | 23,546 |
| M | 17,856 | 17,856 |
| M | 30,280 | 30,280 |
| M | 8,591 | 8,591 |
| M | 5,576 | 5,576 |
| M | 14,151 | 14,151 |
| | 100,000 | 100,000 |

**50** + HATU + DMAP

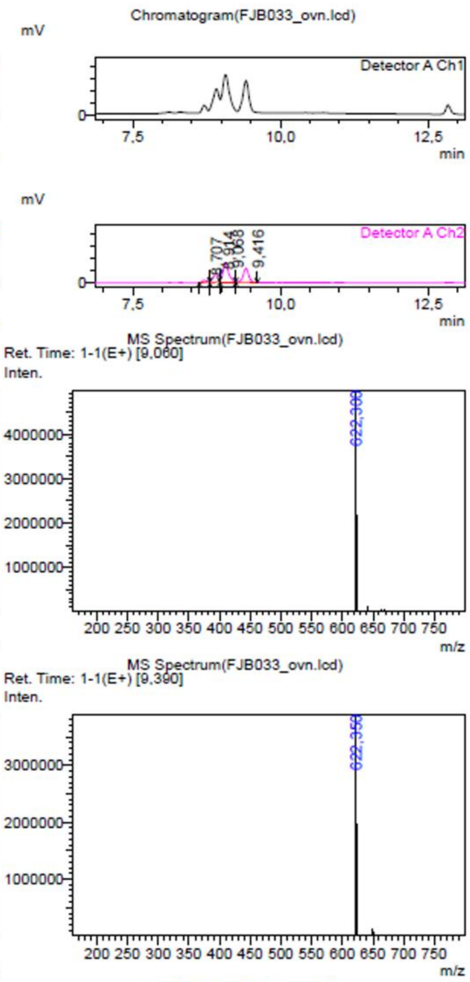

Peak Table(FJB033_ovn.lcd)
Detector A Channel 2 254nm

| Peak# | Ret. Time | Area | Height |
|---|---|---|---|
| 1 | 8,707 | 369418 | 86630 |
| 2 | 8,914 | 2210640 | 336336 |
| 3 | 9,068 | 5173626 | 771742 |
| 4 | 9,416 | 3568954 | 568630 |
| Total | | 11322638 | 1763338 |

| Mark | Conc. | Area% |
|---|---|---|
| M | 3,263 | 3,263 |
| M | 19,524 | 19,524 |
| M | 45,693 | 45,693 |
| M | 31,521 | 31,521 |
| | 100,000 | 100,000 |

**Figure S6:** Crude LC-MS analysis of the example cyclisation reactions described in Figure S4. Major ($t_R$ = 9.06 mins) and minor ($t_R$ = 9.39 mins) products with matching masses were observed to form under the investigated conditions, consistent with aspartimide epimerization. In all cases, the major diastereomer was observed to have the shorter retention time by HPLC-MS. While CDI was used for some preliminary cyclisation attempts, cleaner conversion was observed with HATU, and hence this method was preferred.

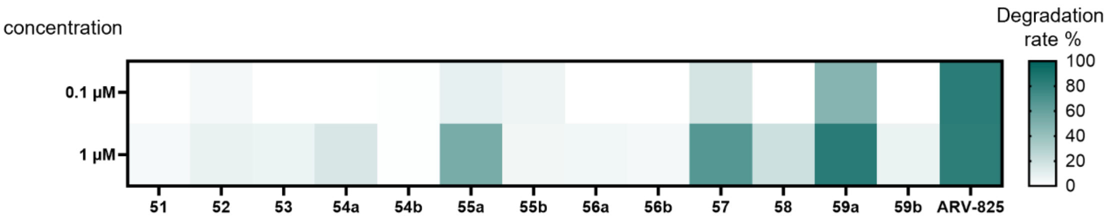

**Figure S7:** Screening for degradation of HiBiT-BRD4 by 1st generation CRBN-PROTAC library. Degradation rate of HiBiT-BRD4 from stable expressing HEK293 cells after treatment with test compounds or ARV-825 at 100 nM or 1 µM for 24h. Data represent mean (n=3 replicates). The major product/first-eluting diastereomer was assigned the suffix 'a', and the minor product/second-eluting diastereomer was assigned suffix 'b'. Examples of 'a' and 'b' for compounds **54**, **55**, **56** and **59** were run in parallel to evaluate relative degradation activity. Thereafter all 'a' compounds were taken forward as the desired *S*-enantiomer products, i.e. **54a** = **54**, **55a** = **55**, etc.

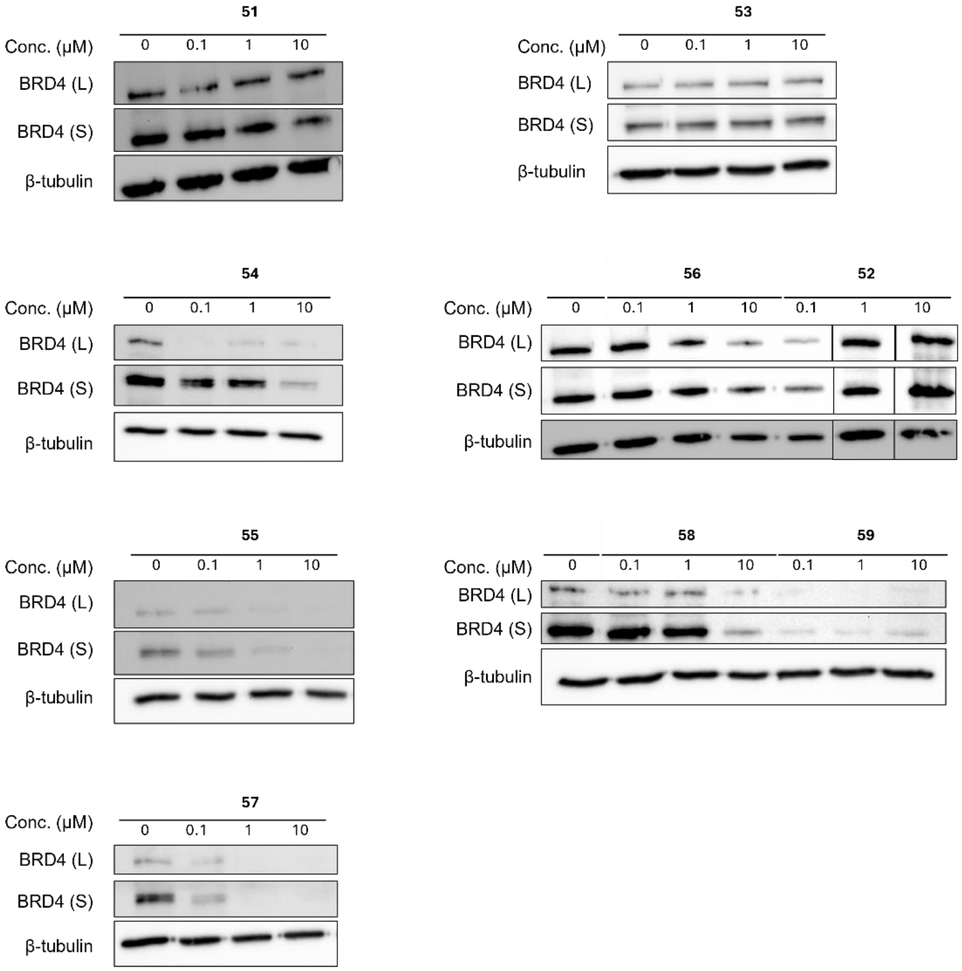

**Figure S8:** Validation of BRD4 degradation for 1st generation CRBN-PROTACs. Western blot analysis of endogenous BRD4 degradation by compounds **51**-**59** in HeLa cells after 24h treatment.

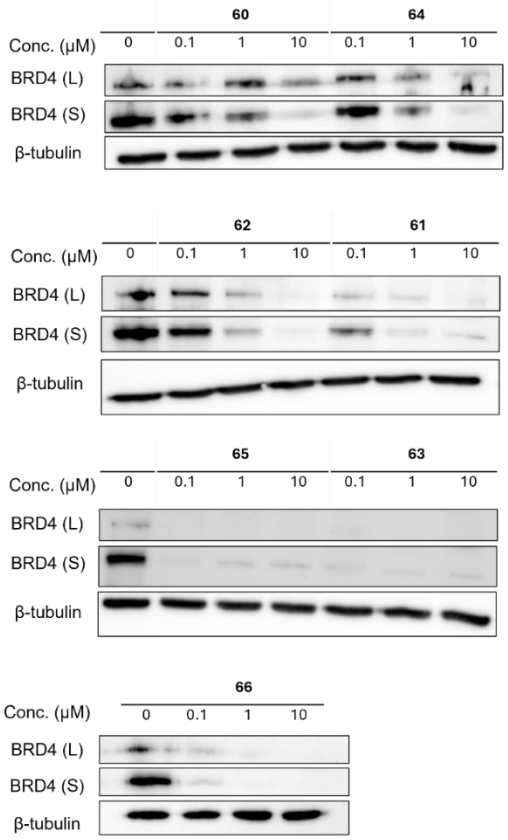

**Figure S9:** Validation of BRD4 degradation for 2nd generation CRBN-PROTACs. Western blot analysis of endogenous BRD4 degradation by compounds **60**-**66** in HeLa cells after 24h treatment.

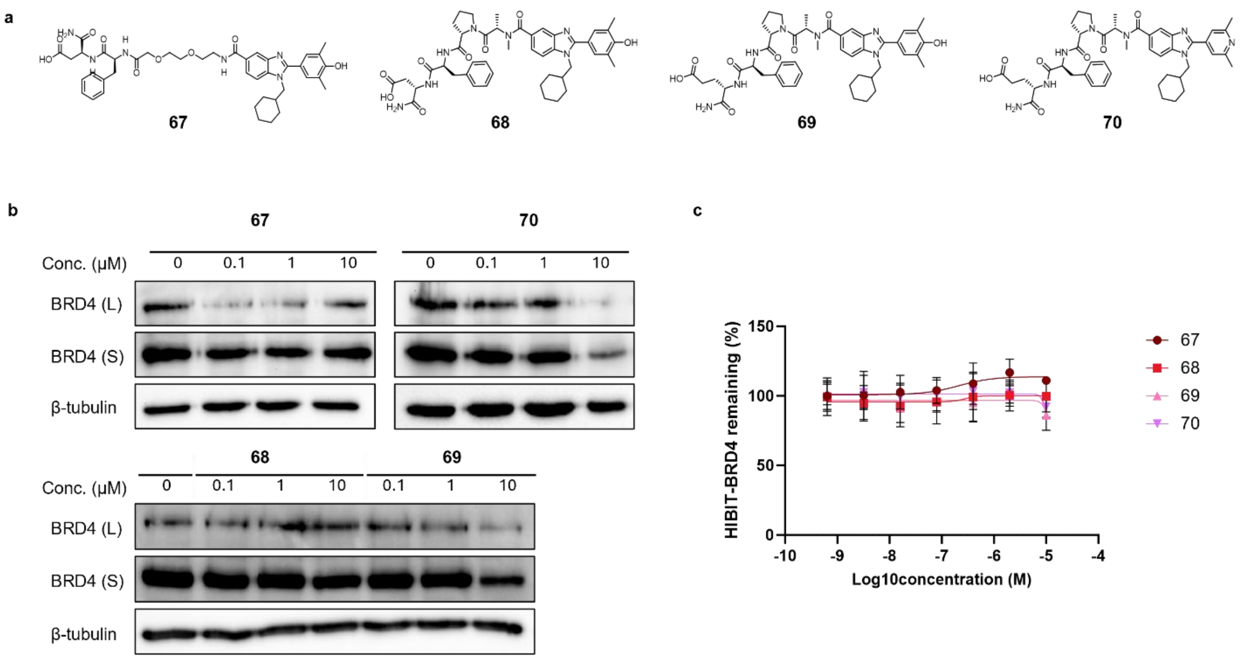

**Figure S10:** Biological evaluation of acyclic CRBN-PROTAC control compounds. a: Structures of exemplary acyclic variants (**67**-**70**) of 1st and 2nd generation CRBN-PROTACs. b: Western blot analysis of endogenous BRD4 degradation by compounds **67**-**70** in HeLa cells after 24h treatment. c: Levels of HiBiT-BRD4 from stable expressing HEK293 cells after treatment with varying concentrations of acyclic compounds **67**-**70** for 24h. Data represent mean ± SD (n=2 separate experiments, with 3 replicates).

### Supplementary Tables

| | General | | Beginning of step | | Mixing/Heating | | End of step | | |
|---|---|---|---|---|---|---|---|---|---|
| | Buffer | Volume (µL) | Release time (s) | Release speed | Mixing time (min) | Temp | Mixing speed | Collect count | Collect time (s) |
| Bead uptake | | 100 | - | - | 5 | r.t. | Bottom mix | 5 | 10 |
| Bead washing (3x) | A | 1000 | 30 | Medium | 3 | r.t. | Medium | 5 | 10 |
| Protein incubation | B | 100 | 30 | Medium | 60 | 10 $^{o}$C | Medium | 5 | 10 |
| Biotin blocking (2x) | C | 1000 | 30 | Medium | 10 | r.t. | Medium | 5 | 10 |
| Bead washing | D | 1000 | 30 | Medium | 3 | r.t. | Medium | 5 | 10 |
| Library Incubation | E | 1000 | 30 | Medium | 60 | 10 $^{o}$C | Medium | 5 | 10 |
| Bead washing (5x) | F | 1000 | 30 | Medium | 0.5 | r.t. | Medium | 5 | 10 |
| Elution (2x) | G | 100 | 30 | Medium | 3 | r.t. | Medium | 5 | 10 |

**Table S1.** The KingFisher program used for affinity selection

| Buffer | CAIX | BRD4 | WDR5 |
|---|---|---|---|
| A | 10% FBS, 1x PBS, 0.02% Tween-20 | 20 mM Tris, 500 mM NaCl, 10% FBS, 0.02% Tween-20 (pH 8.0) | 20 mM Tris, 500 mM NaCl, 10% FBS, 0.02% Tween-20 (pH 8.0) |
| B | CAIX (1.5 µM) in **A** | BRD4 (1.5 µM) in **A** | WDR5 (1.5 µM) in **A** |
| C | 10% FBS, 1x PBS, 0.02% Tween-20, 400 µM d-biotin | 20 mM Tris, 500 mM NaCl, 10% FBS, 0.02% Tween-20, 400 µM d-biotin, (pH 8.0) | 20 mM Tris, 500 mM NaCl, 10% FBS, 0.02% Tween-20, 400 µM d-biotin, (pH 8.0) |
| D | 10% FBS, 1x PBS | 20 mM Tris, 500 mM NaCl, 10% FBS, (pH 8.0) | 20 mM Tris, 500 mM NaCl, 10% FBS, (pH 8.0) |
| E | 100 fmol/member library in buffer **D** | 100 fmol/member library in buffer **D** | 100 fmol/member library in buffer **D** |
| F | 1x PBS | 20 mM Tris, 500 mM NaCl, (pH 8.0) | 20 mM Tris, 500 mM NaCl, (pH 8.0) |
| G | MeCN:MQ (1:1) 0.1%FA | MeCN:MQ (1:1) 0.1%FA | MeCN:MQ (1:1) 0.1%FA |

**Table S1.** Buffer conditions for CAIX, WDR5 and BRD4(DB1)

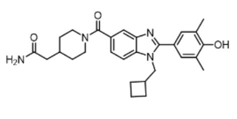

**1**

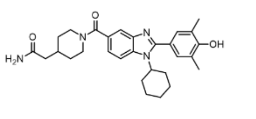

**2**

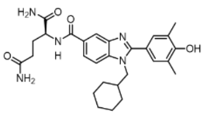

**3**

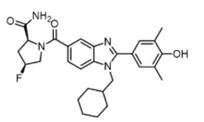

**4**

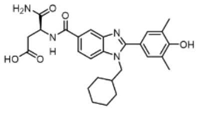

**5**

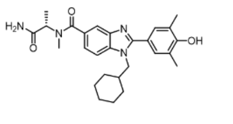

**6**

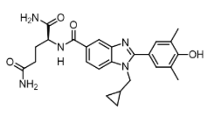

**7**

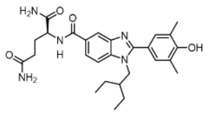

**8**

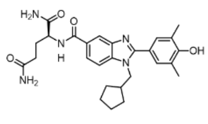

**9**

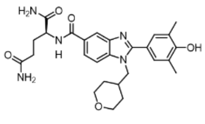

**10**

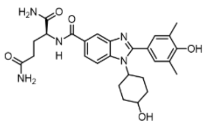

**11**

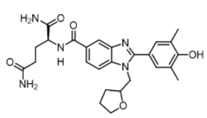

**12**

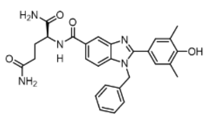

**13**

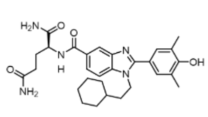

**14**

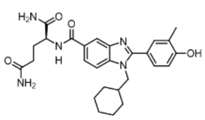

**15**

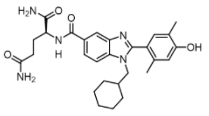

**16**

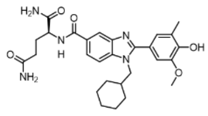

**17**

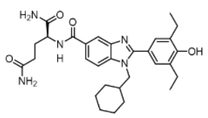

**18**

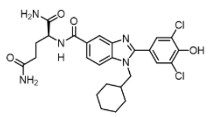

**19**

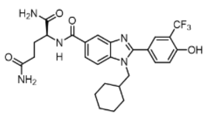

**20**

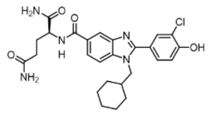

**21**

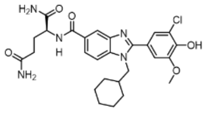

**22**

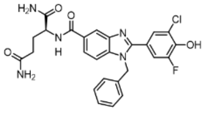

**23**

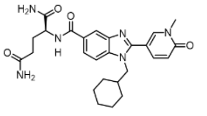

**24**

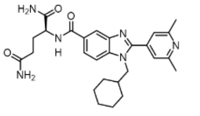

**25**

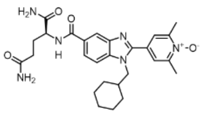

**26**

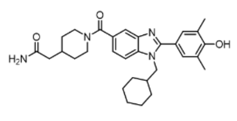

**27**

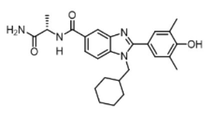

**28**

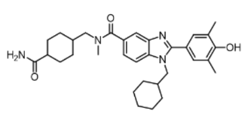

**29**

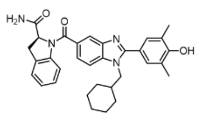

**30**

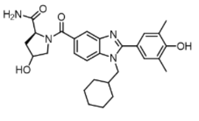

**31**

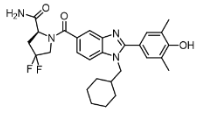

**32**

**Table S3.** Chemical structures of SEL hits **1**-**6** from BRD4 ASMS, and derivatives **7**-**32**.

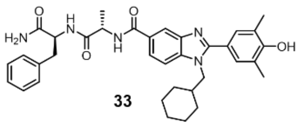

**33**

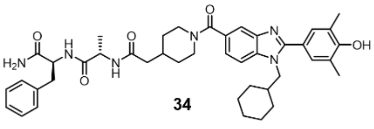

**34**

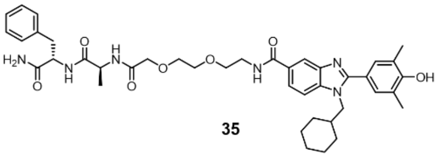

**35**

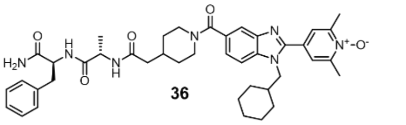

**36**

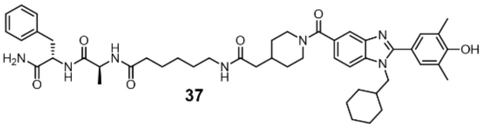

**37**

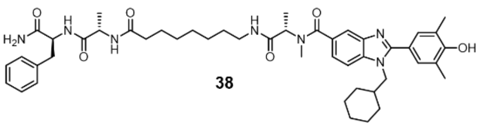

**38**

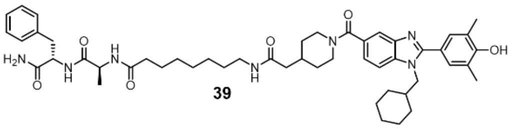

**39**

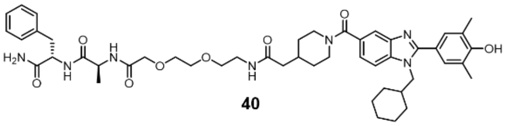

**40**

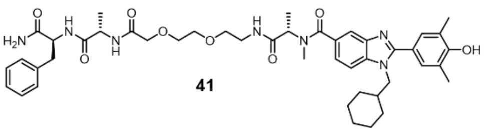

**41**

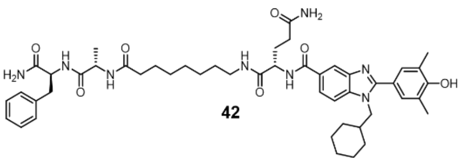

**42**

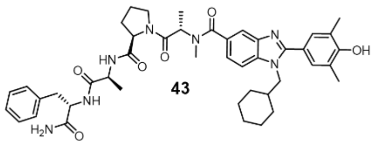

**43**

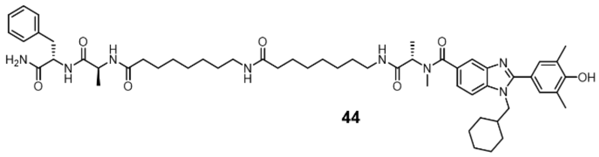

**44**

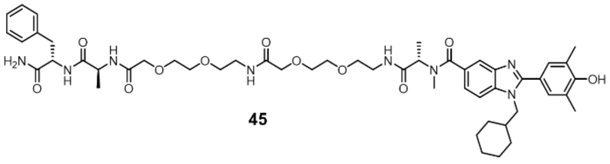

**45**

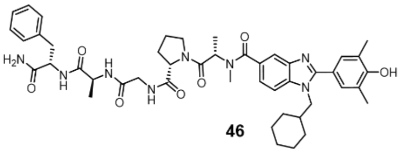

**46**

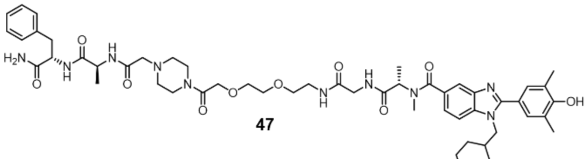

**47**

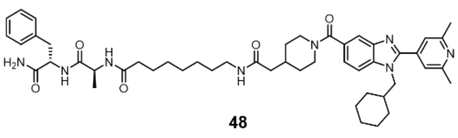

**48**

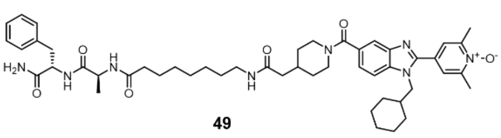

**49**

**Table S4.** Chemical Structures of FBXO31-Recruiting CIPs **33**-**49**

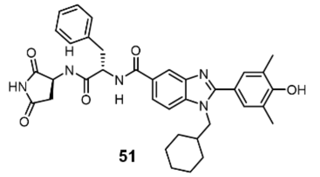

**51**

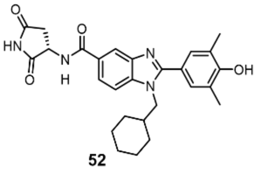

**52**

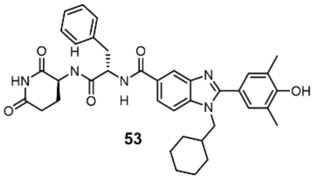

**53**

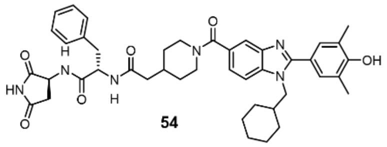

**54**

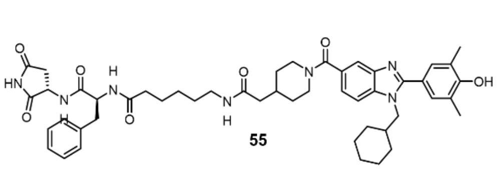

**55**

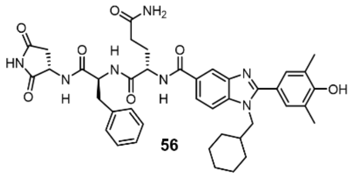

**56**

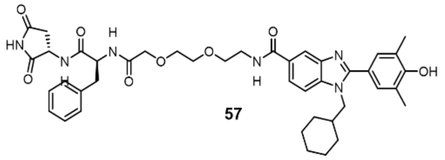

**57**

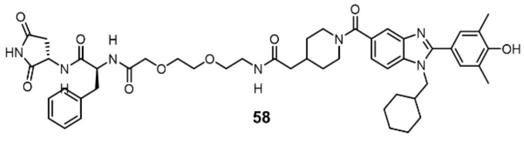

**58**

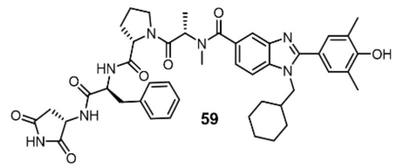

**59**

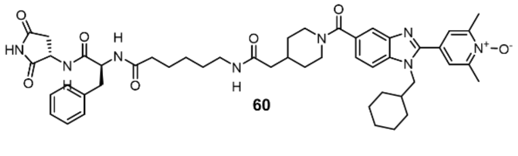

**60**

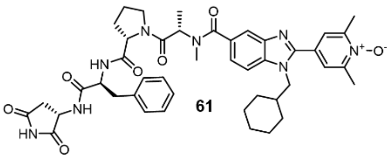

**61**

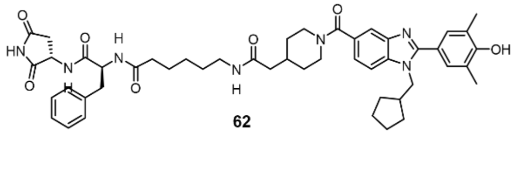

**62**

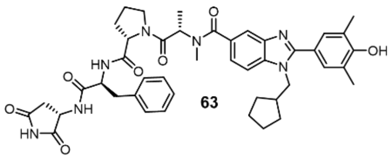

**63**

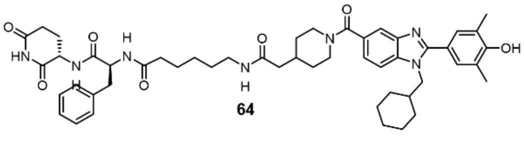

**64**

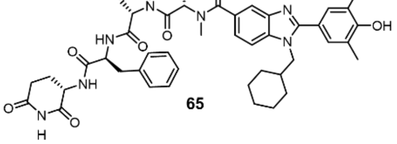

**65**

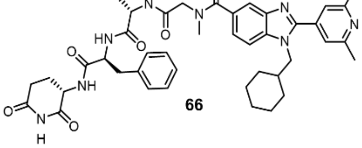

**66**

**Table S5.** Chemical Structures of CRBN-Recruiting CIPs **51**-**66**

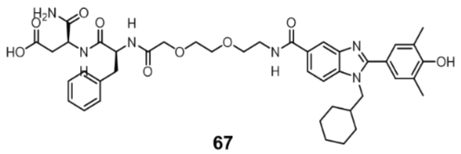

**67**

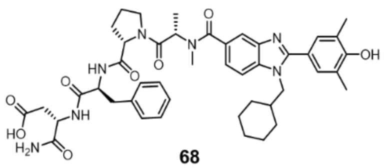

**68**

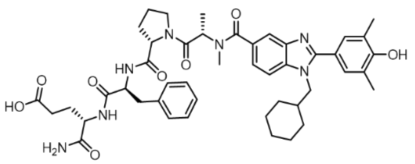

**69**

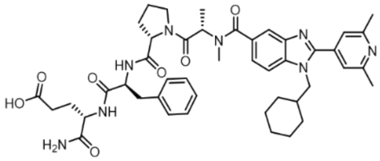

**70**

**Table S6.** Chemical Structures of Acyclic Controls **67**-**70**

**Abbreviations:**

| | |
|---|---|
| ASMS | Affinity-selection mass spectrometry |
| Asp-Phe | Aspartic acid-phenylalanine |
| BB | Building block |
| BLI | Biolayer interferometry |
| BRD2 | BET bromodomain-containing protein 2 |
| BRD4 | BET bromodomain-containing protein 4 |
| CAIX | Carbonic anhydrase IX |
| CDI | Carbonyl diimidazole |
| CIP | Chemical inducer of proximity |
| CRBN | Cereblon |
| D2B | Direct-to-biology |
| $DC_{50}$ | Half-maximal degradation concentration |
| DD-TAC | DNA-damage targeting chimera |
| DEL | DNA-encoded library |
| DMTA | Design-make-test-analyse |
| DUBTAC | Deubiquitinase-targeting chimera |
| FBXO31 | F-box protein 31 |
| FP | Fluorescence polarisation |
| HATU | Hexafluorophosphate azabenzotriazole tetramethyl uronium |
| HEK293 | Immortalized human embryonic kidney cell line |
| HPLC-MS | High-performance liquid chromatography- mass spectrometry |
| $IC_{50}$ | Half-maximal inhibitory concentration |
| KAc | Acetyl-lysine |
| $K_D$ | Equilibrium dissociation constant |
| KO | Knockout |
| LogP | Logarithm of the partition coefficient |
| MW | Molecular weight |
| MQ | Milli-Q Water |
| PEG | Polyethylene glycol |
| Phe-Ala | Phenylalanine-alanine |
| PhosTAC | Phosphorylation-targeting chimera |
| POI | Protein of interest |
| PROTAC | Proteolysis-targeting chimera |

| | |
|---|---|
| RIPTAC | Regulated induced proximity targeting chimera |
| SAR | Structure-activity relationship |
| SEL | Self-encoded library |
| SPS | Solid-phase synthesis |
| TPD | Targeted protein degradation |
| TPSA | Topological polar surface area |
| WDR5 | WD-repeat containing protein 5 |
| WPF | Tryptophan-proline-phenylalanine |

### Experimental

#### Reagents and supplies

**Chemicals**: Reagents and solvents were purchased from Sigma-Aldrich (Merck), Fisher Scientific, BLDpharm, Fluorochem or VWR and were used without further purification unless stated otherwise. Building blocks grouped as Fmoc-protected amino acids, aldehydes and primary amines were purchased from BLDpharm, Tokyo Chemical Industry (TCI), ABCR, and Chemspace. Rink Amide ProTide™ Resin (CEMR002-C) was purchased from CEM. TentaGel® M $NH_2$ 30 µm (M30352) and Polystyrene AM RAM (Rink Amide) 100-200 mesh (H10023) was purchased from Rapp-Polymere.

**Materials for affinity selection**: Dynabeads MyOne Streptavidin T1 was purchased from ThermoFisher Scientific. A KingFisher™ Duo Prime Purification System was used to perform our affinity selection experiments. The protocols were developed with BindIt 4.1 Software.

#### Instrumentation

LC-MS analysis: Compound purity was determined by LC–MS, using the LCMS-2020 system (Shimadzu) with a Gemini 3 µm C18 110 Å column (50 × 3 mm) using the following parameters: flow rate = 0.55 mL/min, scan range = 160-800 m/z, column temperature (°C) = 40. The following gradient of 10–90% MeCN/H2O (0.1% formic acid) over 15 min and measuring UV absorbance at 254 nm was used, unless stated otherwise. Compounds were dissolved in $H_2O$:MeCN:t-BuOH (1:1:1) before injection.

NMR analysis: $^{1}H$ spectra were recorded on a Bruker AV-400 (400 MHz) and Bruker AV 600 MHz spectrometer. $^{13}C$ NMR were recorded on a Bruker AV-400 (400 MHz) spectrometer or Bruker AV-500 (500 MHz) spectrometer. Chemical shift values are reported in parts per million (ppm) and designated by δ. Tetramethylsilane or solvent resonance was used as internal standard. Coupling constants (J) are reported in Hertz (Hz) and multiplicities are indicated by s (singlet), br s (broad singlet), d (doublet), t (triplet), td (triplet of doublets), or m (multiplet).

LC-MS/MS analysis: Analysis was performed on an Vanquish™ Neo UHPLC system (Thermo Scientific) connected to an Orbitrap Exploris 240 mass spectrometer (Thermo Fisher Scientific). Samples were run on a Double nanoViper PepMap Neo column (2 µm particle size, 15 cm x 75 µm, Thermo Fisher Scientific, DNV75150PN) following a PepMap Neo Trap Cartridge (5 µm C18 300 µm X 5 mm). The standard nano-LC method was run without temperature control and a flow rate of 300 nL/min with the following gradient: 0% solvent B ramping linearly to 60% B in A over 78 min, with solvent A = water (0.1% FA), and solvent B = 80% acetonitrile, 20% water (0.1% FA). Positive spray ionization was set at 1900 V. The following parameters were used for MS1 collection: resolution = 120.000, scan range = 275-

650 m/z for SEL 1 and SEL 4, 300-800 m/z for SEL 2 and 275-700 for SEL 3, maximum injection times = 300 ms, RF Lens = 80%, microscans = 1, AGC target = standard, Ion Transfer Tube Temp (°C) = 280, Charge State = +1. For the data dependent MS/MS event the following settings were used: resolution = 15000, number of Dependent Scans= 20, isolation window (m/z) = 1.5, intensity threshold = 1E5, MIPS mode = small molecule, absolute collision energy = 15, 25 eV, AGC Target (%) = 50, microscans = 1, RF Lens(%) = 70, dynamic exclusion = on (exclude after n times = 3, Exclusion duration (s) = 15, excluding isotopes, 10 ppm mass tolerance).

**Software**:

KNIME 5.1.2 software was used for library enumeration, molecular property calculations and the generation of the enriched building block plots in Figure 2. The following extensions were installed: RDkit Nodes version 4.7. and CDK version 1.5.6.

COMET v1.1.0-SNAPSHOT was used for library annotations and can be downloaded via github.com/sirius-ms/comet/releases.

**Protein Preparation**

Biotinylated Human Carbonic Anhydrase IX (38-414), His,Avitag (CA9-H82E3) was purchased from ACROBiosystems.

BRD4(BD1) and WDR5(22–334) were expressed, purified and biotinylated using the following protocol.

**BRD4 Expression and Purification**: Escherichia Coli BL21(DE3) competent cells were collected from the -80 fridge and put on ice. BRD4(BD1) plasmids were collected from the fridge and diluted separately to 45 µg/µL on ice. Plasmids (2 µL) were added to the E.coli BL21(DE3) cells separately and incubated for 20 minutes on ice. The cells were shocked at 42 °C for 60 to 90 seconds and incubated on ice for 5 minutes. PB-medium (1 mL) was added, and the cells were left to incubate at 37 °C, 200 rpm for 1 hour. After incubation the cells were centrifuged for 3 minutes at 3000 rpm, the supernatant was discraded and the pellet resuspended in PB-medium (100 µL). The cells were plated on agar plates and spread evenly, after which the cells were incubated for 16 hours at 37 °C.

Human BRD4(BD1) [135 residues] was transformed into pET-22b(+) vector and overexpressed in Escherichia Coli BL21(DE3) competent cells. A colony of the BRD4(BD1) agar plate was picked and pre-cultured in 100 mL of PB-medium with Ampicillin at 200 rpm, 37 °C for 16 hours. Main culture was performed in four 2L Erlenmeyer flasks, each containing 1 L PB-medium, 500 µL penicillin and 25 mL pre-culture at 200 rpm and at 37 °C. The cells were incubated until an OD-600 between 0.6 and 0.8 was reached, after which the temperature was lowered to 16 °C. At an OD600 of 1.2 1 mL of 0.5 mM IPTG was added to each Erlenmeyer flask and the cells were left shaking at 200 rpm for 16 hours. Cells were harvested using centrifugation. The pellet was resuspended in lysis buffer (50 mM HEPES, 500 mM NaCl, 20 mM imidazole, 5% glycerol [v/v], 0.5 mM PMSF, pH 8.0), and lysed using the sonicator. Lysed cells were centrifuged using the ultra-centrifuge at 4 °C for 1 hour. Nickel column purification on the ÄKTA was performed with running buffer (50 mM HEPES, 500 mM NaCl, 10 mM Imidazole, 0.5 mM TCEP, 1 mM PMSF, pH 8.0) and elution buffer (50 mM HEPES, 500 mM NaCl, 100 mM Imidazole, 0.5 mM TCEP, 1 mM PMSF, pH 8.0). Resuspended BRD4(BD1) was loaded onto the column, after which elution was performed, and fractions were collected with an increasing elution buffer concentration. The purification success was evaluated by SDS PAGE and LC-MS measurement.

**BRD4(BD1) Biotinylation**: To 10 mL BRD4(BD1) (130.9 µM), 650 µL EZ-Link™ NHS-PEG4-Biotin (6.5 µmol dissolved in 1 mL BRD4(BD1) running buffer) was added and the solution was incubated overnight on ice. The biotinylated BRD4(BD1) protein was concentrated, and the protein concentration was determined using a NanoDrop spectrophotometer and stored at -80 °C.

**WDR5 Expression and Purification**: A truncated human WDR5 construct encoding amino acids 22–334(GSSHHHHHHSSGLVPRGSHMMSATQSKPTPVKPNYALKFTLAGHTKAVSSVKFS
PNGEWLASSSADKLIKIWGAYDGKFEKTISGHKLGISDVAWSSDSNLLVSASDDKTLKIWDVSSGK
CLKTLKGHSNYVFCCNFNPQSNLIVSGSFDESVRIWDVKTGKCLKTLPAHSDPVSAVHFNRDGSLI
VSSSYDGLCRIWDTASGQCLKTLIDDDNPPVSFVKFSPNGKYILAATLDNTLKLWDYSKGKCLKTYT
GHKNEKYCIFANFSVTGGKWIVSGSEDNLVYIWNLQTKEIVQKLQGHTDVVISTACHPTENIIASAAL
ENDKTIKLWKSDCAE) was synthesized by GenScript and subcloned into a pET-15b expression vector containing an N-terminal 6×His-SUMO tag using the NdeI and BlpI restriction sites. The WDR5 plasmid was then transformed into E. coli BL21 (DE3) cells. The cells were cultured in Luria–Bertani medium at 37 °C. When the optical density at 600 nm (OD600) reached 0.8, the temperature was lowered to 25 °C. Protein expression was induced by adding 1 mM isopropyl-β-D-thiogalactoside (IPTG), and the incubation continued for 16 hours at this temperature. The cells were harvested by centrifugation, resuspended in lysis buffer (20 mM Tris-HCl, 500 mM NaCl, pH 8.0), and then lysed using a homogenizer (FPG12800). The lysate was cleared by centrifugation, and the supernatant was collected. The protein was then bound to a nickel affinity column (HisTrap™, Cytiva) using an ÄKTA system. Protein elution was carried out applying an imidazole gradient. The purified protein was analyzed by SDS PAGE and concentrated using a 10K molecular weight cut-off centrifugal concentrator. The concentration was determined using a NanoDrop spectrophotometer.

**WDR5 Biotinylation**: WDR5 was biotinylated using the EZ-Link™ NHS-PEG4-Biotin reagent (Promega) according to the manufacturer's protocol. Briefly, the purified WDR5 protein was diluted to a concentration of approximately 1 mg/mL. The NHS-PEG4-biotin reagent was then added to the protein solution to a final concentration of 10–20 µM. The reaction was incubated for 1 hour at room temperature with gentle agitation to facilitate the biotinylation process. After the incubation, unreacted biotin reagent was removed using a 10K molecular weight cut-off centrifugal concentrator. The biotinylated WDR5 protein was stored at −80 °C for future use. The degree of biotinylation was confirmed using BioLayer Interferometry to verify biotin incorporation.

**Fluorescence Polarization Assays**

**Direct fluorescent polarization (FP) binding assay to BRD4**

The $K_D$ of FITC labeled JQ1 was determined by fluorescence polarization assay. Measurements were performed in buffer containing 20 mM Tris-HCl, 500 mM NaCl, pH 8.0, using 20 nM FITC-JQ1 as the fluorescent tracer and increasing concentrations of BRD4. Assays were carried out in black 384-well microplates (Corning®, CLS3573) in a total volume of 60 µL per well by mixing 30 µL 2x protein solution with 30 µL 2x tracer solution. The mixtures were gently mixed, briefly centrifuged, and incubated for 30 min at room temperature before measurement. Fluorescence polarization was recorded on a FlexStation3 plate reader (Molecular Devices) with an excitation wavelength of 490 nm and an emission wavelength of 530 nm.

The $K_D$ value was determined by fitting the experimental data according a model described by Wang *et.al.*[1] using the software Graphpad Prism 10 (Dotmatics, Boston, MA). Reported values are the mean ± SD of at least two separate experiments in triplicates.

Determined $K_D$ value: 358 ± 11 nM

**Competitive Fluorescent polarization (FP) assay to BRD4**

Binding affinities of resynthesized selection hits and iteratively synthesized derivatives towards BRD4 were determined by competitive fluorescence polarization assay using FITC-JQ1 as fluorescencent tracer. Measurements were performed in buffer containing 20 mM Tris-HCl, 500 mM NaCl, pH 8.0, using 20 nM FITC-JQ1, 500 nM BRD4 and increasing concentrations of test compounds. Assays were carried out in black 384-well microplates (Corning®, CLS3573) in a total volume of 60 µL per well by mixing 30 µL 2x protein solution with 15 µL 4x tracer solution and 15 µL 4x test compound solution. The mixtures were gently mixed, briefly centrifuged, and incubated for 30 min at room temperature before measurement. Fluorescence polarization was recorded on a FlexStation3 plate reader

(Molecular Devices) with an excitation wavelength of 490 nm and an emission wavelength of 530 nm.

The $K_i$ values were determined by fitting experimental binding curves according a model described by Wang et al.[1] using the software Graphpad Prism 10 (Dotmatics, Boston, MA). Reported values are the mean ± SD of at least two separate experiments in triplicates.

**Cell Lines and Cell Culture**

The human cell lines HeLa and HEK293T were purchased from American Type Culture Collection (ATCC, Manassas, VA, USA). The HEK293 HiBiT BRD4 cell line was a gift from the Ciulli lab.

Generation and culturing of of FBXO31 KO HEK293T cells was as previously described.[2]

Hela was cultured in the MEM cell culture medium supplemented with 10% fetal bovine, 100 IU /ml penicillin, 100 µg/mL streptomycin and 2 mM GlutaMAX. HEK293T and HEK293 HiBiT-BRD4 were cultured in the DMEM cell culture medium supplemented with 10% fetal bovine, 100 IU /ml penicillin, 100 µg/mL streptomycin and 2 mM GlutaMAX.

All cell lines were cultured in a humidified 5% CO2 incubator at 37 °C. All cell lines were monitored for mycoplasma contamination.

**Cell Viability Assay**

Cytotoxicity was assessed using the CellTiter-Glo luminescent cell viability assay (Promega, cat. #G7573). Briefly, HEK293T FBXO31-KO cells and FBXO31-KO cells re-expressing FBXO31 were plated in white, clear-bottom 96-well plates (Corning, cat. #3610) at 3,000 cells per well in 100 µL of DMEM and treated with serially diluted compounds for 72 h, maintaining a final DMSO concentration of 0.5% (v/v) in a total volume of 100 µL. At the end of the treatment period, CellTiter-Glo reagent (50 µL per well) was dispensed, and plates were equilibrated at room temperature for 10 min before luminescence was recorded on a CLARIOstar Plus plate reader (BMG LABTECH). Viability was normalized to DMSO-treated cells of the corresponding cell line.

**HiBiT assay**

Degradation of HiBiT-BRD4 by test compounds was determined by NanoGlo® HiBiT Lytic detection system (Promega). HEK293 cells stably expressing HiBiT-BRD4 were seeded in a white opaque 96-well culture plate (Costar REF 3917) at 2 x$10^4$ cells/well in 80 µL Opti-MEM with 4 % FBS. After incubation for 24h at 37 °C with 5% CO2 cells were treated with 20 µL of varying concentrations of 5x test compound or DMSO in Opti-MEM with 4 % FBS (final DMSO concentrations of 0.1%) for 24h. Cells were equilibrated at rt for 15 min before removal of 50 µL of medium and lysis by addition of 50 µL lytic buffer supplemented with LgBiT protein (1:100) and Nano-Glo substrate (1:50). After incubation in the dark for 10 min on an orbital shaker (250 rpm) luminescence was measured using an EnVision multilabel plate reader (PerkinElmer). Empty HEK293 cells were used for background luminescence subtraction. Dose-response curves were analyzed using the Graphpad Prism 10 software (Dotmatics, Boston, MA). $DC_{50}$ values were obtained by nonlinear regression analysis. Reported values are the mean ± SD of at least two separate experiments in triplicates.

**Western Blot**

**Western blot analysis for FBXO-PROTACs**

Endogenous BRD4 degradation by test compounds 33-49 in FBXO31 KO HEK293T or FBXO31 KO HEK293T cells reexpressing FBXO31 was analyzed as previously described.[2]

**Western blot analysis for CRBN-PROTACs**

Endogenous BRD4 degradation by test compounds (51-70) was validated by western blot analysis in HeLa cells. Cells were seeded in a 6-well plate at 2.8 x10^5 cells per well in MEM supplemented with 10% FBS and 100 U ml−1 penicillin–streptomycin and incubated for 24h at 37 °C with 5% CO2. The old culture medium was replaced and cells were treated with different concentrations of test compounds or DMSO diluted in culture medium with a final DMSO concentration of 0.1%. After treatment for 22h cells were washed twice with ice-cold PBS and the lysed on ice adding 100 µL/well radioimmunoprecipitation assay (RIPA) buffer (Thermofisher) supplemented with 1 mM phenylmethylsulfonyl fluoride (PMSF), 1 mM NaF and Roche cOmplete™ Protease Inhibitor Cocktail. Lysates were centrifuged at 20,000g for 30 min at 4 °C and total protein concentrations were subsequently determined using Thermo Scientific™ Pierce™ BCA Protein Assay. 20 µg of each protein lysate were loaded onto an 8% SDS-PAGE gel and separated by gel electrophoresis. Protein transfer to PVDF membranes (BioRad) was performed using the Trans-Blot Turbo Transfer System (Bio-Rad). Membranes were cut to separately detect BRD4 and beta-Tubulin as normalization control and blocked by incubation in 0.5 % milk in TBST for 1h at rt. Blots were then incubated with primary antibodies (anti-BRD4 rabbit monoclonal antibody, Abcam, ab128874; 1:1,000 or anti-beta-Tubulin mouse monoclonal antibody, Invitrogen, BT7R, 1:2000) in 0.5 % milk at 4 °C overnight. After washing with TBST and 2x TBS membranes were incubated with respective secondary antibodies in 0.5 % milk in TBST (1. polyclonal goat anti-rabbit HRP-coupled antibody, Jackson ImmunoResearch Laboratories, 111-035-003, 1:5000; 2. polyclonal goat anti-mouse HRP-coupled antibody, Jackson ImmunoResearch Laboratories, 115-035-003, 1:5000) for 1h at rt. After additional washing membranes were visualized on a ChemiDoc MP imaging system (Bio-Rad) after treatment with the Thermo Scientific™ Pierce™ ECL Western Blotting Substrate.

**Global Proteomics**

Cell pellets were lysed by sonication (three cycles of five pulses at 40% amplitude) in 100 µL of PBS containing cOmplete protease inhibitor cocktail, and protein concentrations were determined by the DC protein assay. For each sample, 100 µg of protein in 50 µL PBS was denatured by mixing with an equal volume of 12 M urea in PBS. Disulfide reduction was carried out with 5 µL of 200 mM DTT (65 °C, 15 min), and free cysteines were alkylated with 5 µL of 400 mM iodoacetamide in water (37 °C, 30 min, protected from light). After dilution with 300 µL of PBS, samples were digested with 2 µg of trypsin/LysC for 12 h at 37 °C. For isobaric labeling, 8.5 µg of peptides was reconstituted in 35 µL PBS containing 9 µL of acetonitrile and incubated with TMT reagent (5 µL per channel, 20 µg/µL in dry acetonitrile) for 1 h at room temperature. Reactions were quenched by adding 6 µL of 5% hydroxylamine followed by 2.5 µL of formic acid. The labeled samples were pooled, separated into ten fractions with a high-pH reversed-phase peptide fractionation kit (Thermo Scientific, cat. #84868), dried under vacuum centrifugation, and subjected to LC-MS analysis on an Orbitrap Eclipse Tribrid mass spectrometer interfaced with a Vanquish Neo UHPLC system.

Peptides were separated on an EASY-Spray C18 analytical column (2 µm particle size, 75 µm i.d. × 150 mm) at a flow rate of 0.25 µL/min using a multistep gradient of mobile phase A (0.1% formic acid in water) and mobile phase B (0.1% formic acid in 80% acetonitrile): 5% B for 0–15 min, a linear increase from 5% to 45% B over 15–155 min, and a final ramp from 45% to 100% B between 155 and 180 min. The nanoelectrospray voltage was 1.5 kV. Each acquisition cycle started with a full MS scan in the Orbitrap (resolution, 60,000; scan range, m/z 375–1600; RF lens, 60%; standard AGC target; automatic maximum injection time). Precursors selected for MS2 were isolated in the quadrupole with a 0.7 m/z window and fragmented by HCD (normalized collision energy, 27%), with fragment ions detected in the ion trap (standard AGC target; maximum injection time, 35 ms). For each MS2 spectrum, up to ten fragment ions were co-isolated by synchronous precursor selection (SPS) and subjected to MS3 analysis, in which ions were fragmented by HCD (collision energy, 55%) and analyzed in the Orbitrap (AGC target, 250%; maximum injection time, 200 ms; resolution, 60,000). Spectra were recorded with Xcalibur software (v4.5.445.18, Thermo Scientific).

Raw files were processed in Proteome Discoverer (v2.5, Thermo Scientific), and peptides were identified with the Sequest HT search engine against the UniProt human reference proteome (UP000005640_9606_Human.fasta). Searches were performed with full tryptic specificity, a precursor mass tolerance of 10 ppm, and a fragment mass tolerance of 0.6 Da. Cysteine carbamidomethylation and TMT modification of peptide N-termini and lysine side chains were set as fixed modifications; methionine oxidation and protein N-terminal acetylation were allowed as variable modifications. Peptide-spectrum matches were scored with Percolator, and results were filtered to a 1% false discovery rate at both the peptide and protein levels. TMT reporter ion intensities were extracted from SPS-MS3 scans with a 20 ppm integration window, and protein abundances were computed from unique and razor peptides.

**Library Enumeration:**

A benzimidazole library was enumerated from a curated set of 53 amino acids, 52 primary amines and 64 aldehydes in a KNIME workflow using reaction SMARTS in RDKit.

| | Reaction SMARTS |
|---|---|
| C-terminus amidation | [#8;A;X1H0-,X2H1:4][#6;A;X3:2]=[O:1]>>[#7;A;X3:3][#6;X3:2]=[O;X1:1] |
| Amide Coupling | [#7;A;X3;H2,H1;!$(NC=O);!$(NC=CC=O);!$(NC=S);!$(NC=N)!$(N-S):3].[#8;A;X1H0-,X2H1][#6;A;X3:2]=[O:1]>>[#7;A;X3:3][#6;X3:2]=[O;X1:1] |
| $S_NAr$ | [#7:1]-[#6:2](=[O:3])-[c:4]1[c:5][c:6][c:7](F)[c:8]([c:10]1)-[#7+](-[#8-])=O.[#7;A;H2X3;!$(NC=[!#6]);!$(N-S):15]>>[#7:1]-[#6:2](=[O:3])-[c:4]1[c:5][c:6][c:7](-[#7:15])[c:8]([c:10]1)-[#7+](-[#8-])=O |
| Nitro Reduction | [#7:9]-[#6:7](=[O:10])-[c:5]1[c:4][c:3][c:2][c:1]([c:6]1)-[#7+](-[#8-])=O>>[#7:9]-[#6:7](=[O:10])-[c:5]1[c:4][c:3][c:2][c:1](-[#7:8])[c:6]1 |
| Heterocyclisation | [#7:11]-[#6:9](=[O:10])-[c:6]1[c:5][c:4][c:2](-[#7:18])[c:3](-[#7:8])[c:7]1.[#6:13][#6;A;H1X3:12]=[O:15]>>[#6:13]-[c:12]1[n:18][c:2]2[c:4][c:5][c:6]([c:7][c:3]2[n:8]1)-[#6:9](-[#7:11])=[O:10] |
| | Deprotection SMARTS |
| Fmoc | [#7:17]-[#6:16](=[O:18])-[#8:15]-[#6:14]-[#6:1]-1-[c:5]2[c:9][c:8][c:7][c:6][c:4]2-[c:3]2[c:10][c:11][c:12][c:13][c:2]-12>>[#7:17] |
| Trityl (amine) | [N:13][C:14]([c:12]1[c:1][c:2][c:3][c:4][c:5]1)([c:11]1[c:6][c:7][c:8][c:9][c:10]1)[c:15]1[c:16][c:17][c:18][c:19][c:20]1>>[N;A;X3,H2:13] |
| Trityl (aromatic amine) | [n:13][C:14]([c:12]1[c:1][c:2][c:3][c:4][c:5]1)([c:11]1[c:6][c:7][c:8][c:9][c:10]1)[c:15]1[c:16][c:17][c:18][c:19][c:20]1>>[n;A;X3,H1:13] |
| Boc | [N:3]-[#6:4](=[O:10])-[#8:5][C:6]([#6:7])([#6:8])[#6:9]>>[N;A;X3;H2:3] |
| Boc (aromatic) | [n:3]-[#6:4](=[O:10])-[#8:5][C:6]([#6:7])([#6:8])[#6:9]>>[n;A;X3;H1:3] |
| tBu | [#6;H3:5][C:2]([#6;H3:4])([#6;H3:3])[#8:1]>>[#8:1] |
| Trityl (N,O,S) | [#7,#8,#16;A:13][C:14]([c:12]1[c:1][c:2][c:3][c:4][c:5]1)([c:11]1[c:6][c:7][c:8][c:9][c:10]1)[c:15]1[c:16][c:17][c:18][c:19][c:20]1>>[#7,#8,#16;A:13] |
| Pbf | [#6:11]-[#8:10]-[c:9]1[c:8][c:6](-[#6:7])[c:5]([c:14](-[#6:15])[c:12]1-[#6:13])[S:2]([#7:1])(=[O:3])=[O:4]>>[#7:1] |

### Library Synthesis

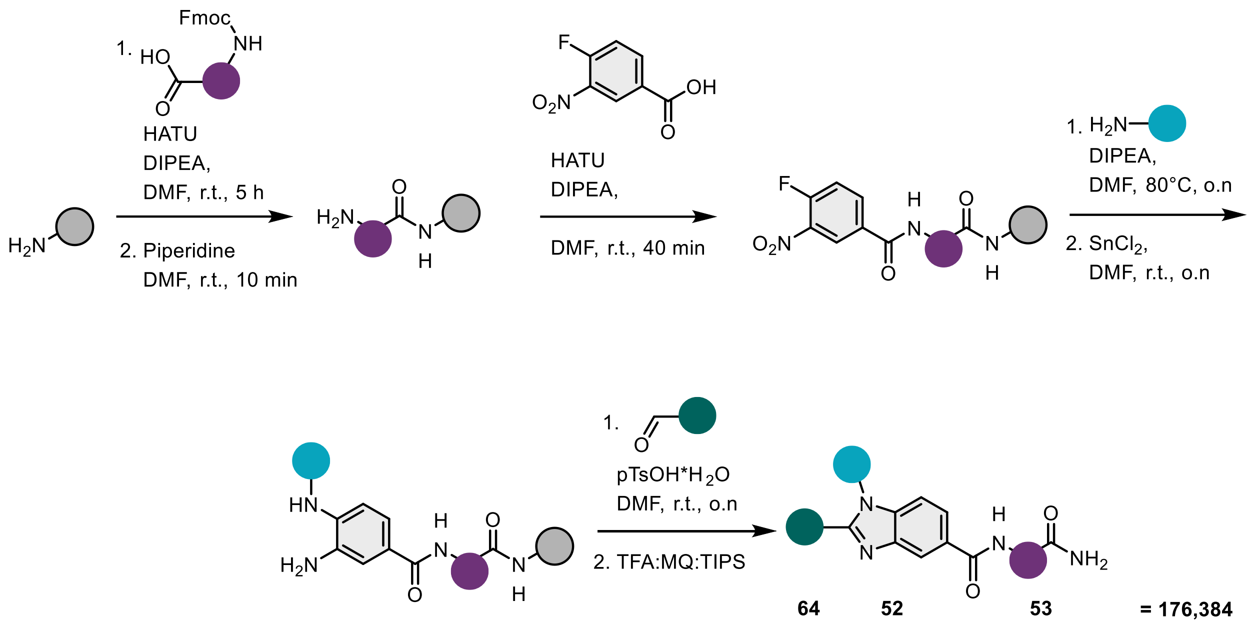

A 176,384-member benzimidazole library was synthesised via split-and-pool combinatorial chemistry. TentaGel S $NH_2$ resin (30 µm, 0.27 mmol/g loading, 982 mg, 265 µmol, 1.0 eq.) functionalized with Fmoc-Rink Amide linker was divided over 53 fritted syringes. A solution of each Fmoc-protected amino-acid (15 µmol, 3.0 eq), HATU (14.90 µmol, 0.2 M, 2.98 eq) and DIPEA (45 µmol, 9.0 eq) in DMF was added to the resin and reacted for 5 hours. The resin was pooled and washed with DMF (5 x 2 mL) and 20% piperidine in DMF (1x, 2 mL) before incubating with 20% piperidine in DMF for 10 min. The resin was washed with DMF (5 x 2 mL) and a solution of 4-fluoro-3-nitrobenzoic acid (147 mg, 795 µmol, 3.0 eq), HATU (1.974 mL, 789 µmol, 2.98 eq) and DIPEA (415 µL, 2.385 mmol, 9.0 eq) in DMF was added to the resin. After 40 min of reaction the resin was washed with DMF (5 x 2 mL).

The resin was divided over 52, eppendorf tubes, to which amines (50 µmol, 10 eq.) and DIPEA (8.71 µL, 50.00 µmol, 10 eq.) in DMF (110 µL) were added. The mixture was shaken at 600 rpm overnight at 80 °C (16:00-9:00). The resin was pooled and washed with DMF (5 x 2 mL) and DCM (5 x 2 mL). A solution of 1.0 M $SnCl_2$ (3.73 g, 16.5 mmol, 62.5 eq.) in DMF was added to the resin (265 µmol). The mixture was incubated overnight at r.t., whereafter the resin was washed with DMF:$H_2O$ (1:1, 5 x 2 mL), with DMF (5x, 2 mL) and DCM (5x, 2 mL).

The resin (4.14 µmol, 1.0 eq.) was split over 64 eppendorf tubes to which the appropriate aldehyde (20.7 µmol, 5 eq.) and p-TsOH•$H_2O$ (3.94 mg, 20.7µmol, 5 eq.) and DMF (120 µL) were added. The mixture was incubated overnight at 600 rpm at r.t. After incubation, the resin was washed with DMF (5 x 2 mL) and DCM (5 x 2 mL). The resin was incubated for 1.5 hours with a solution of TFA:DCM:TIPS (50:50:5) and washed once with a solution of TFA:DCM:TIPS

(50:50:5). TFA was evaporated under a stream of $N_2$ and the library was purified using reverse phase column chromatography with a stepwise gradient of 0-70-100% MeCN:$H_2O$ (0.1% TFA).

**Affinity Selection Procedure**

*Affinity Selection*: The affinity selection experiments were executed using a KingFisher™ Duo Prime Purification System, as previously reported.[3] All operational protocols were programmed and managed via BindIt 4.1 Software. Selection assays targeting BRD4, CAIX and WDR5 were conducted in triplicate. The selections against CAIX and WDR5 were used as negative controls.

For each run, 150 pmol of biotinylated protein was immobilized onto 1 mg of Dynabeads MyOne Streptavidin T1. This immobilized target was then exposed to the library, which was applied at a concentration of 100 fmol per member. Unless otherwise specified, these procedures followed the standardized KingFisher protocol detailed in **Table S1**-**2**.

*Sample preparation after Affinity Selection*: Samples from the affinity selection procedure were lyophilized and resuspended in 50 µL MQ 0.1%FA. The StageTips were prepared as described by Rappsilber *et al*.[4] using C18 material from Empore SPE 47 mm discs (66883-U, Merck). The StageTips were pre-conditioned with 200 µL MeOH, 200 µL of 0.1% (v/v) FA in MeCN and 200 µL of 0.1% (v/v) FA in MQ, respectively by centrifuging for 3 min at 300 rpm. The samples were then loaded on the StageTips and washed with 200 µL of 0.1% (v/v) FA in MQ. Compounds were eluted by adding 200 µL of 0.1% (v/v) FA in MeCN:MQ (7:3). The samples were lyophilized before resuspending in 25 µL 0.1% (v/v) FA in UPLC-MS grade water. The samples were centrifuged for 5 min at 13.000 rpm. Afterwards, 23 µL was transferred to an LC-MS vial and 20 µL was injected into the LC-MS/MS system.

**Data processing**:

Raw files (.raw) from the Orbitrap Exploris 240 mass spectrometer were converted to .mzML files using MSConvert selecting Peak Picking>MS levels 1-2. The resulting .mzML files were processed by mzmine.

MZmine 4.5.0 was used to perform a background subtraction using a three-fold intensity threshold with samples measured against streptavidin-coated magnetic beads. The mzwizard feature was used with the following settings:

> HPLC - Smoothing= yes. Stable ionization across samples= yes. Crop retention time= 0.00-78.00 min. Max peaks in chromatography= 15. Minimum consecutive scans= 4. Approximate feature FWMN= 0.10 min. RT tolerance (intra-sample)= 0.08 min. RT tolerance (sample-to-sample)= 5 min.

Orbitrap - Ion mode= positive with absolute intensity. Noise threshold: MS1= 1.0E5, MS2= 0. Minimum feature height= 3.0E5. m/z tolerance (scan-to-scan)= 0.0010 m/z or 5.0 ppm. m/z tolerance (intra-sample)= 0.0015 m/z or 5.0 ppm. m/z tolerance (sample-to-sample)= 0.0015 m/z or 5.0 ppm.
Filters - Original feature list= remove. Min samples per aligned feature= max of 1 sample or 0.0%. Only keep features with 13C= no.
Annotation - Local compound database search= no. Annotate lipids= no.
The generated feature list was used in the Feature list blank subtraction using the following parameters: Minimum # detection in blanks= 1. Quantification= height. Ratio type= maximum. Fold charge increase= 300%.
The Feature list row filter was used on the generated feature list to remove features that are only enriched in 1 out of the 3 selection samples.
Minimum aligned features (samples) = 2
Never remove features with MS2 = no

The subtracted feature list was subsequently exported to SIRIUS and SIRIUS:COMET spectra annotation was performed using the following settings:

Building blocks = “ENL413.csv”
Scaffold formula = C8H3N2O
MS1 mass accuracy (ppm) = 5
Enable peak matching filter = on
Considered fragment types = S[1;2],S[0;2],S[2], 0, 1,
Minimum number of matching peaks = 1
Number of considered peaks = 5
Number of allowed hydrogen shifts = 1
MS2 mass accuracy (ppm) = 5

The remaining scans were selected and computed with the following settings

Instrument = Orbitrap
MS2 mass accuracy (ppm) = 5 ppm
Fix formula for detected lipid = on
Fallback adducts = [M+H]+
Molecular formula generation = Database search
ZODIAC = off
Predict properties CSI:FingerID = on
Score threshold = on
CSI:FingerID = Rank with EPIMETHEUS

**Automated Parallel Synthesis**

Automated synthesis steps were performed using a Biotage Syro I according to a set of synthetic sequences previously described.[5] Automated methods followed a sequence of procedures, here referred to by single letter codes:

Resin Swelling and Fmoc deprotection (Preliminary setup)*:*

In a fritted syringe, Fmoc-protected polystyrene AM RAM resin (60 µmoles) was treated with DMF (2 mL) and incubated at ambient temperature for 5 min. Syringe was thereafter drained under reduced pressure. To resin was added 20% piperidine in DMF solution (1.5 mL) and agitated at 70 °C for 10 mins. The resin was drained and thereafter subject to washes with DMF (3 x 1.2 mL).

[A] Amide coupling:

To resin was added a 0.4M solution of Fmoc-protected amino acid in DMF (0.75 mL, 0.3 mmoles, 5.0 equiv.), followed by treatment with 0.36M HATU in DMF (0.75 mL, 0.27 mmol, 4.5 equiv.) and DIPEA (85 µL, 0.48 mmol, 8.0 equiv.). The reaction was agitated at 70 °C for 40 mins, and thereafter resin was subject to washes with DMF (3 x 1.2 mL).

[B] Fmoc Deprotection:

To resin was added 20% piperidine in DMF solution (1.5 mL) and agitated at 70 °C for 4 mins. The resin was drained and thereafter subject to washes with DMF (5 x 1.2 mL).

[C] Amide Coupling:

To resin was added a 0.4M solution of 4-fluoro-3-nitrobenzoic acid in DMF (0.45 mL, 0.18 mmoles, 3.0 equiv.), followed by treatment with 0.36M HATU in DMF (0.45 mL, 0.16 mmol, 2.7 equiv.) and DIPEA (85 µL, 0.48 mmol, 8.0 equiv.). The reaction was agitated at 70 °C for 40 mins, and thereafter resin was subject to washes with DMF (3 x 1.2 mL).

[D] Nucleophilic Aromatic Substitution:

Resin was washed with DMSO (3 x 1.5 mL), and thereafter treated with a 0.4M solution of primary amine in DMSO (1.5 mL, 0.6 mmoles, 10.0 equiv.) and DIPEA (105 µL, 0.6 mmol, 10.0 equiv.). The reaction was agitated at 80 °C for 8 h, and thereafter resin was subject to washes with DMF (3 x 1.2 mL).

[E] Nitro reduction:

To resin was added a 1M solution of tin (II) chloride dihydrate in DMF (2.4 mL, 2.4 mmol, 40 equiv.). The reaction was agitated at ambient temperature for 16 h, and thereafter resin was subjected to washes with 50% MilliQ $H_2O$ in DMF (5 x 1.5 mL) and with DMF (5 x 1.2 mL).

[F] Hetereocyclisation:

To resin was added a 0.4M solution of aldehyde in DMF (0.75 mL, 0.3 mmoles, 5.0 equiv.) and 0.4 M solution of *p*-toluene sulfonic acid in DMF (0.75 mL, 0.3 mmoles, 5.0 equiv.). The reaction was agitated at ambient temperature for 5 h and the syringe was thereafter drained under reduced pressure. Resin was washed with DMF (3 x 1.2 mL).

[G] Amide Coupling:

To resin was added a 1M solution of bromoacetic acid in DMF (0.6 mL, 0.6 mmoles, 10.0 equiv.) and 1M DIC in DMF (0.6 mL, 0.6 mmoles, 10.0 equiv.). The reaction was agitated at ambient temperature for 2 h, and thereafter resin was subject to washes with DMF (3 x 1.2 mL).

[H] Nucleophilic Substitution ($S_N2$):

To resin was added a 0.4M solution of piperazine in DMF (1.5 mL, 0.6 mmoles, 10.0 equiv.). The reaction was agitated at ambient temperature for 2 h, and thereafter resin was subject to washes with DMF (3 x 1.2 mL).

**Manual Transformations**

The following steps proceeded automated synthesis and were performed manually in parallel.

[I] Linker Cleavage:

Resin was washed with DCM (5 x 2 mL), then capped and treated with cleavage cocktail (1 mL). The reaction was agitated for 1 h at ambient temperature. Eluent was collected from syringe, washed with DCM (1 mL) and evaporated to dryness under a flow of compressed air. Crude residue was dissolved in DMF (0.2 mL) prior to purification.

[J] Pyridine N-Oxidation:

Dimethylpyridine-containing intermediate (17 µmol, 1.0 equiv.) in DCM (5 mL) was treated with mCPBA (26 µmol, 1.5 equiv.) at 0 °C, and allowed to warm to rt while stirring overnight. After the formation of the target product was confirmed by HPLC-MS, the reaction was concentrated *in vacuo*, the crude residue was redissolved in DMF (0.2 mL) and purified by reverse-phase chromatography with a gradient of 5-100% MeCN/MQ (0.1% TFA).

[K1] Cyclic imide formation (HATU):

C-terminal Asp/Glu-containing precursors (1.0 equiv.) were prepared as 0.2M solutions in THF, to which was added 0.4M HATU in DMF (3.0 equiv.) and DMAP (3.0 equiv.). The reaction was stirred at 60 °C for 5 h, concentrated in vacuo and purified by reverse-phase chromatography with a stepwise gradient of 5-100% MeCN/MQ

[K2] Cyclic imide formation (CDI):

C-terminal Asp containing precursor (1.0 equiv.) were prepared as 0.2M solutions in THF, to which was added 0.4M CDI in DMF (3.0 equiv.) and DMAP (1.0 equiv.). The reaction was stirred at 60 °C overnight, concentrated *in vacuo* and purified by reverse-phase chromatography with a stepwise gradient of 5-100% MeCN/MQ

**Semi-automated purification**:

**C-18 Column chromatography***:* Crude products were purified by automated reverse phase chromatography using a Biotage Selekt System with a pre-packed column (Biotage® Sfär Bio 10 g, C18 - Duo 300 Å 20 µm). A standardized 5–100% MeCN/MilliQ $H_2O$ (0.1% TFA), 18 min gradient was applied and UV absorbance was measured at 254 nm.

**Sample Preparation***:* Column fractions containing a UV-active product at 254 nm and a matching mass ion for the target product were concentrated *in vacuo* and lyophilized to dryness. 10 mM DMSO stock solutions were prepared and stored at -20 °C.

**Pharmacophore Hybrid Modelling**:

The pose of a reported dimethyl pyridone-based ligand engaging the KAc pocket of BRD4 was analysed from the crystal structure 6ZCI.[6,7] A model for docking pharmacophore hybrids in the KAc pocket was designed from 6ZCI using the browser-based tool Pharmit,[8] where the following node configurations were defined:

1. An aromatic ring sandwiched between Val87 and Ile146; (-11.5, -5.8, -1.1), r = 1.0
2. H-bond acceptor group orientated to Asn140; (-13.8, -4.6, -1.1), r = 1.2
3. Hydrophobic group adjacent to Phe83 and Ile146; (-11.7, -3.3, -2.5), r = 1.3
4. Hydrophobic group adjacent to the WPF shelf; (-8.3, -6.1, 2.5), r = 1.5

Pharmit was used to align each conformer to the defined pharmacophoric nodes. An energy-minimization step was applied, followed by filtering using an energy score threshold (< –6) and a maximum mRMSD filter (< 3.0 Å) to remove low-quality poses.

Compound **6** was aligned to the defined nodes and ranked using Pharmit's integrated AutoDock Vina scoring function. The top-ranked pose was used to generate a model for the benzimidazole series, in which alternative KAc pocket-binding groups could be modelled.

### Building Blocks

### Amino Acids

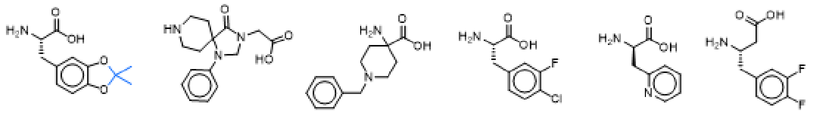

AA1 AA2 AA3 AA4 AA5 AA6

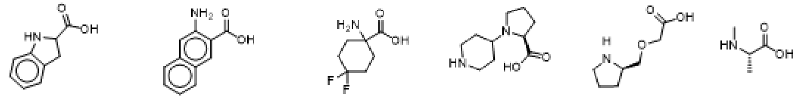

AA7 AA8 AA9 AA10 AA11 AA12

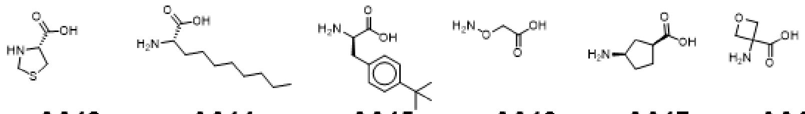

AA13 AA14 AA15 AA16 AA17 AA18

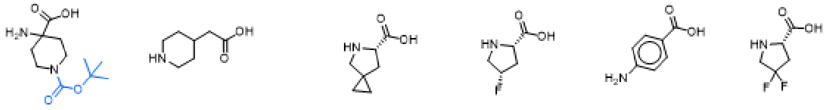

AA19 AA20 AA21 AA22 AA23 AA24

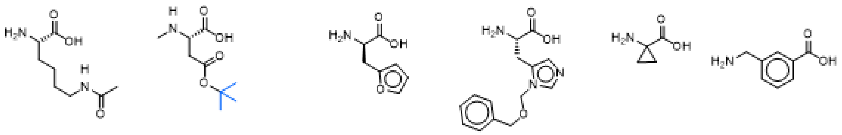

AA25 AA26 AA27 AA28 AA29 AA30

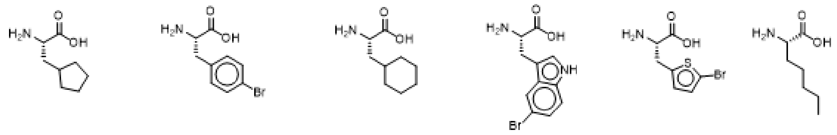

AA31 AA32 AA33 AA34 AA35 AA36

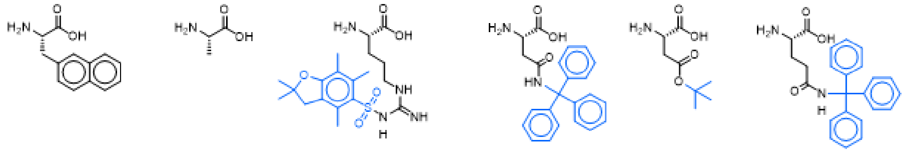

AA37 AA38 AA39 AA40 AA41 AA42

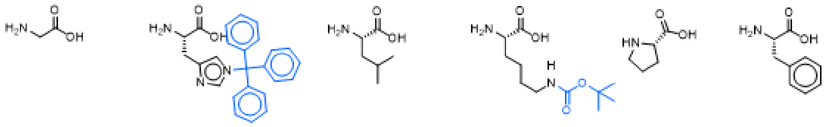

AA43 AA44 AA45 AA46 AA47 AA48

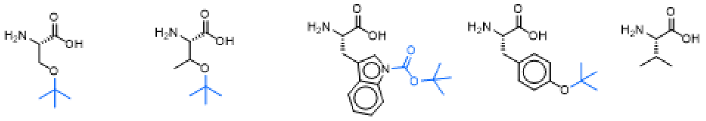

AA49 AA50 AA51 AA52 AA53

### Aldehydes

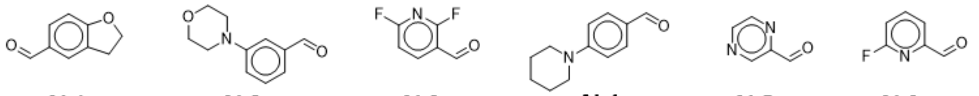

AL1 AL2 AL3 AL4 AL5 AL6

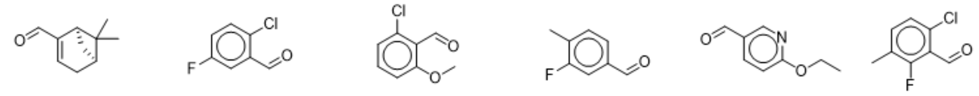

AL7 AL8 AL9 AL10 AL11 AL12

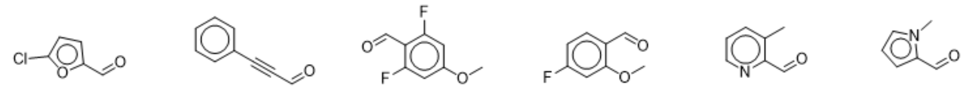

AL13 AL14 AL15 AL16 AL17 AL18

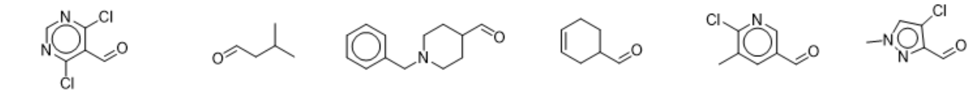

AL19 AL20 AL21 AL22 AL23 AL24

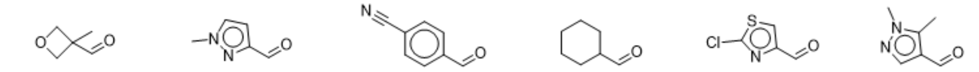

AL25 AL26 AL27 AL28 AL29 AL30

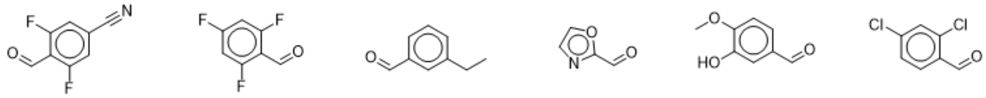

AL31 AL32 AL33 AL34 AL35 AL36

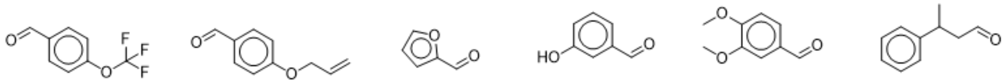

AL37 AL38 AL39 AL40 AL41 AL42

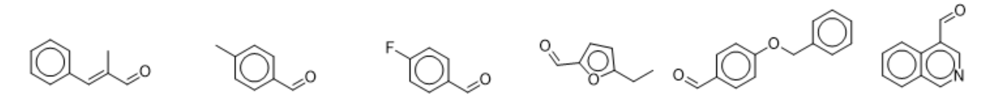

AL43 AL44 AL45 AL46 AL47 AL48

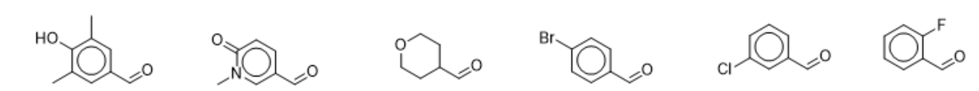

AL49 AL50 AL51 AL52 AL53 AL54

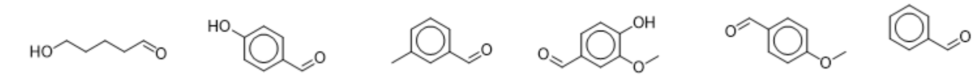

AL55 AL56 AL57 AL58 AL59 AL60

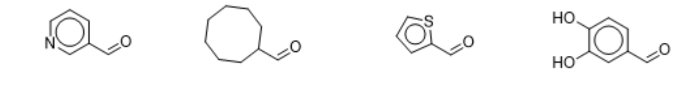

AL61 AL62 AL63 AL64

### Primary Amines

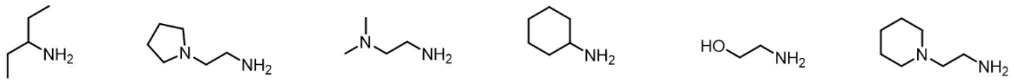
**AM1** **AM2** **AM3** **AM4** **AM5** **AM6**

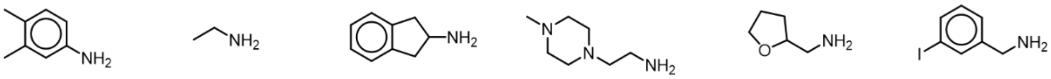
**AM7** **AM8** **AM9** **AM10** **AM11** **AM12**

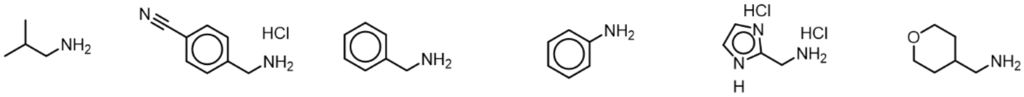
**AM13** **AM14** **AM15** **AM16** **AM17** **AM18**

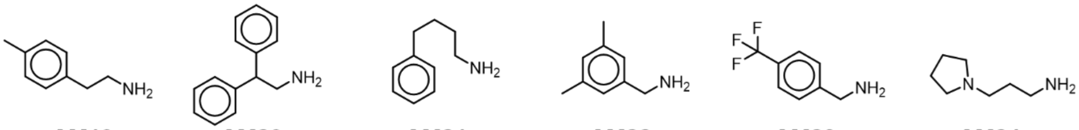
**AM19** **AM20** **AM21** **AM22** **AM23** **AM24**

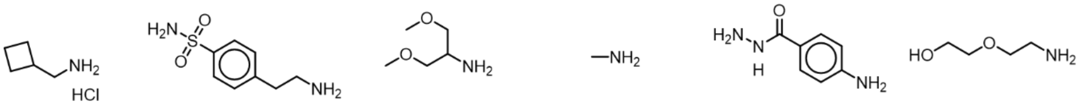
**AM25** **AM26** **AM27** **AM28** **AM29** **AM30**

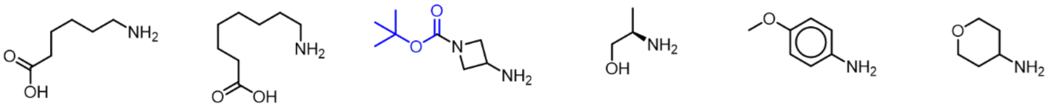
**AM31** **AM32** **AM33** **AM34** **AM35** **AM36**

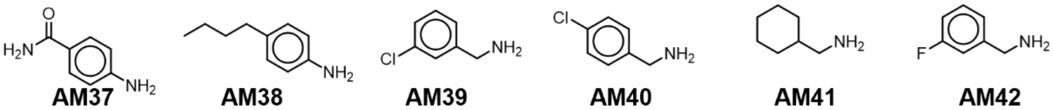
**AM37** **AM38** **AM39** **AM40** **AM41** **AM42**

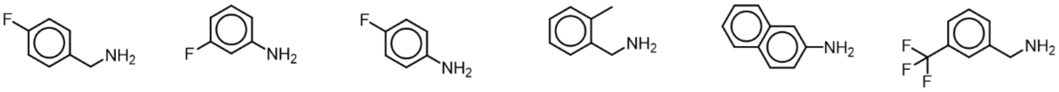
**AM43** **AM44** **AM45** **AM46** **AM47** **AM48**

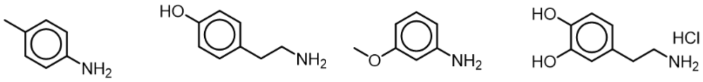
**AM49** **AM50** **AM51** **AM52**

### Synthetic Sequences

#### Synthetic Sequence for Compounds 1-32

| Compound | Automated Synthesis Sequence | Manual Sequence | Mass (mg) | µmoles | Yield (%) |
|---|---|---|---|---|---|
| **1** | [A][B][C][D][E][F] | [I] | 2.5 | 5.4 | 8.9 |
| **2** | [A][B][C][D][E][F] | [I] | 2.9 | 5.9 | 9.8 |
| **3** | [A][B][C][D][E][F] | [I] | 5.8 | 11.5 | 17.4 |
| **4** | [A][B][C][D][E][F] | [I] | 4.8 | 9.7 | 16.7 |
| **5** | [A][B][C][D][E][F] | [I] | 6.1 | 14.1 | 23.5 |
| **6** | [A][B][C][D][E][F] | [I] | 5.1 | 11.0 | 18.3 |
| **7** | [A][B][C][D][E][F] | [I] | 9.4 | 20.2 | 30.6 |
| **8** | [A][B][C][D][E][F] | [I] | 6.8 | 14.1 | 21.4 |
| **9** | [A][B][C][D][E][F] | [I] | 7.5 | 15.3 | 23.2 |
| **10** | [A][B][C][D][E][F] | [I] | 8.3 | 16.3 | 24.6 |
| **11** | [A][B][C][D][E][F] | [I] | 2.3 | 4.6 | 7.0 |
| **12** | [A][B][C][D][E][F] | [I] | 3.2 | 6.5 | 9.8 |
| **13** | [A][B][C][D][E][F] | [I] | 5.3 | 10.5 | 15.9 |
| **14** | [A][B][C][D][E][F] | [I] | 8.8 | 17.0 | 25.8 |
| **15** | [A][B][C][D][E][F] | [I] | 3.3 | 6.7 | 12.6 |
| **16** | [A][B][C][D][E][F] | [I] | 11.9 | 23.5 | 35.6 |
| **17** | [A][B][C][D][E][F] | [I] | 3.2 | 6.2 | 11.7 |
| **18** | [A][B][C][D][E][F] | [I] | 3.4 | 6.3 | 12.0 |
| **19** | [A][B][C][D][E][F] | [I] | 10.2 | 18.7 | 28.3 |
| **20** | [A][B][C][D][E][F] | [I] | 10.9 | 20.1 | 30.4 |
| **21** | [A][B][C][D][E][F] | [I] | 4.4 | 8.5 | 16.1 |
| **22** | [A][B][C][D][E][F] | [I] | 4.2 | 7.8 | 14.7 |
| **23** | [A][B][C][D][E][F] | [I] | 6.8 | 12.7 | 24.1 |
| **24** | [A][B][C][D][E][F] | [I] | 5.2 | 10.5 | 15.9 |
| **25** | [A][B][C][D][E][F] | [I] | 18.2 | 37.0 | 56.1 |
| **26** | [A][B][C][D][E][F] | [I][J] | 2.8 | 5.4 | 17.8 |
| **27** | [A][B][C][D][E][F] | [I] | 6.7 | 11.3 | 17.2 |
| **28** | [A][B][C][D][E][F] | [I] | 7.4 | 16.4 | 24.9 |
| **29** | [A][B][C][D][E][F] | [I] | 6.4 | 12.1 | 18.3 |
| **30** | [A][B][C][D][E][F] | [I] | 0.6 | 1.1 | 1.6 |
| **31** | [A][B][C][D][E][F] | [I] | 7.8 | 15.8 | 24.0 |
| **32** | [A][B][C][D][E][F] | [I] | 8.2 | 16.0 | 24.2 |

### Synthetic Sequence for Compounds 33-49

| Compound | Automated Synthesis Sequence | Manual Sequence | Mass (mg) | µmoles | Yield (%) |
|---|---|---|---|---|---|
| **33** | [A][B][A][B][C][D][E][F] | [I] | 3.5 | 5.9 | 9.9 |
| **34** | [A][B][A][B][A][B][C][D][E][F] | [I] | 6.9 | 9.6 | 15.9 |
| **35** | [A][B][A][B][A][B][C][D][E][F] | [I] | 6.3 | 8.5 | 14.1 |
| **36** | [A][B][A][B][A][B][C][D][E][F] | [I][J] | 2.8 | 3.9 | 6.5 |
| **37** | [A][B][A][B][A][B][A][B][C][D][E][F] | [I] | 11.5 | 13.8 | 23.0 |
| **38** | [A][B][A][B][A][B][A][B][C][D][E][F] | [I] | 13.1 | 15.9 | 26.5 |
| **39** | [A][B][A][B][A][B][A][B][C][D][E][F] | [I] | 4.7 | 5.5 | 9.1 |
| **40** | [A][B][A][B][A][B][A][B][C][D][E][F] | [I] | 17.3 | 20.0 | 33.3 |
| **41** | [A][B][A][B][A][B][A][B][C][D][E][F] | [I] | 6.1 | 7.3 | 12.2 |
| **42** | [A][B][A][B][A][B][A][B][C][D][E][F] | [I] | 3.6 | 4.1 | 6.9 |
| **43** | [A][B][A][B][A][B][A][B][C][D][E][F] | [I] | 9.9 | 12.7 | 21.2 |
| **44** | [A][B][A][B][A][B][A][B][A][B][C][D][E][F] | [I] | 5.4 | 5.6 | 9.3 |
| **45** | [A][B][A][B][A][B][A][B][A][B][C][D][E][F] | [I] | 7.8 | 8.1 | 13.4 |
| **46** | [A][B][A][B][A][B][A][B][A][B][C][D][E][F] | [I] | 7.2 | 8.6 | 14.3 |
| **47** | [A][B][A][B][G][H][A][B][A][B][A][B][C][D][E][F] | [I] | 0.9 | 0.9 | 1.6 |
| **48** | [A][B][A][B][A][B][A][B][C][D][E][F] | [I] | 20.4 | 24.1 | 40.1 |
| **49** | [A][B][A][B][A][B][A][B][C][D][E][F] | [I][J] | 0.7 | 0.8 | 2.2 |

### Synthetic Sequence for Compounds 51-66

| Compound | Automated Synthesis Sequence | Manual Sequence | Mass (mg) | µmoles | Yield (%) |
|---|---|---|---|---|---|
| **51** | [A][B][A][B][C][D][E][F] | [I][K1/K2] | 1.9 | 3.1 | 12.2 |
| **52** | [A][B][C][D][E][F] | [I][K2] | 7.0 | 14.7 | 12.2 |
| **53** | [A][B][A][B][C][D][E][F] | [I][K1/K2] | 2.2 | 3.3 | 7.2 |
| **54** | [A][B][A][B][A][B][C][D][E][F] | [I][K2] | 1.3 | 1.7 | 2.8 |
| **55** | [A][B][A][B][A][B][C][D][E][F] | [I][K2] | 0.5 | 0.6 | 0.9 |
| **56** | [A][B][A][B][A][B][C][D][E][F] | [I][K2] | 0.9 | 1.1 | 1.8 |
| **57** | [A][B][A][B][A][B][C][D][E][F] | [I][K2] | 2.5 | 3.3 | 5.4 |
| **58** | [A][B][A][B][A][B][A][B][C][D][E][F] | [I][K2] | 2.1 | 2.4 | 3.9 |
| **59** | [A][B][A][B][A][B][A][B][C][D][E][F] | [I][K1] | 1.6 | 2.0 | 3.2 |
| **60** | [A][B][A][B][A][B][A][B][C][D][E][F] | [I][K1][J] | 1.5 | 1.8 | 1.4 |
| **61** | [A][B][A][B][A][B][A][B][C][D][E][F] | [I][K1][J] | 1.7 | 2.0 | 1.6 |
| **62** | [A][B][A][B][A][B][A][B][C][D][E][F] | [I][K1] | 1.9 | 2.3 | 3.6 |
| **63** | [A][B][A][B][A][B][A][B][C][D][E][F] | [I][K1] | 1.4 | 1.8 | 2.8 |
| **64** | [A][B][A][B][A][B][A][B][C][D][E][F] | [I][K1] | 3.1 | 3.6 | 5.7 |
| **65** | [A][B][A][B][A][B][A][B][C][D][E][F] | [I][K1] | 1.9 | 2.4 | 3.8 |
| **66** | [A][B][A][B][A][B][A][B][C][D][E][F] | [I][K1] | 2.9 | 3.6 | 13.9 |

### Synthetic Sequence for Compounds 50, 67-70

| Compound | Automated Synthesis Sequence | Manual Sequence | Mass (mg) | µmoles | Yield (%) |
|---|---|---|---|---|---|
| **50** | [A][B][A][B][C][D][E][F] | [I] | 11.9 | 18.6 | 31.1 |
| **67** | [A][B][A][B][A][B][C][D][E][F] | [I] | 11.2 | 14.3 | 23.8 |
| **68** | [A][B][A][B][A][B][A][B][C][D][E][F] | [I] | 13.2 | 16.0 | 26.7 |
| **69** | [A][B][A][B][A][B][A][B][C][D][E][F] | [I] | 6.4 | 7.6 | 12.7 |
| **70** | [A][B][A][B][A][B][A][B][C][D][E][F] | [I] | 23.2 | 28.3 | 47.1 |

### Analytical Data

### HPLC-MS Data for compounds 1-70

**1:** Exact mass: 474.26; observed $[M + H]^+$: 475.25

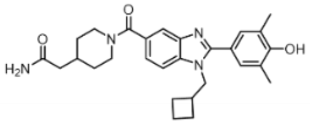

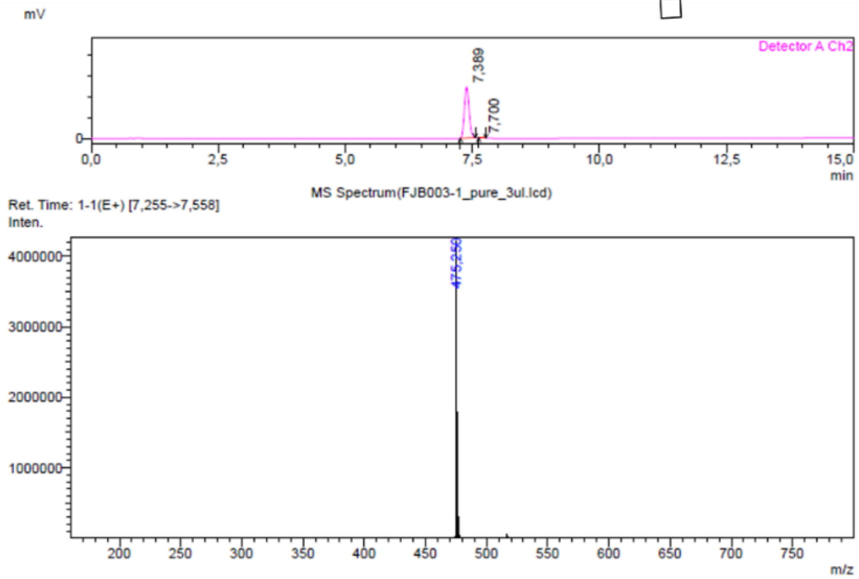

Peak Table(FJB003-1_pure_3ul.lcd)

Detector A Channel 2 254nm

| Peak# | Ret. Time | Area | Height | Mark | Conc. | Area% |
|---|---|---|---|---|---|---|
| 1 | 7,389 | 7961922 | 1218600 | M | 99,031 | 99,031 |
| 2 | 7,700 | 77938 | 17303 | M | 0,969 | 0,969 |
| Total | | 8039860 | 1235903 | | 100,000 | 100,000 |

**2:** Exact mass: 488.28; observed $[M + H]^+$: 489.20

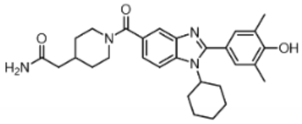

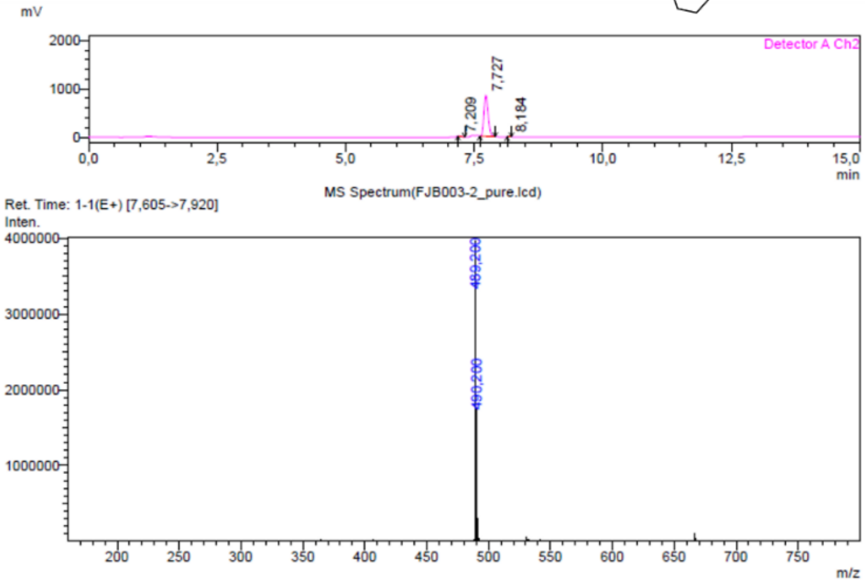

Peak Table(FJB003-2_pure.lcd)

Detector A Channel 2 254nm

| Peak# | Ret. Time | Area | Height | Mark | Conc. | Area% |
|---|---|---|---|---|---|---|
| 1 | 7,209 | 24105 | 6761 | M | 0,496 | 0,496 |
| 2 | 7,727 | 4821903 | 844509 | M | 99,173 | 99,173 |
| 3 | 8,184 | 16092 | 5041 | M | 0,331 | 0,331 |
| Total | | 4862099 | 856311 | | 100,000 | 100,000 |

**3:** Exact mass: 505.27; observed $[M + H]^+$: 506.25

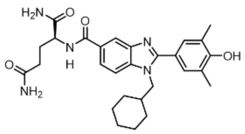

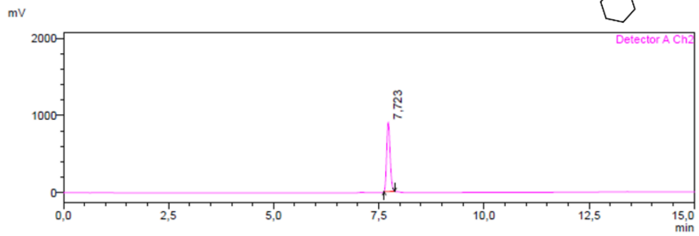

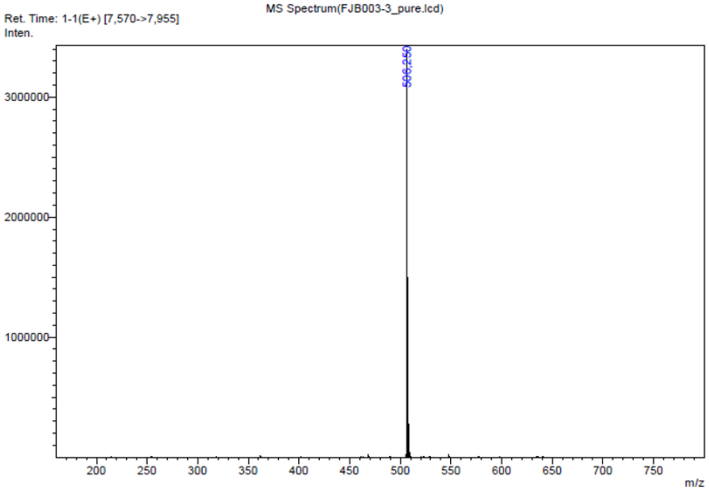

**4:** Exact mass: 492.25; observed $[M + H]^+$: 493.20

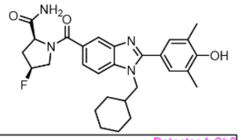

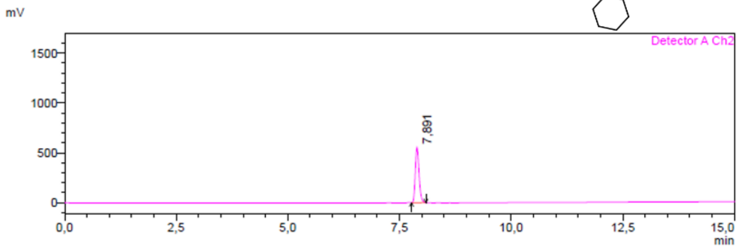

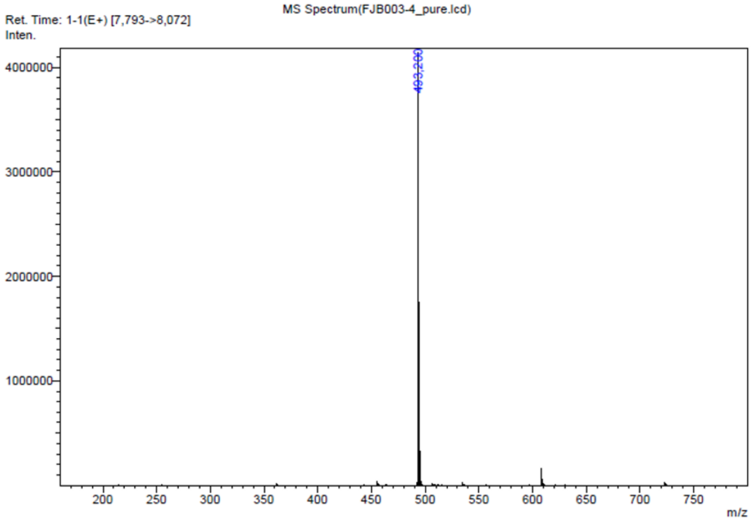

**5:** Exact mass: 492.24; observed $[M + H]^{+}$: 493.20

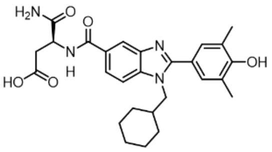

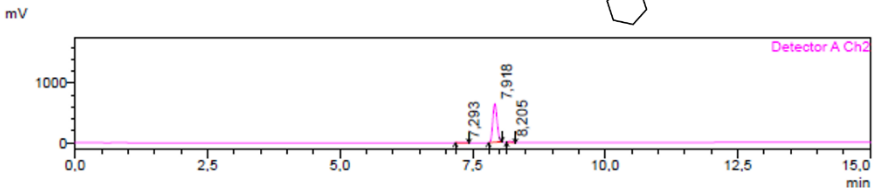

MS Spectrum(FJB003-5_pure.lcd)

Ret. Time: 1-1(E+) [7,815->8,072]
Inten.

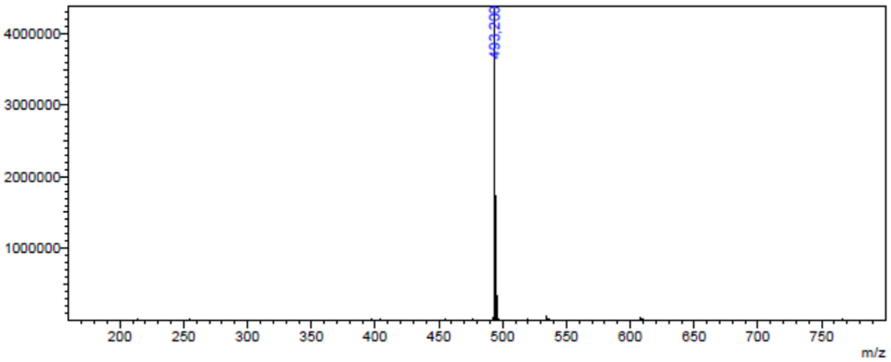

Peak Table(FJB003-5_pure.lcd)

Detector A Channel 2 254nm

| Peak# | Ret. Time | Area | Height | Mark | Conc. | Area% |
|---|---|---|---|---|---|---|
| 1 | 7,293 | 41900 | 6100 | M | 1,114 | 1,114 |
| 2 | 7,918 | 3618418 | 642689 | M | 96,208 | 96,208 |
| 3 | 8,205 | 100704 | 20226 | M | 2,678 | 2,678 |
| Total | | 3761021 | 669015 | | 100,000 | 100,000 |

**6:** Exact mass: 462.26; observed $[M + H]^{+}$: 463.20

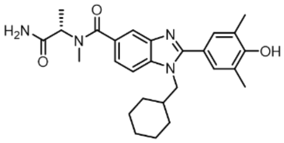

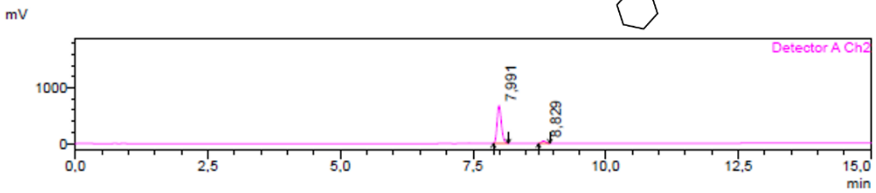

MS Spectrum(FJB003-6_pure.lcd)

Ret. Time: 1-1(E+) [7,885->8,130]
Inten.

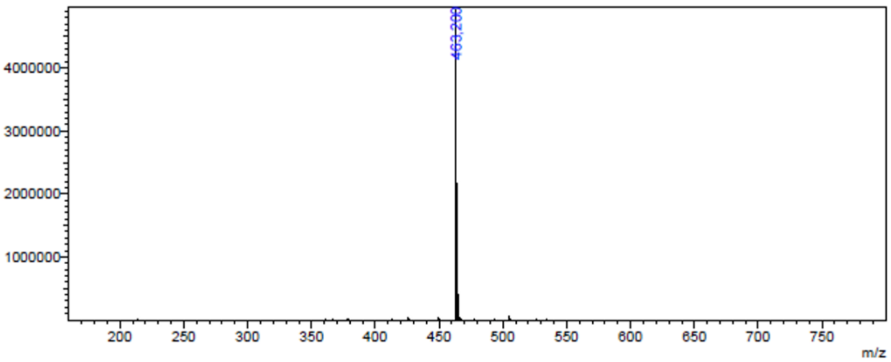

Peak Table(FJB003-6_pure.lcd)

Detector A Channel 2 254nm

| Peak# | Ret. Time | Area | Height | Mark | Conc. | Area% |
|---|---|---|---|---|---|---|
| 1 | 7,991 | 3827825 | 677751 | M | 93,889 | 93,889 |
| 2 | 8,829 | 249151 | 43538 | M | 6,111 | 6,111 |
| Total | | 4076975 | 721288 | | 100,000 | 100,000 |

**7:** Exact mass: 463.22; observed $[M + H]^{+}$: 464.20

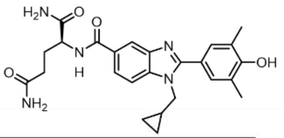

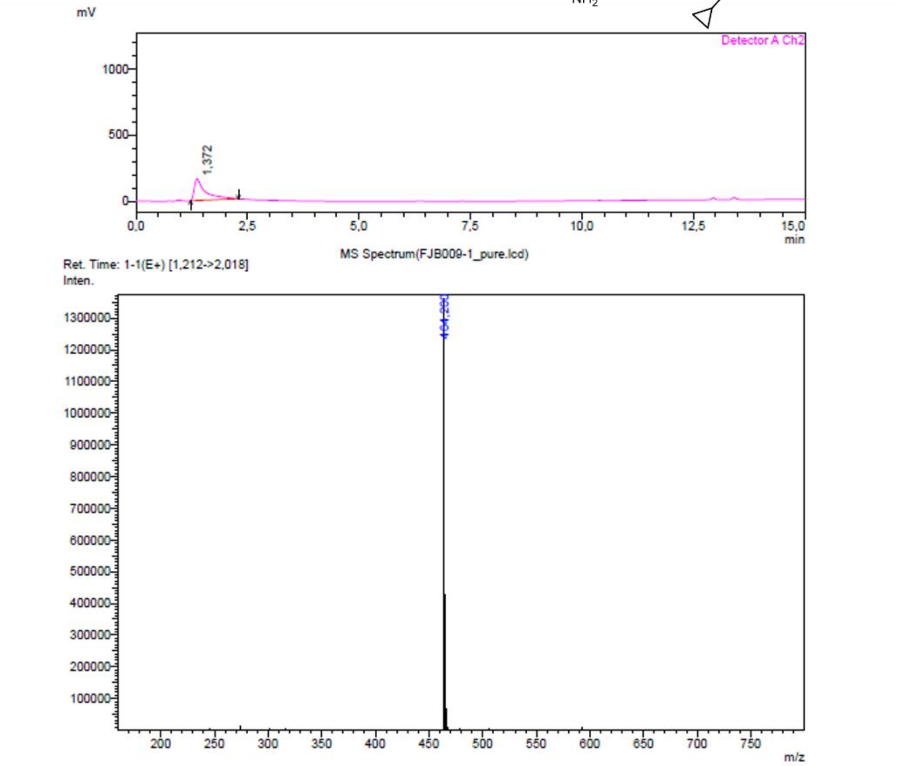

**8:** Exact mass: 479.25; observed $[M + H]^{+}$: 480.20

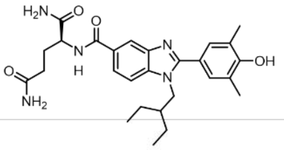

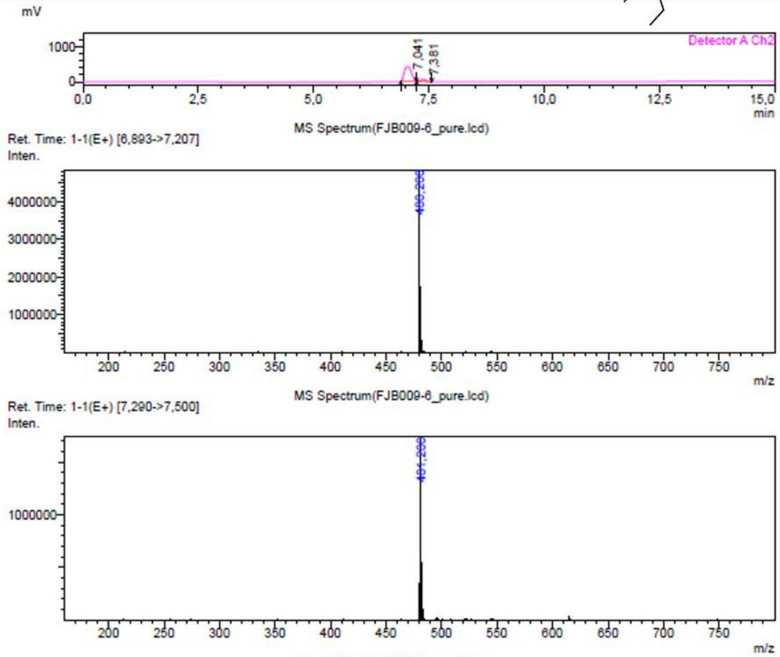

Peak Table(FJB009-6_pure.lcd)

Detector A Channel 2 254nm

| Peak# | Ret. Time | Area | Height | Mark | Conc. | Area% |
|---|---|---|---|---|---|---|
| 1 | 7,041 | 4566771 | 423189 | M | 91,060 | 91,060 |
| 2 | 7,381 | 448341 | 70714 | M | 8,940 | 8,940 |
| Total | | 5015113 | 493903 | | 100,000 | 100,000 |

**9:** Exact mass: 491.25; observed $[M + H]^+$: 492.20

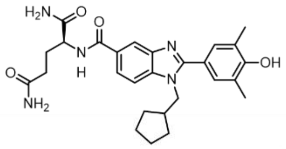

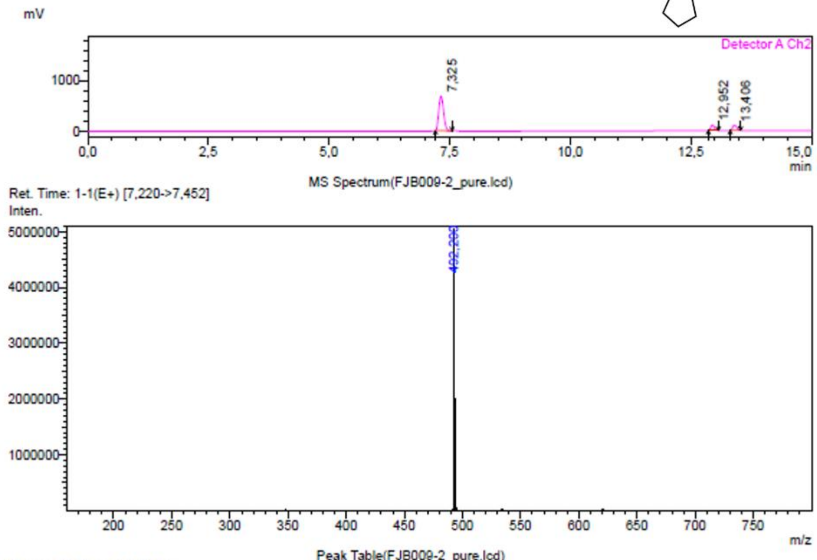

Peak Table(FJB009-2_pure.lcd)

Detector A Channel 2 254nm

| Peak# | Ret. Time | Area | Height | Mark | Conc. | Area% |
|---|---|---|---|---|---|---|
| 1 | 7,325 | 4753024 | 683126 | M | 79,938 | 79,938 |
| 2 | 12,952 | 586664 | 113105 | M | 9,867 | 9,867 |
| 3 | 13,406 | 606163 | 104458 | M | 10,195 | 10,195 |
| Total | | 5945851 | 900688 | | 100,000 | 100,000 |

**10:** Exact mass: 507.25; observed $[M + H]^+$: 508.20

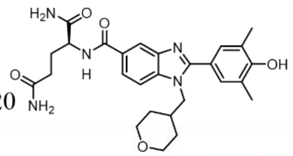

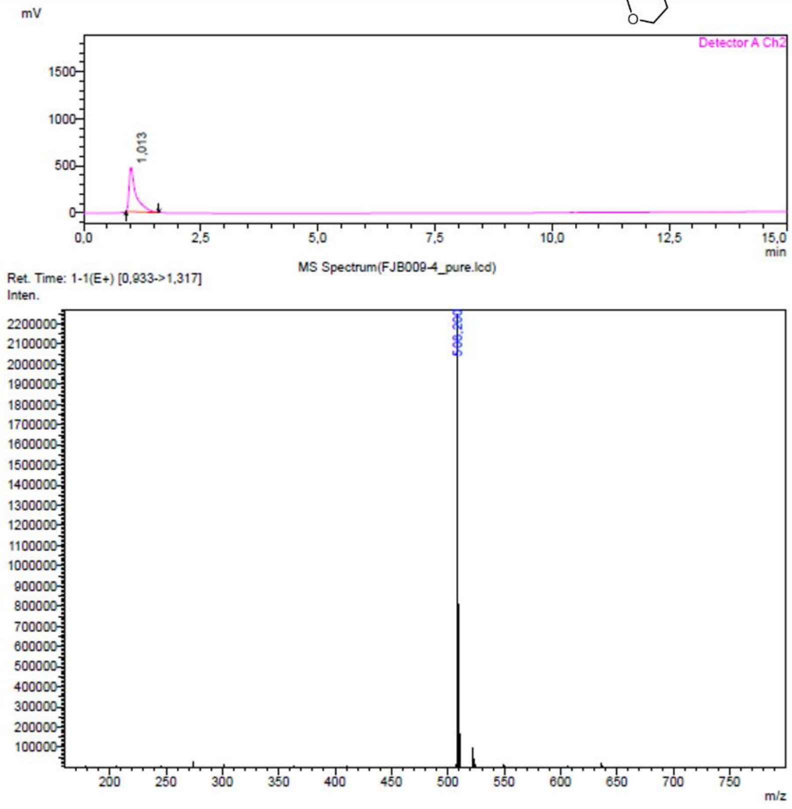

**11:** Exact mass: 507.25; observed $[M + H]^+$: 508.20

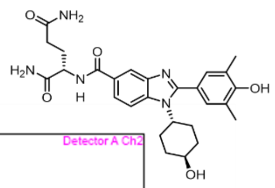

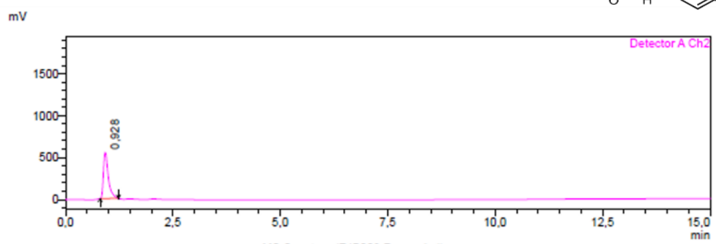

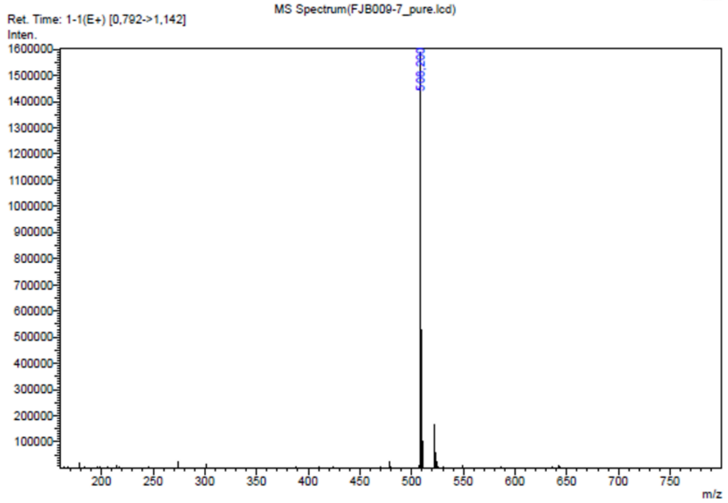

**12:** Exact mass: 493.23; observed $[M + H]^+$: 494.20

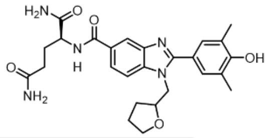

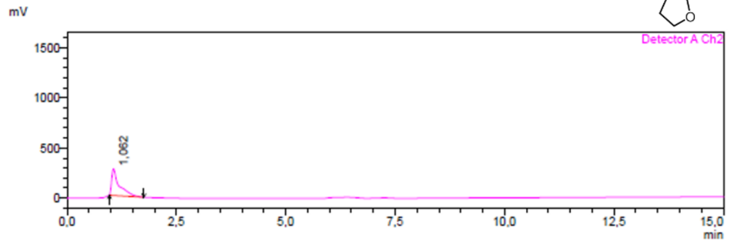

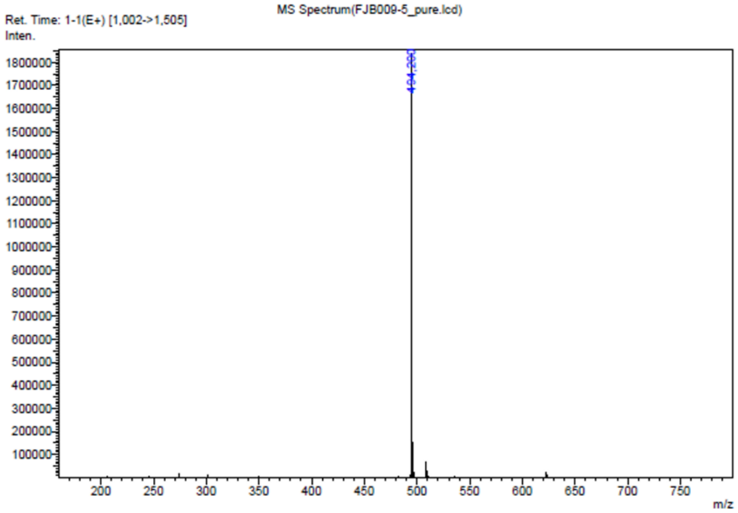

**13:** Exact mass: 499.22; observed $[M + H]^+$: 500.15 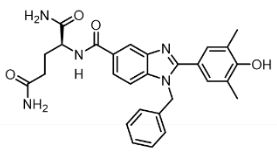

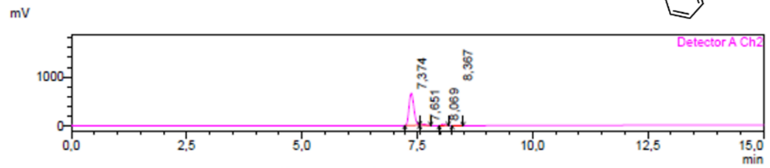

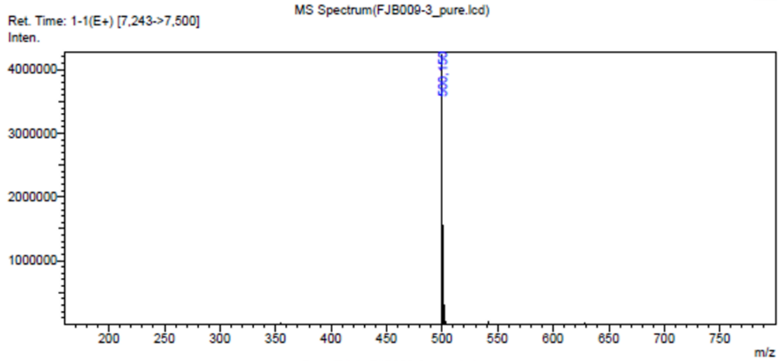

Peak Table(FJB009-3_pure.lcd)

Detector A Channel 2 254nm

| Peak# | Ret. Time | Area | Height | Mark | Conc. | Area% |
|---|---|---|---|---|---|---|
| 1 | 7,374 | 4823127 | 659815 | M | 93,469 | 93,469 |
| 2 | 7,651 | 106818 | 20870 | M | 2,070 | 2,070 |
| 3 | 8,069 | 185223 | 33033 | M | 3,590 | 3,590 |
| 4 | 8,367 | 44963 | 7932 | M | 0,871 | 0,871 |
| Total | | 5160131 | 721651 | | 100,000 | 100,000 |

**14:** Exact mass: 519.28; observed $[M + H]^+$: 520.30 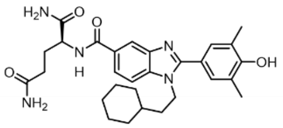

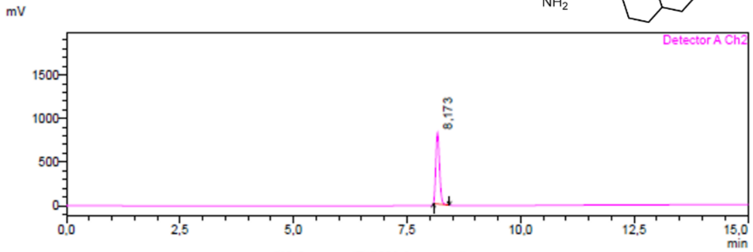

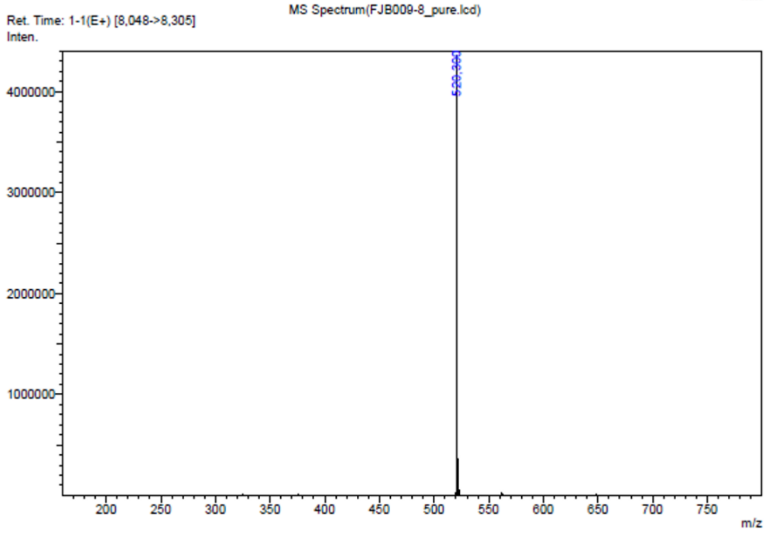

**15:** Exact mass: 491.25; observed [M + H]$^+$: 492.20

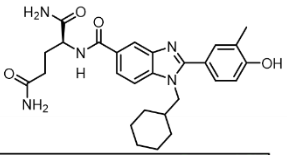

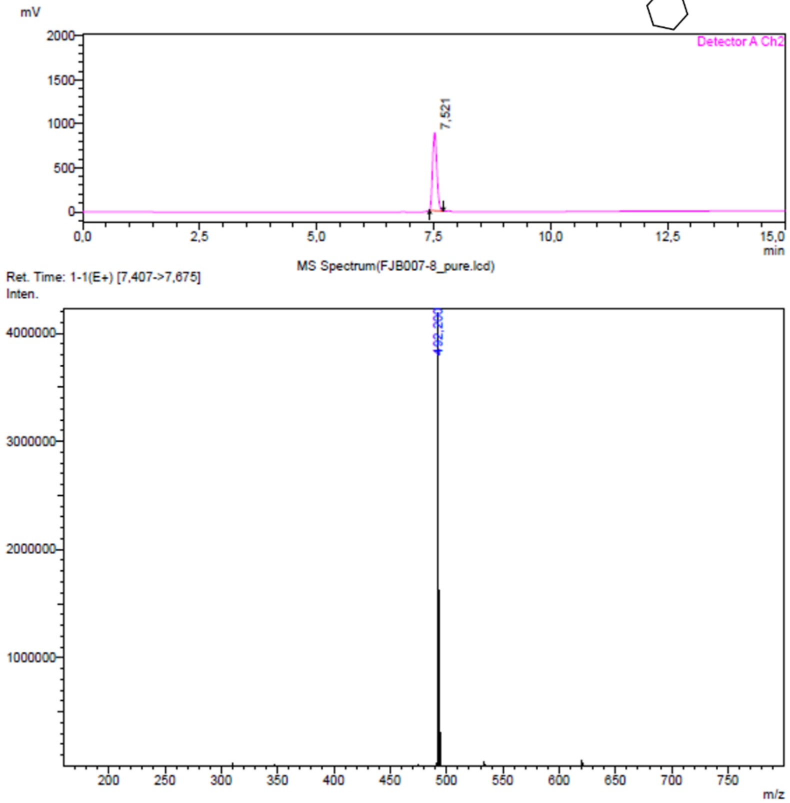

**16:** Exact mass: 505.27; observed [M + H]$^+$: 506.20

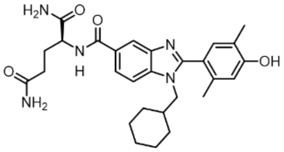

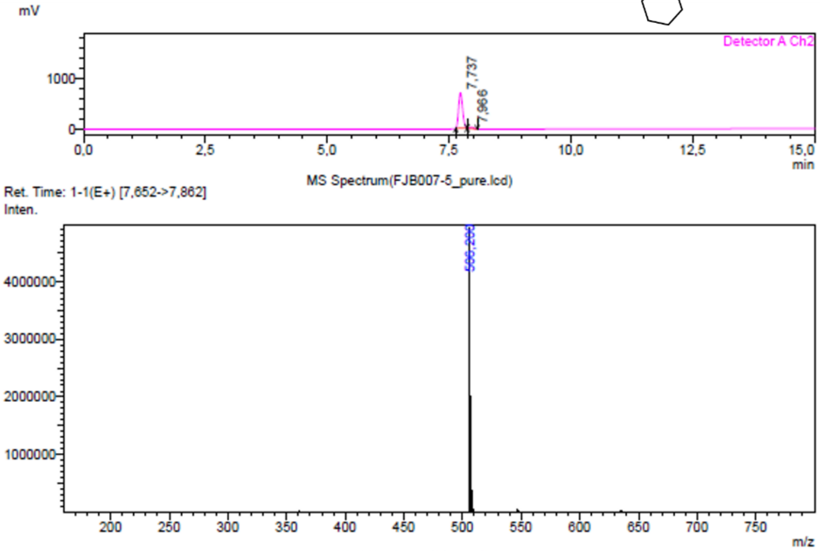

Peak Table(FJB007-5_pure.lcd)

Detector A Channel 2 254nm

| Peak# | Ret. Time | Area | Height | Mark | Conc. | Area% |
|---|---|---|---|---|---|---|
| 1 | 7.737 | 4017679 | 698052 | M | 97.212 | 97.212 |
| 2 | 7.966 | 115222 | 26441 | M | 2.788 | 2.788 |
| Total | | 4132901 | 724493 | | 100.000 | 100.000 |

**17:** Exact mass: 521.26; observed $[M + H]^{+}$: 522.25

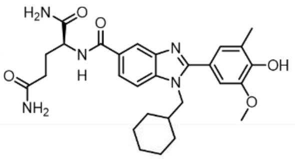

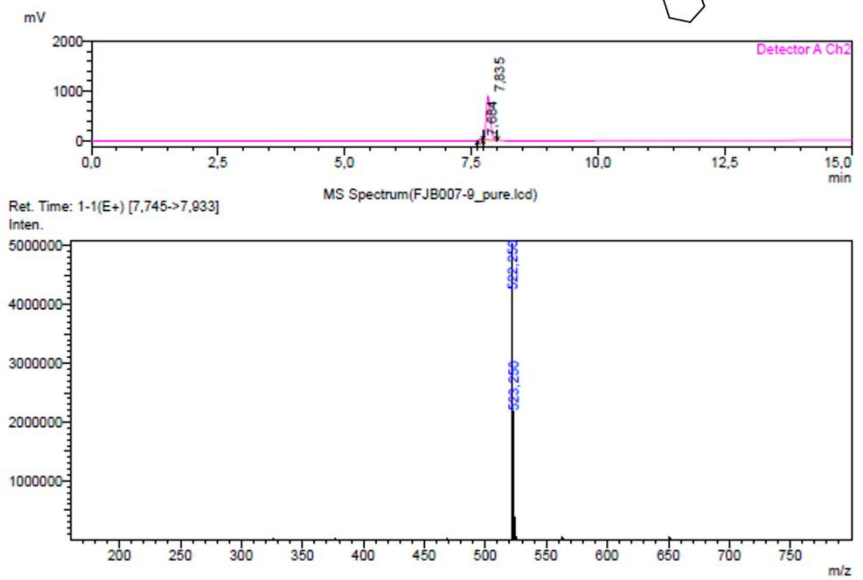

Peak Table(FJB007-9_pure.lcd)

Detector A Channel 2 254nm

| Peak# | Ret. Time | Area | Height | Mark | Conc. | Area% |
|---|---|---|---|---|---|---|
| 1 | 7.684 | 41524 | 11036 | M | 0.817 | 0.817 |
| 2 | 7.835 | 5039568 | 869027 | M | 99.183 | 99.183 |
| Total | | 5081093 | 880064 | | 100.000 | 100.000 |

**18:** Exact mass: 533.30; observed $[M + H]^{+}$: 534.25

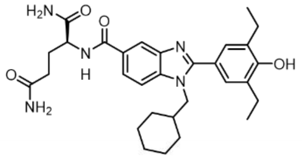

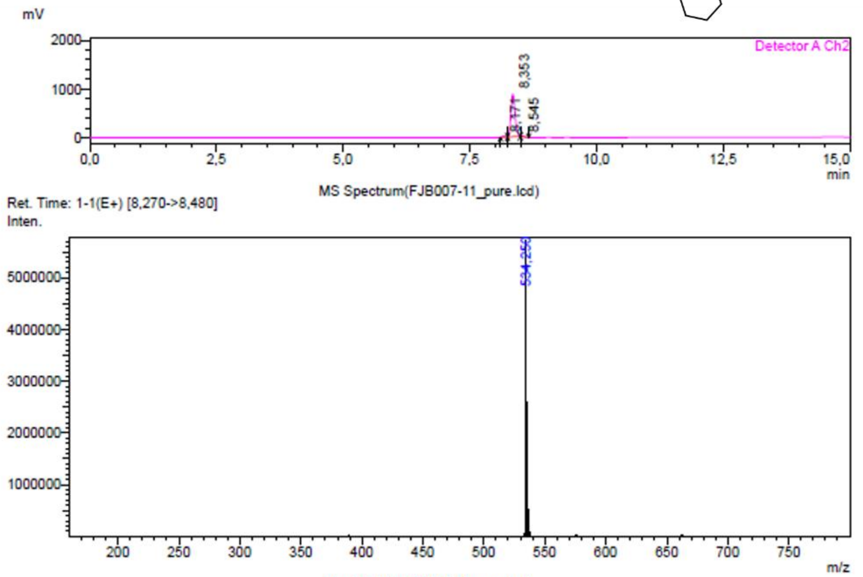

Peak Table(FJB007-11_pure.lcd)

Detector A Channel 2 254nm

| Peak# | Ret. Time | Area | Height | Mark | Conc. | Area% |
|---|---|---|---|---|---|---|
| 1 | 8.171 | 83338 | 20087 | M | 1.635 | 1.635 |
| 2 | 8.353 | 4985219 | 872257 | M | 97.829 | 97.829 |
| 3 | 8.545 | 27319 | 8407 | M | 0.536 | 0.536 |
| Total | | 5095875 | 900751 | | 100.000 | 100.000 |

**19:** Exact mass: 545.16; observed $[M + H]^+$: 546.15

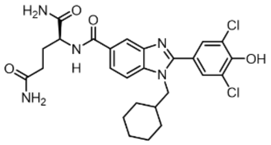

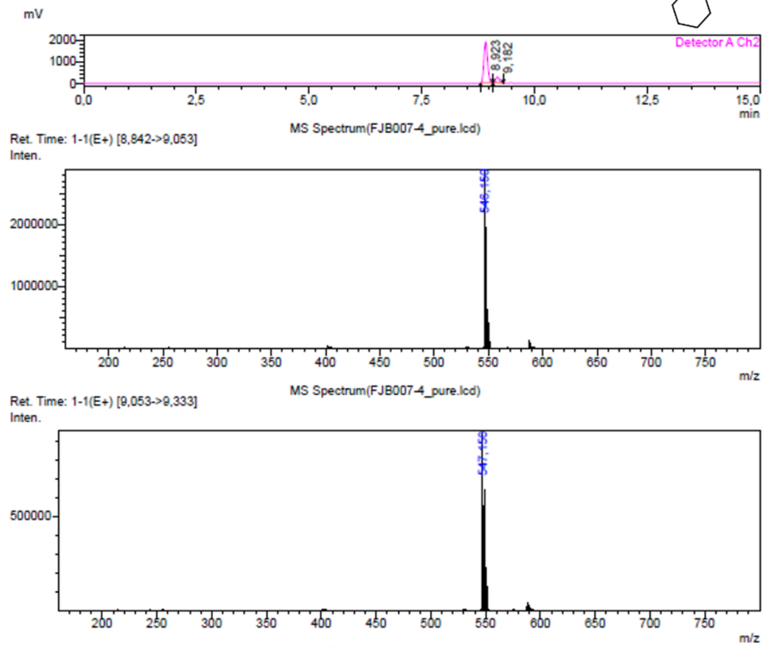

Peak Table(FJB007-4_pure.lcd)

Detector A Channel 2 254nm

| Peak# | Ret. Time | Area | Height | Mark | Conc. | Area% |
|---|---|---|---|---|---|---|
| 1 | 8,923 | 10706739 | 1886791 | M | 88,019 | 88,019 |
| 2 | 9,182 | 1457443 | 264396 | M | 11,981 | 11,981 |
| Total | | 12164182 | 2151186 | | 100,000 | 100,000 |

**20:** Exact mass: 545.22; observed $[M + H]^+$: 546.20

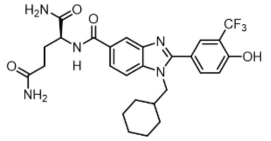

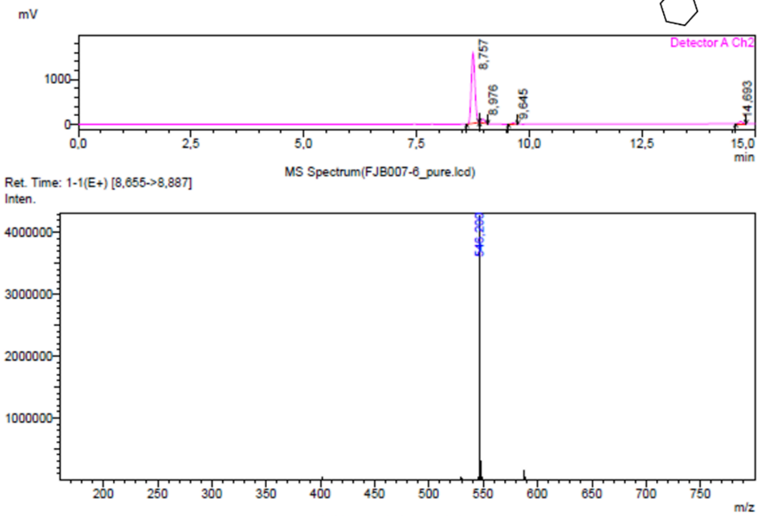

Peak Table(FJB007-6_pure.lcd)

Detector A Channel 2 254nm

| Peak# | Ret. Time | Area | Height | Mark | Conc. | Area% |
|---|---|---|---|---|---|---|
| 1 | 8,757 | 8770874 | 1561922 | M | 91,414 | 91,414 |
| 2 | 8,976 | 433945 | 88400 | M | 4,523 | 4,523 |
| 3 | 9,645 | 118822 | 22268 | M | 1,238 | 1,238 |
| 4 | 14,693 | 271056 | 48253 | M | 2,825 | 2,825 |
| Total | | 9594697 | 1720842 | | 100,000 | 100,000 |

**21:** Exact mass: 511.20; observed $[M + H]^+$: 512.15

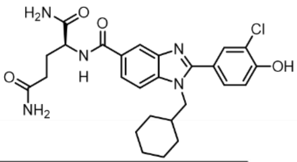

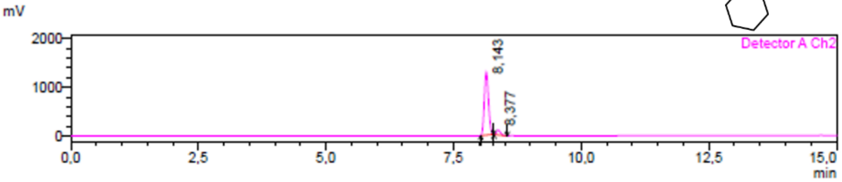

MS Spectrum(FJB007-10_pure.lcd)

Ret. Time: 1-1(E+) [8,048->8,270]
Inten.

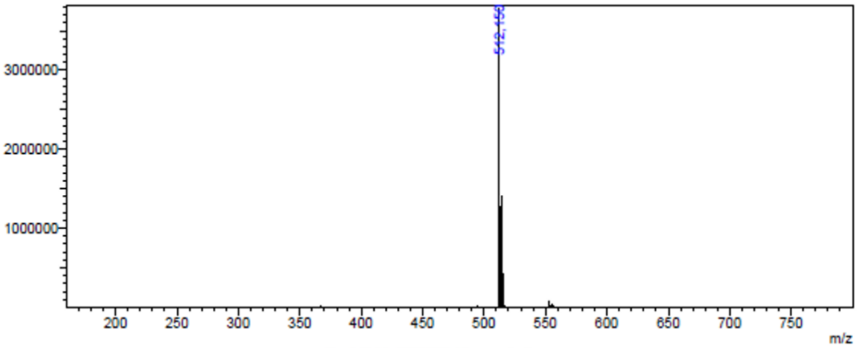

Peak Table(FJB007-10_pure.lcd)

Detector A Channel 2 254nm

| Peak# | Ret. Time | Area | Height | Mark | Conc. | Area% |
|---|---|---|---|---|---|---|
| 1 | 8.143 | 7387505 | 1292230 | M | 93.524 | 93.524 |
| 2 | 8.377 | 511513 | 100772 | M | 6.476 | 6.476 |
| Total | | 7899018 | 1393003 | | 100.000 | 100.000 |

**22:** Exact mass: 541.21; observed $[M + H]^+$: 542.15

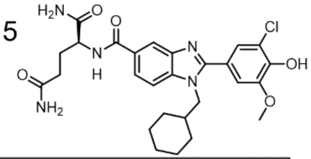

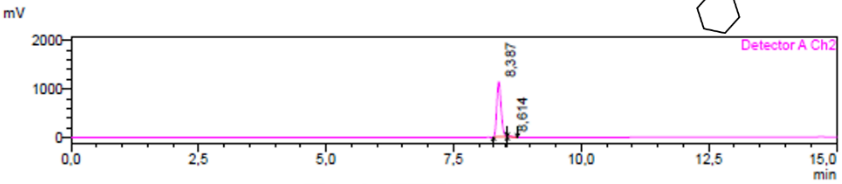

MS Spectrum(FJB007-7_pure.lcd)

Ret. Time: 1-1(E+) [8,283->8,492]
Inten.

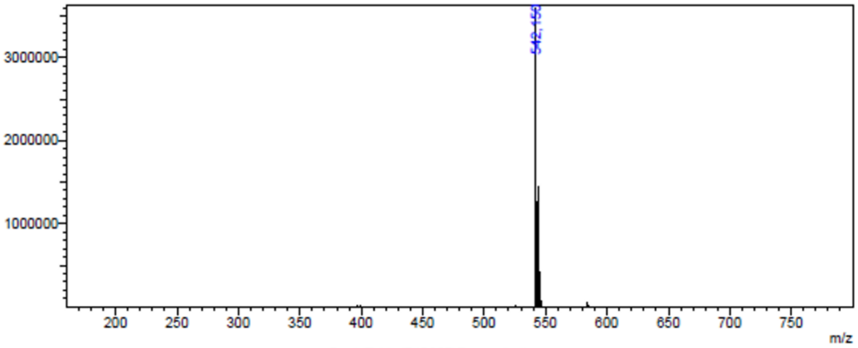

Peak Table(FJB007-7_pure.lcd)

Detector A Channel 2 254nm

| Peak# | Ret. Time | Area | Height | Mark | Conc. | Area% |
|---|---|---|---|---|---|---|
| 1 | 8.387 | 6468569 | 1131818 | M | 99.422 | 99.422 |
| 2 | 8.614 | 37603 | 10152 | M | 0.578 | 0.578 |
| Total | | 6506173 | 1141971 | | 100.000 | 100.000 |

**23:** Exact mass: 529.19; observed $[M + H]^{+}$: 530.15

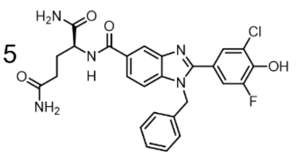

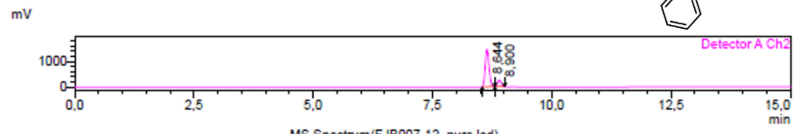

MS Spectrum(FJB007-12_pure.lcd)

Ret. Time: 1-1(E+) [8,527->8,795]
Inten.

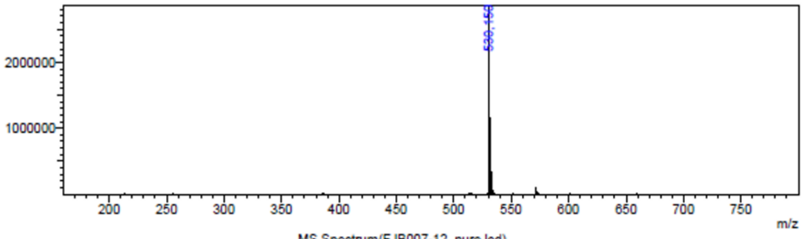

MS Spectrum(FJB007-12_pure.lcd)

Ret. Time: 1-1(E+) [8,795->8,992]
Inten.

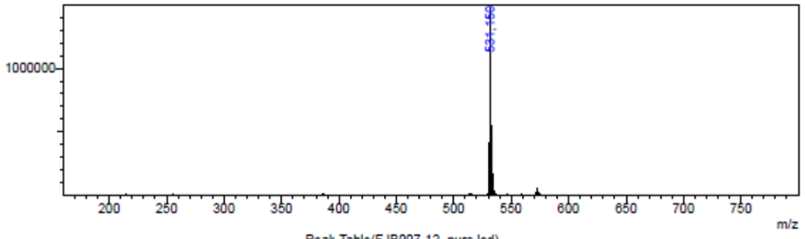

Peak Table(FJB007-12_pure.lcd)

Detector A Channel 2 254nm

| Peak# | Ret. Time | Area | Height | Mark | Conc. | Area% |
|---|---|---|---|---|---|---|
| 1 | 8,644 | 8159589 | 1447606 | M | 86,061 | 86,061 |
| 2 | 8,900 | 1321555 | 248919 | M | 13,939 | 13,939 |
| Total | | 9481144 | 1696526 | | 100,000 | 100,000 |

**24:** Exact mass: 492.25; observed $[M + H]^{+}$: 493.20

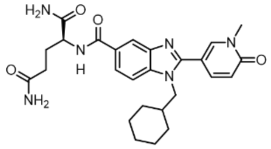

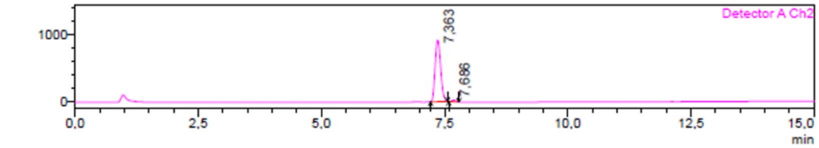

MS Spectrum(FJB007-2_pure.lcd)

Ret. Time: 1-1(E+) [7,243->7,548]
Inten.

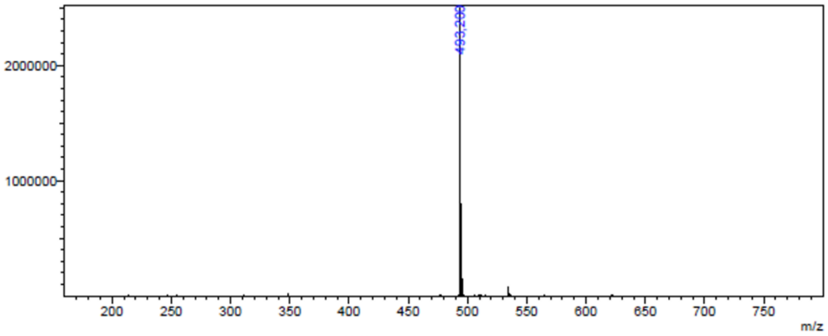

Peak Table(FJB007-2_pure.lcd)

Detector A Channel 2 254nm

| Peak# | Ret. Time | Area | Height | Mark | Conc. | Area% |
|---|---|---|---|---|---|---|
| 1 | 7,363 | 7196300 | 918091 | M | 98,048 | 98,048 |
| 2 | 7,686 | 143301 | 24727 | M | 1,952 | 1,952 |
| Total | | 7339601 | 942818 | | 100,000 | 100,000 |

**25:** Exact mass: 490.27; observed [M + H]+: 491.20

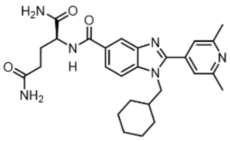

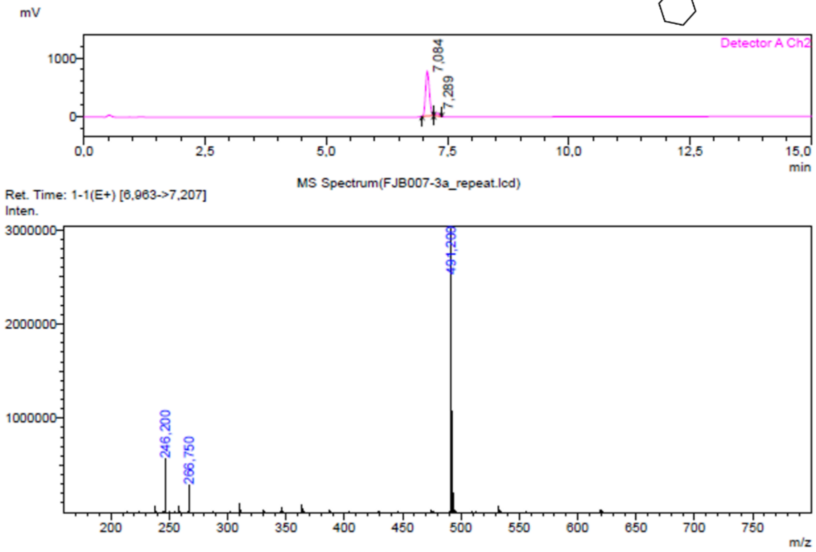

Peak Table(FJB007-3a_repeat.lcd)

Detector A Channel 2 254nm

| Peak# | Ret. Time | Area | Height | Mark | Conc. | Area% |
|---|---|---|---|---|---|---|
| 1 | 7,084 | 4157667 | 757188 | M | 94,754 | 94,754 |
| 2 | 7,289 | 230203 | 50180 | M | 5,246 | 5,246 |
| Total | | 4387870 | 807368 | | 100,000 | 100,000 |

**26:** Exact mass: 522.26; observed [M + H]+: 523.25

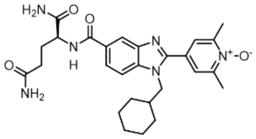

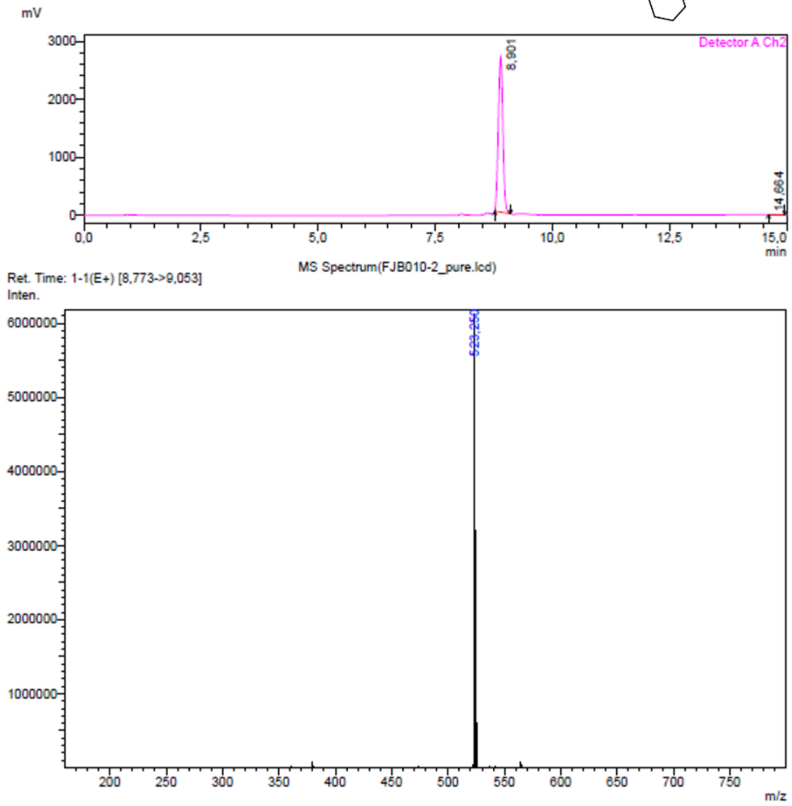

**27:** Exact mass: 502.27; observed [M + H]$^+$:503.25

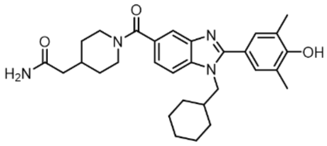

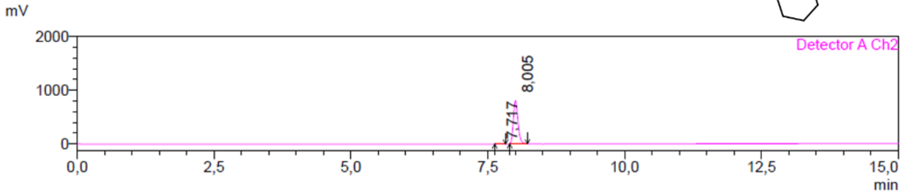

MS Spectrum(FJB003-7_pure.lcd)

Ret. Time: 1-1(E+) [7,898->8,188]
Inten.

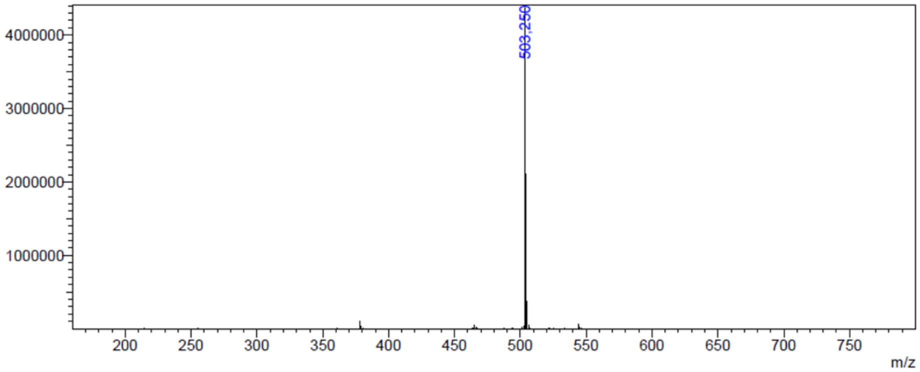

Peak Table(FJB003-7_pure.lcd)

Detector A Channel 2 254nm

| Peak# | Ret. Time | Area | Height | Mark | Conc. | Area% |
|---|---|---|---|---|---|---|
| 1 | 7,717 | 35392 | 6203 | M | 0,748 | 0,748 |
| 2 | 8,005 | 4696236 | 799747 | M | 99,252 | 99,252 |
| Total | | 4731627 | 805950 | | 100,000 | 100,000 |

**28:** Exact mass: 448.25; observed [M + H]$^+$: 449.15

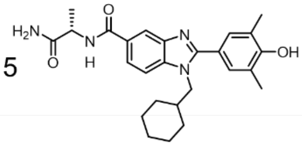

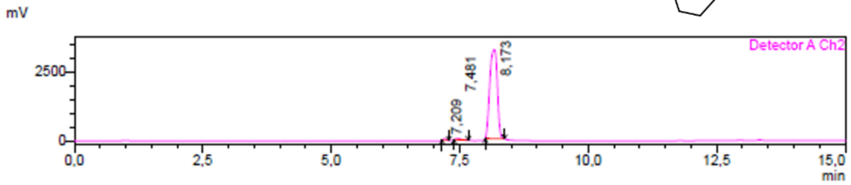

MS Spectrum(FJB010-1_pure.lcd)

Ret. Time: 1-1(E+) [8,012->8,340]
Inten.

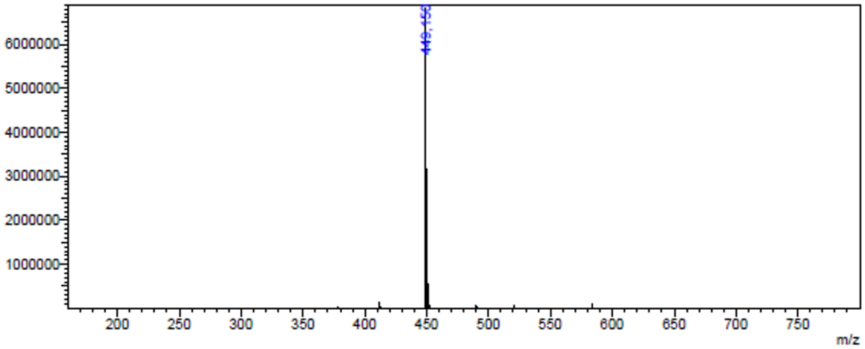

Peak Table(FJB010-1_pure.lcd)

Detector A Channel 2 254nm

| Peak# | Ret. Time | Area | Height | Mark | Conc. | Area% |
|---|---|---|---|---|---|---|
| 1 | 7,209 | 215608 | 49997 | M | 0,646 | 0,646 |
| 2 | 7,481 | 280155 | 44041 | M | 0,839 | 0,839 |
| 3 | 8,173 | 32895102 | 3216991 | M | 98,515 | 98,515 |
| Total | | 33390865 | 3311030 | | 100,000 | 100,000 |

**25:** Exact mass: 490.27; observed [M + H]$^+$: 491.20

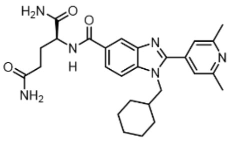

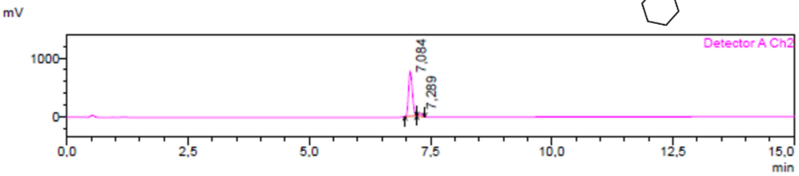

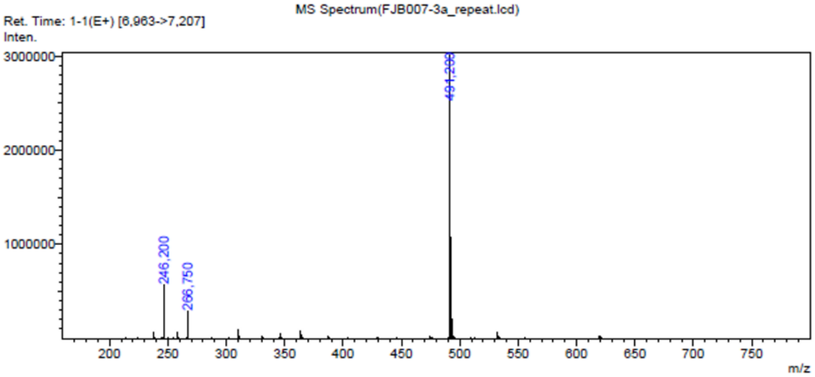

Peak Table(FJB007-3a_repeat.lcd)

Detector A Channel 2 254nm

| Peak# | Ret. Time | Area | Height | Mark | Conc. | Area% |
|---|---|---|---|---|---|---|
| 1 | 7,084 | 4157667 | 757188 | M | 94,754 | 94,754 |
| 2 | 7,289 | 230203 | 50180 | M | 5,246 | 5,246 |
| Total | | 4387870 | 807368 | | 100,000 | 100,000 |

**26:** Exact mass: 522.26; observed [M + H]$^+$: 523.25

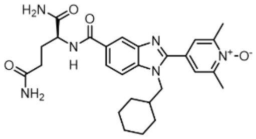

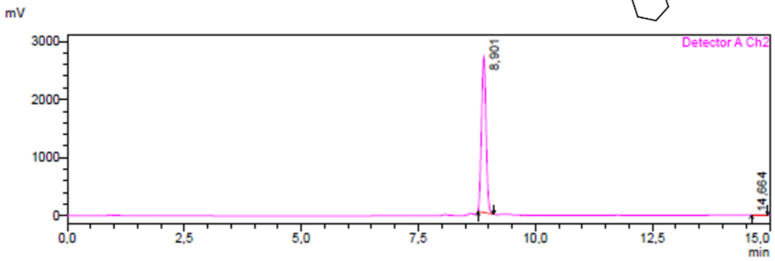

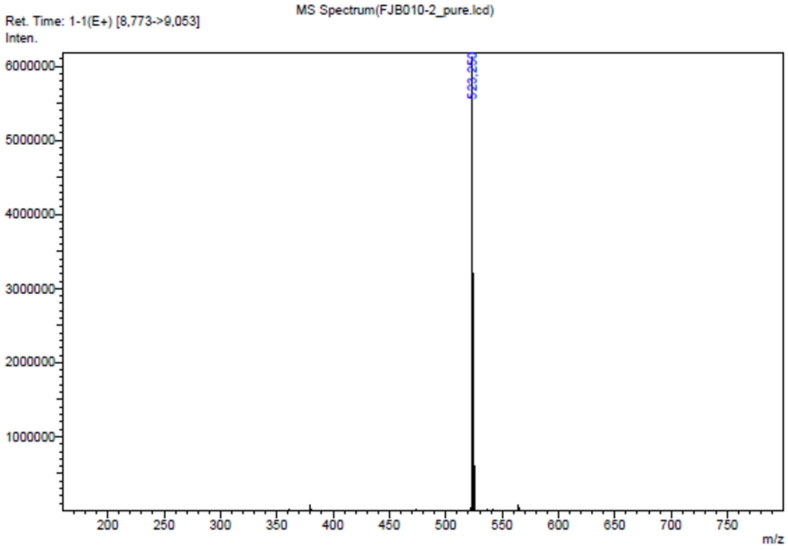

**27:** Exact mass: 502.27; observed [M + H]$^+$:503.25

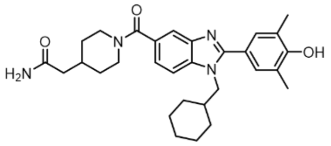

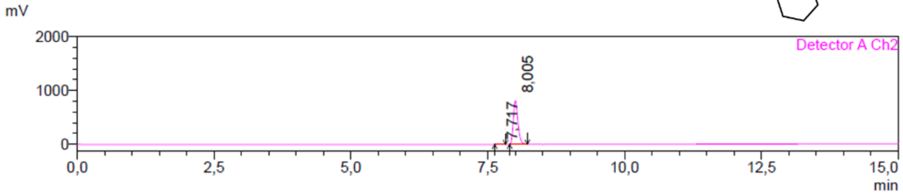

MS Spectrum(FJB003-7_pure.lcd)

Ret. Time: 1-1(E+) [7,898->8,188]
Inten.

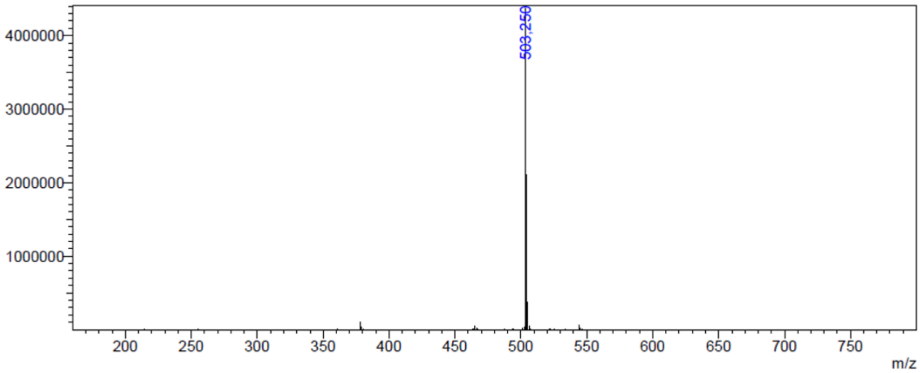

Peak Table(FJB003-7_pure.lcd)

Detector A Channel 2 254nm

| Peak# | Ret. Time | Area | Height | Mark | Conc. | Area% |
|---|---|---|---|---|---|---|
| 1 | 7,717 | 35392 | 6203 | M | 0,748 | 0,748 |
| 2 | 8,005 | 4696236 | 799747 | M | 99,252 | 99,252 |
| Total | | 4731627 | 805950 | | 100,000 | 100,000 |

**28:** Exact mass: 448.25; observed [M + H]$^+$: 449.15

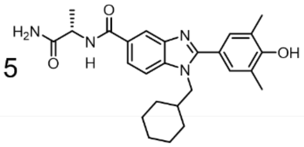

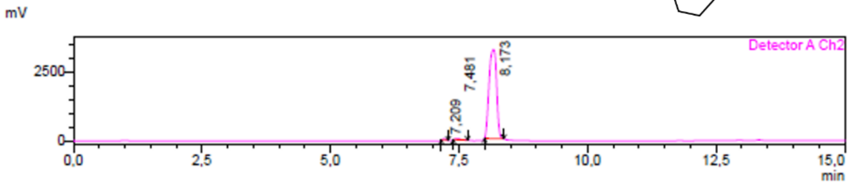

MS Spectrum(FJB010-1_pure.lcd)

Ret. Time: 1-1(E+) [8,012->8,340]
Inten.

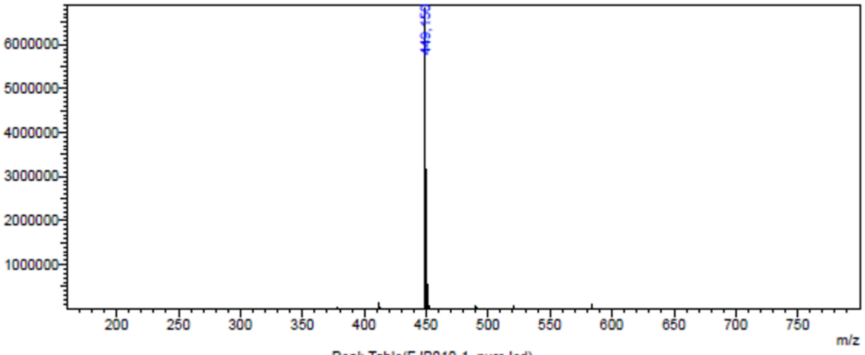

Peak Table(FJB010-1_pure.lcd)

Detector A Channel 2 254nm

| Peak# | Ret. Time | Area | Height | Mark | Conc. | Area% |
|---|---|---|---|---|---|---|
| 1 | 7,209 | 215608 | 49997 | M | 0,646 | 0,646 |
| 2 | 7,481 | 280155 | 44041 | M | 0,839 | 0,839 |
| 3 | 8,173 | 32895102 | 3216991 | M | 98,515 | 98,515 |
| Total | | 33390865 | 3311030 | | 100,000 | 100,000 |

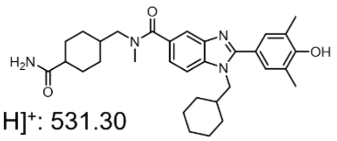

**29:** Exact mass: 530.33; observed $[M + H]^+$: 531.30

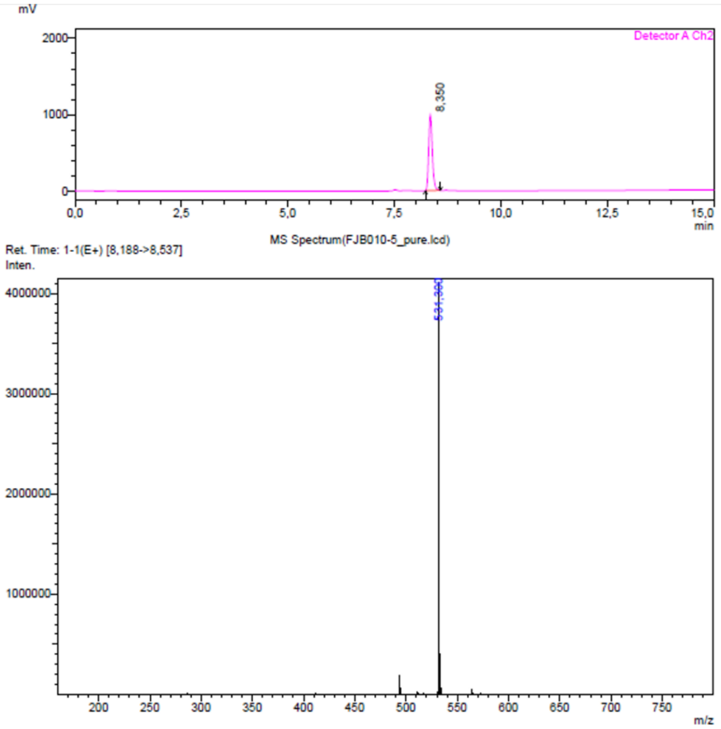

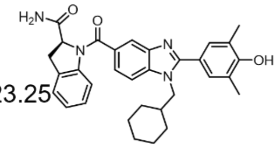

**30:** Exact mass: 522.26; observed $[M + H]^+$: 523.25

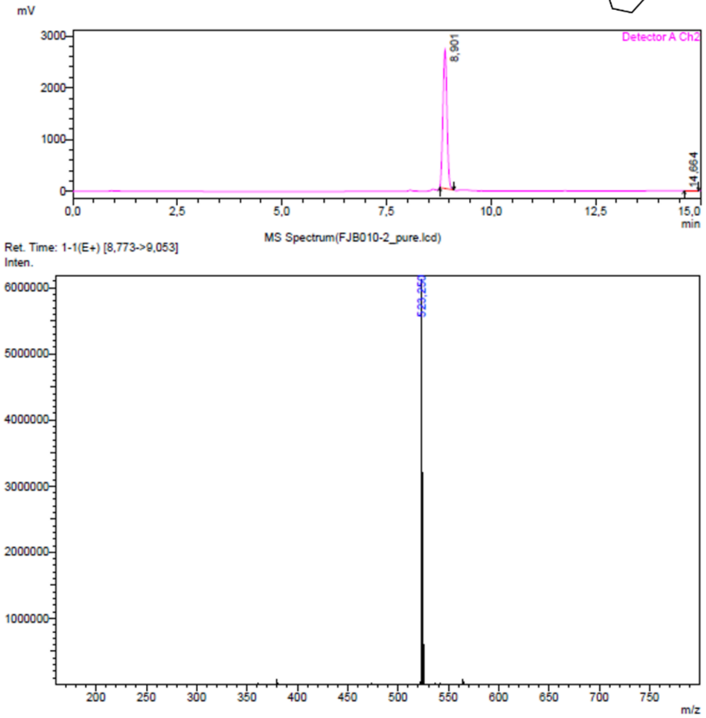

**31:** Exact mass: 490.26; observed [M + H]$^+$: 491.20

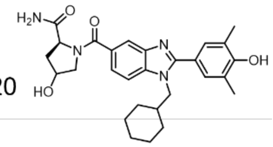

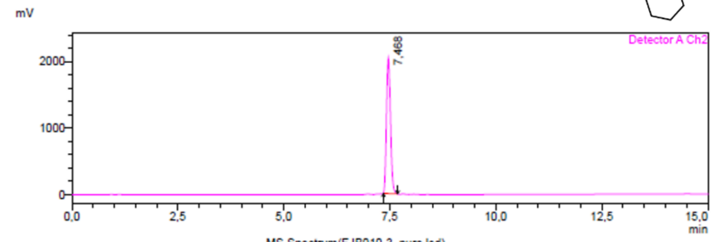

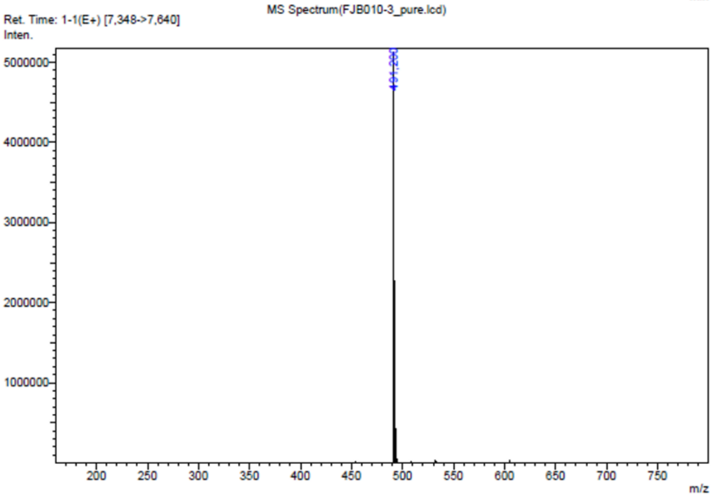

**32:** Exact mass: 510.24; observed [M + H]$^+$: 511.25

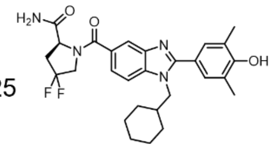

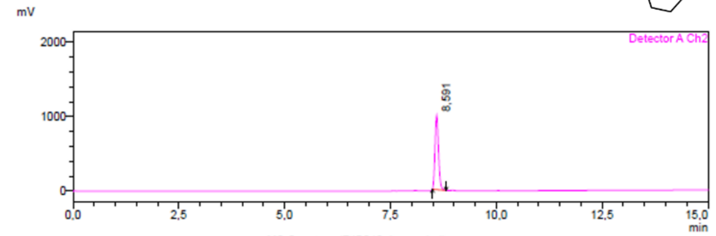

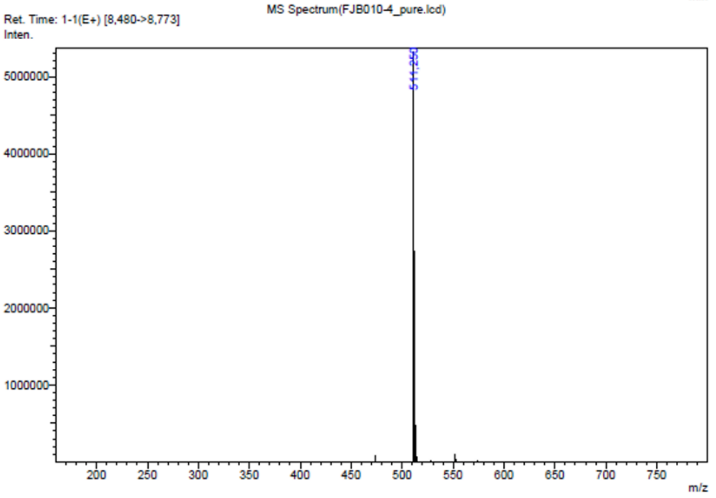

**33:** Exact mass: 595.32; observed $[M + H]^+$: 596.30

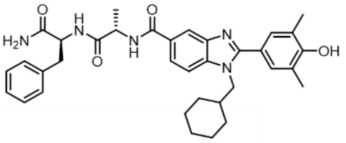

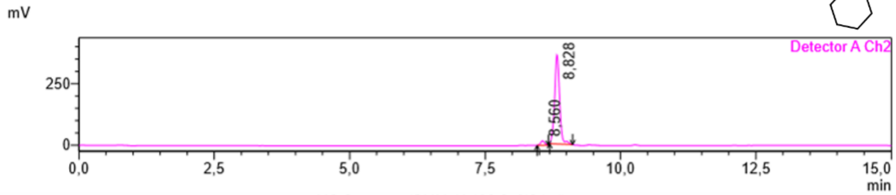

MS Spectrum(SMMcK-196-2_f12.lcd)

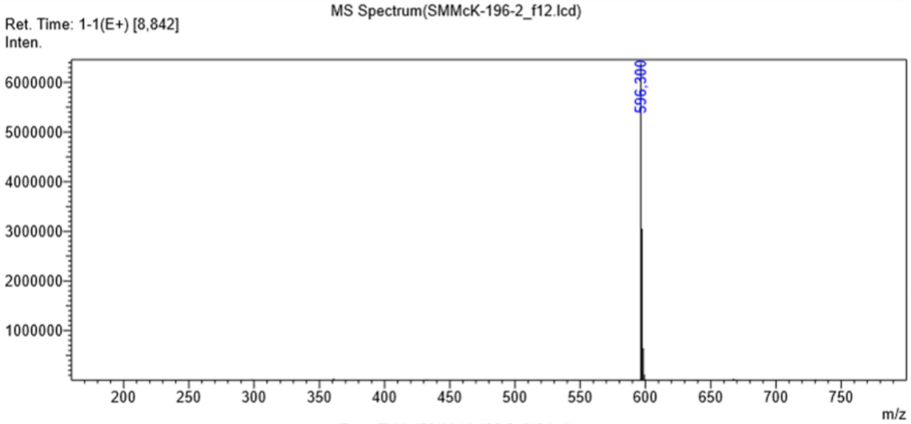

Peak Table(SMMcK-196-2_f12.lcd)

Detector A Channel 1 254nm

| Peak# | Ret. Time | Area | Height | Mark | Conc. | Area% |
|---|---|---|---|---|---|---|
| 1 | 8.558 | 73877 | 15380 | M | 3.610 | 3.610 |
| 2 | 8.830 | 1967841 | 355386 | M | 96.167 | 96.167 |
| 3 | 9.003 | 4561 | 2665 | M | 0.223 | 0.223 |
| Total | | 2046280 | 373430 | | 100.000 | 100.000 |

**34:** Exact mass: 720.40; observed $[M + H]^+$: 721.40

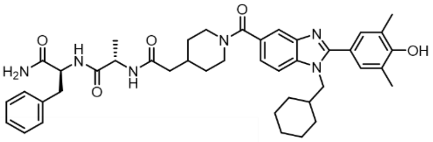

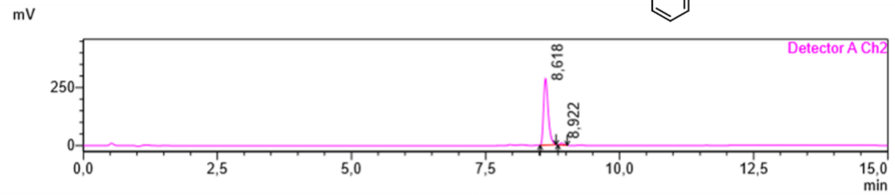

MS Spectrum(SMMcK-197-O-f16-18.lcd)

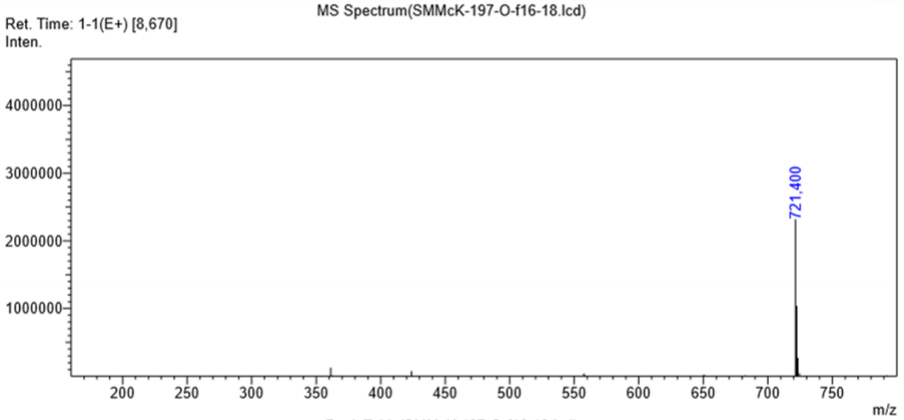

Peak Table(SMMcK-197-O-f16-18.lcd)

Detector A Channel 2 275nm

| Peak# | Ret. Time | Area | Height | Mark | Conc. | Area% |
|---|---|---|---|---|---|---|
| 1 | 8.618 | 1721640 | 286877 | M | 97.670 | 97.670 |
| 2 | 8.922 | 41070 | 8235 | M | 2.330 | 2.330 |
| Total | | 1762710 | 295112 | | 100.000 | 100.000 |

**35:** Exact mass: 740.39; observed $[M + H]^+$: 741.35

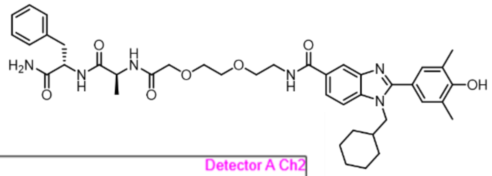

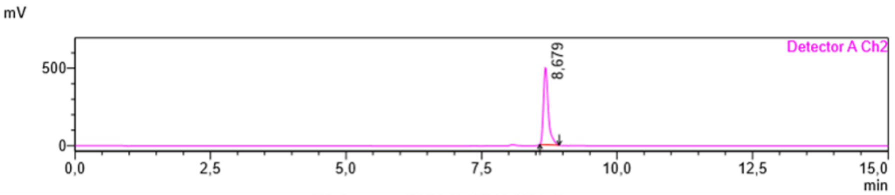

MS Spectrum(SMMcK-197-N_f10.lcd)

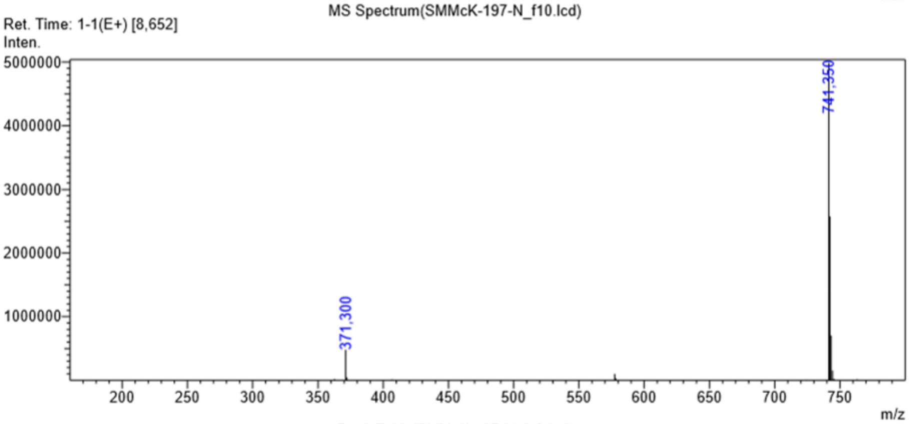

Peak Table(SMMcK-197-N_f10.lcd)

Detector A Channel 1 254nm

| Peak# | Ret. Time | Area | Height | Mark | Conc. | Area% |
|---|---|---|---|---|---|---|
| 1 | 8.678 | 3423222 | 582153 | M | 100.000 | 100.000 |
| Total | | 3423222 | 582153 | | 100.000 | 100.000 |

**36:** Exact mass: 721.40; observed $[M + H]^+$: 722.40

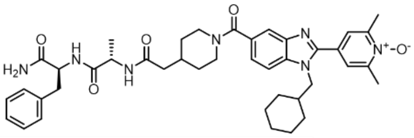

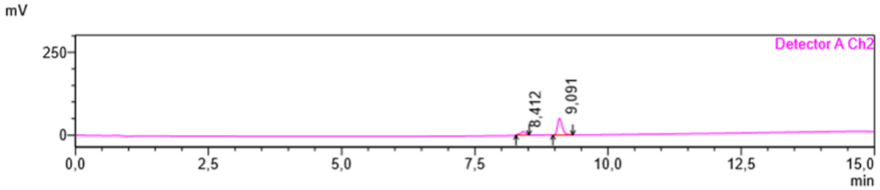

MS Spectrum(SMMcK-197-B_N-ox_rep_f6.lcd)

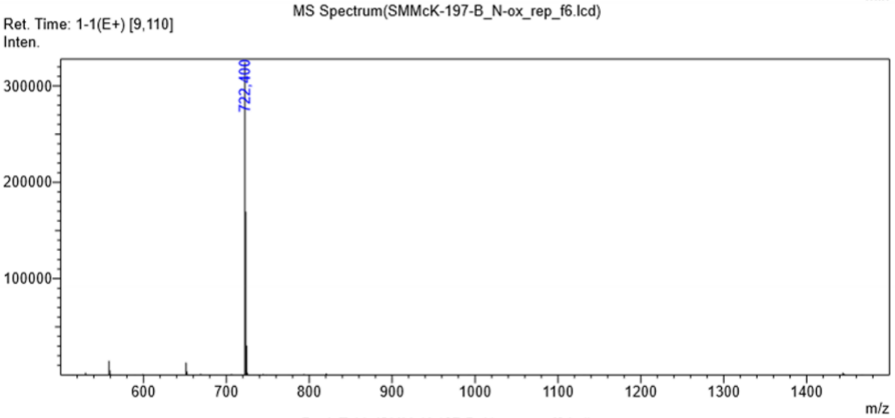

Peak Table(SMMcK-197-B_N-ox_rep_f6.lcd)

Detector A Channel 2 254nm

| Peak# | Ret. Time | Area | Height | Mark | Conc. | Area% |
|---|---|---|---|---|---|---|
| 1 | 8.412 | 70882 | 9720 | M | 18.419 | 18.419 |
| 2 | 9.091 | 313946 | 50041 | M | 81.581 | 81.581 |
| Total | | 384828 | 59761 | | 100.000 | 100.000 |

**37:** Exact mass: 833.48; observed $[M + H]^+$: 834.50

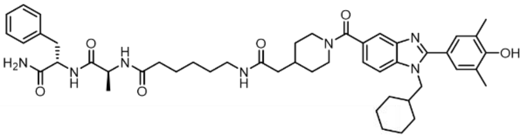

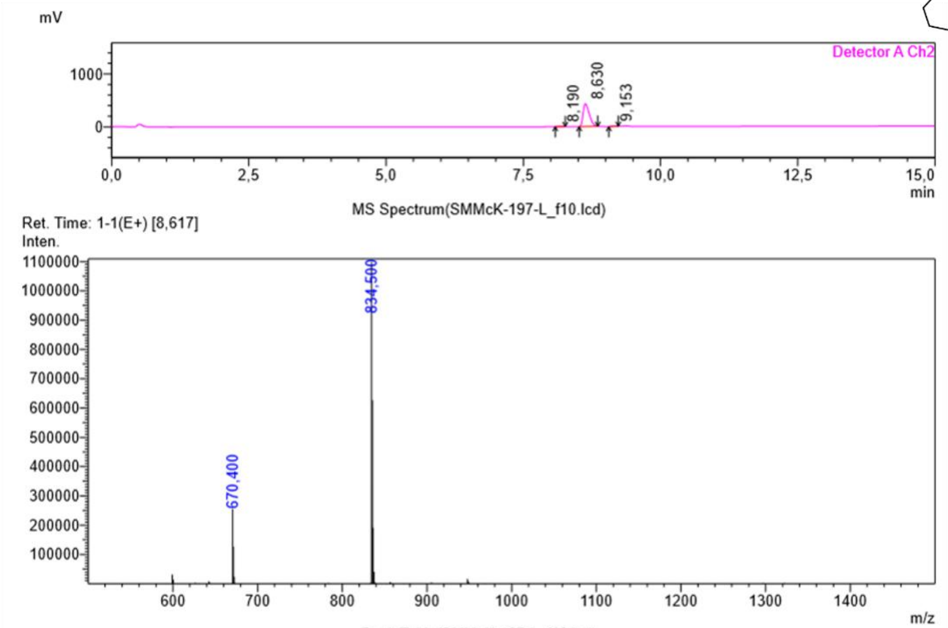

Peak Table(SMMcK-197-L_f10.lcd)

Detector A Channel 2 254nm

| Peak# | Ret. Time | Area | Height | Mark | Conc. | Area% |
|---|---|---|---|---|---|---|
| 1 | 8.190 | 21480 | 3313 | M | 0.623 | 0.623 |
| 2 | 8.630 | 3373939 | 431539 | M | 97.850 | 97.850 |
| 3 | 9.153 | 52643 | 10926 | M | 1.527 | 1.527 |
| Total | | 3448062 | 445778 | | 100.000 | 100.000 |

**38:** Exact mass: 821.48; observed $[M + H]^+$: 721.40

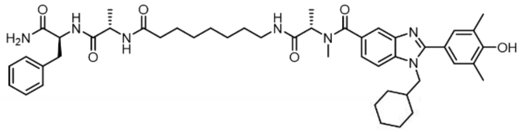

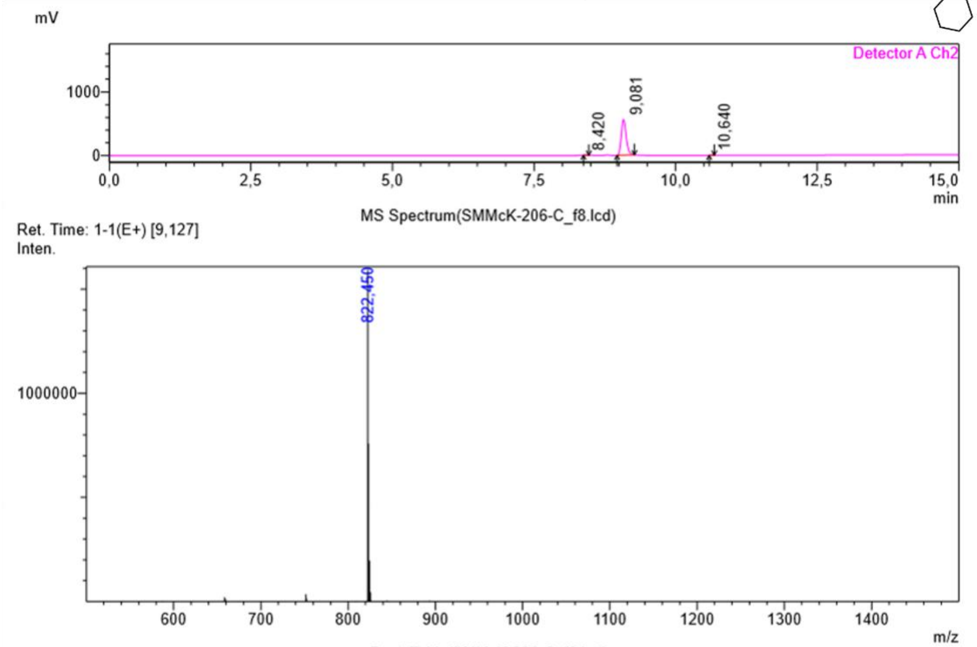

Peak Table(SMMcK-206-C_f8.lcd)

Detector A Channel 2 254nm

| Peak# | Ret. Time | Area | Height | Mark | Conc. | Area% |
|---|---|---|---|---|---|---|
| 1 | 8.420 | 5888 | 1704 | M | 0.172 | 0.172 |
| 2 | 9.081 | 3414954 | 560445 | M | 99.589 | 99.589 |
| 3 | 10.640 | 8215 | 2362 | M | 0.240 | 0.240 |
| Total | | 3429057 | 564511 | | 100.000 | 100.000 |

**39:** Exact mass: 861.52; observed [M + H]+: 862.55

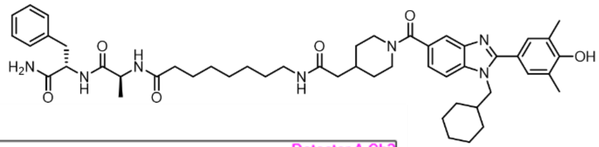

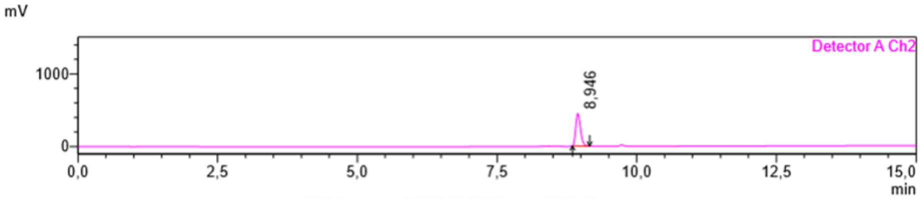

MS Spectrum(SMMcK-197-k_rep_f16.lcd)

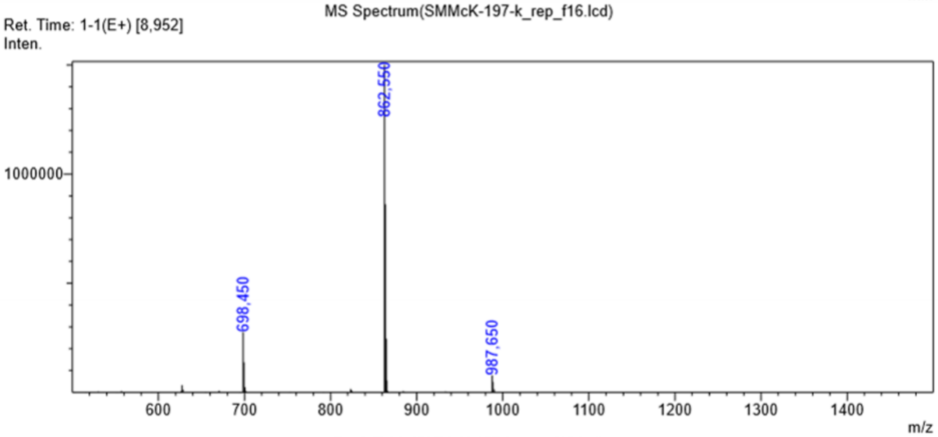

Peak Table(SMMcK-197-k_rep_f16.lcd)

Detector A Channel 1 214nm

| Peak# | Ret. Time | Area | Height | Mark | Conc. | Area% |
|---|---|---|---|---|---|---|
| 1 | 8.948 | 7454669 | 1165909 | M | 100.000 | 100.000 |
| Total | | 7454669 | 1165909 | | 100.000 | 100.000 |

**40:** Exact mass: 865.47; observed [M + H]+: 866.45

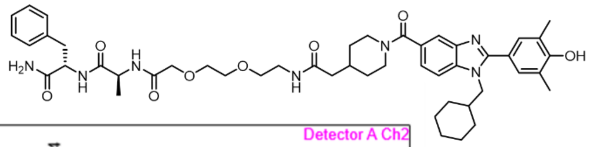

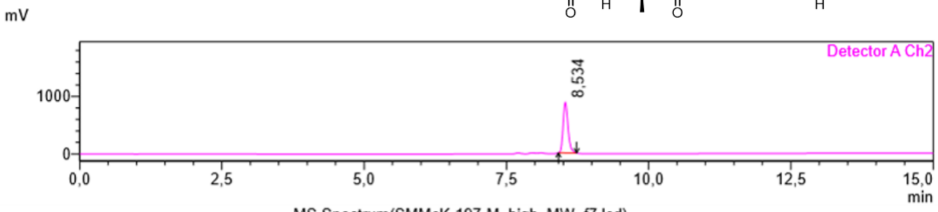

MS Spectrum(SMMcK-197-M_high_MW_f7.lcd)

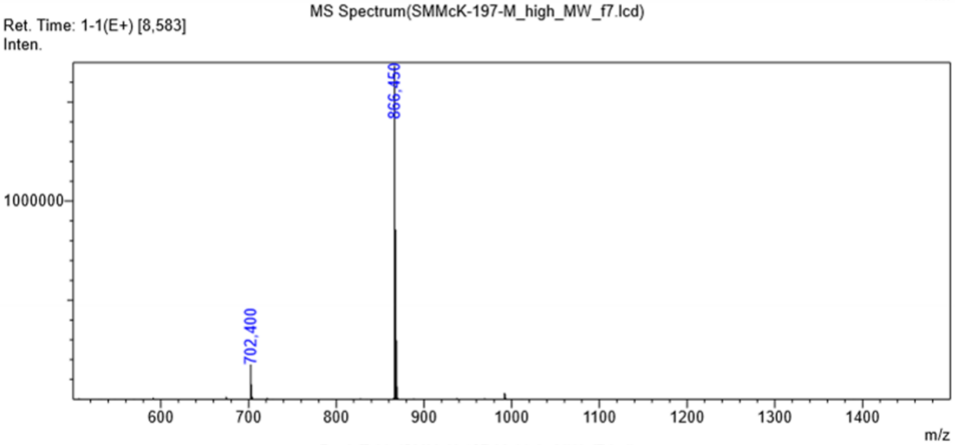

Peak Table(SMMcK-197-M_high_MW_f7.lcd)

Detector A Channel 2 254nm

| Peak# | Ret. Time | Area | Height | Mark | Conc. | Area% |
|---|---|---|---|---|---|---|
| 1 | 8.534 | 5294250 | 881325 | M | 100.000 | 100.000 |
| Total | | 5294250 | 881325 | | 100.000 | 100.000 |

**41:** Exact mass: 825.44; observed $[M + H]^{+}$: 826.45

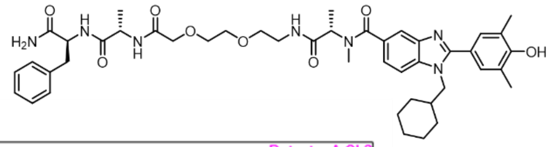

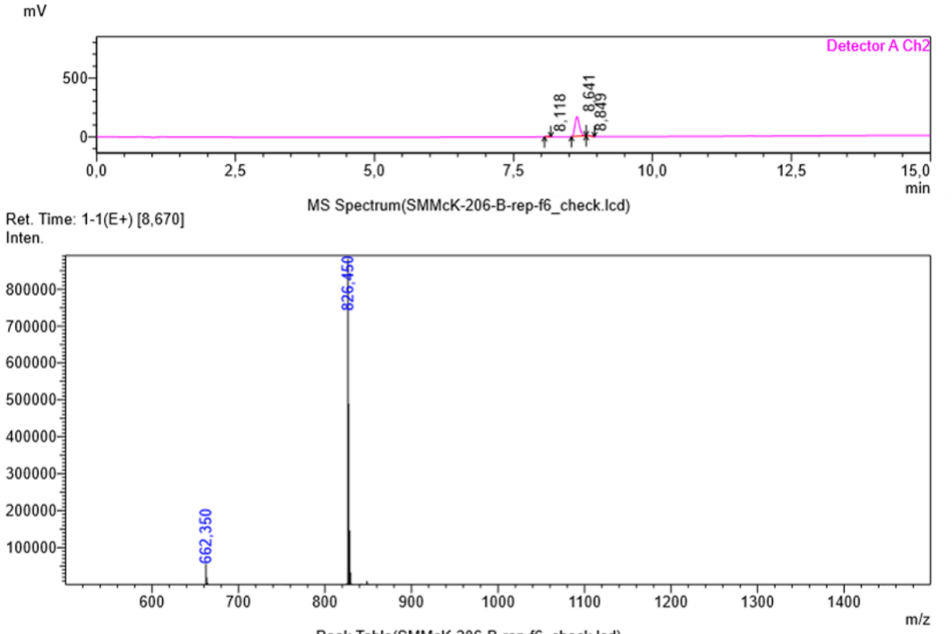

Peak Table(SMMcK-206-B-rep-f6_check.lcd)

Detector A Channel 2 254nm

| Peak# | Ret. Time | Area | Height | Mark | Conc. | Area% |
|---|---|---|---|---|---|---|
| 1 | 8.118 | 12291 | 2896 | M | 1.243 | 1.243 |
| 2 | 8.641 | 965338 | 168807 | M | 97.586 | 97.586 |
| 3 | 8.849 | 11592 | 2964 | M | 1.172 | 1.172 |
| Total | | 989221 | 174667 | | 100.000 | 100.000 |

**42:** Exact mass: 864.49; observed $[M + H]^{+}$: 865.50

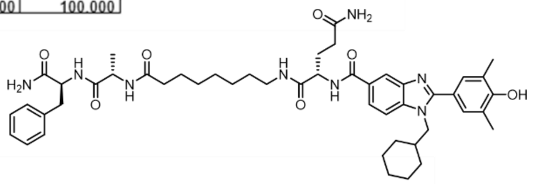

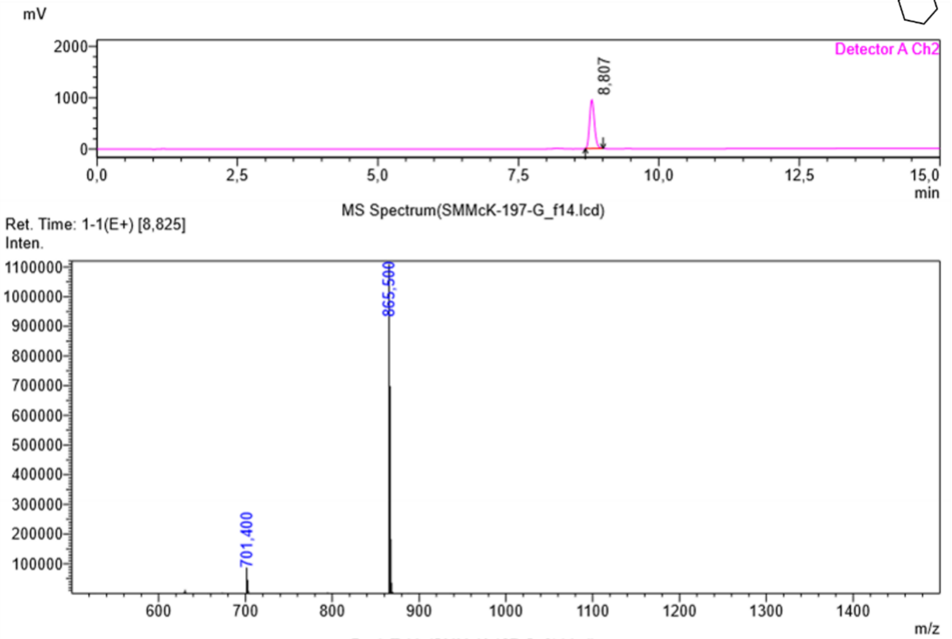

Peak Table(SMMcK-197-G_f14.lcd)

Detector A Channel 2 254nm

| Peak# | Ret. Time | Area | Height | Mark | Conc. | Area% |
|---|---|---|---|---|---|---|
| 1 | 8.807 | 5689227 | 950112 | M | 100.000 | 100.000 |
| Total | | 5689227 | 950112 | | 100.000 | 100.000 |

**43:** Exact mass: 777.97; observed $[M + H]^+$: 778.40

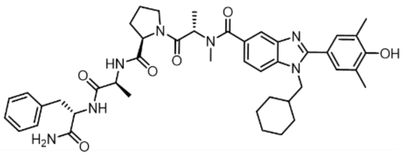

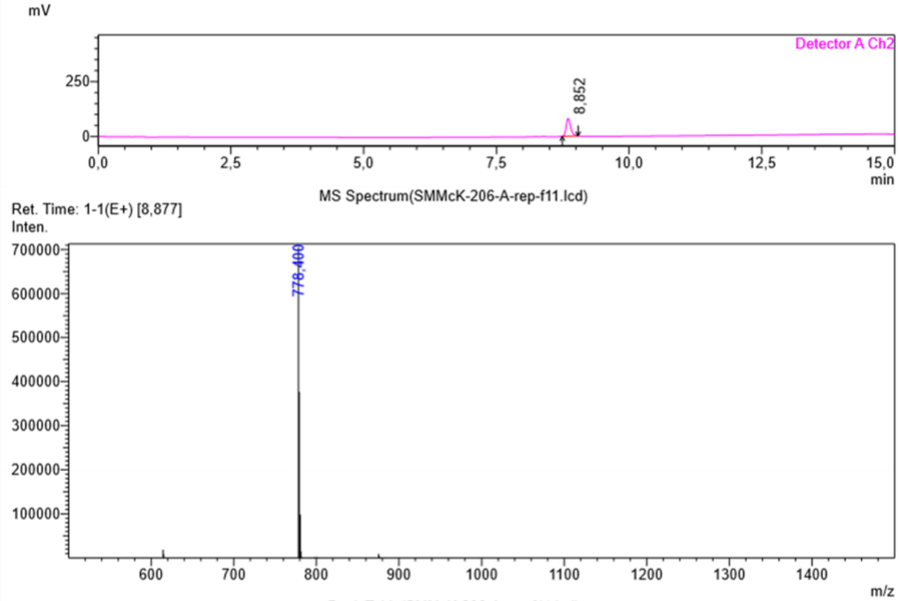

Peak Table(SMMcK-206-A-rep-f11.lcd)

Detector A Channel 2 254nm

| Peak# | Ret. Time | Area | Height | Mark | Conc. | Area% |
|---|---|---|---|---|---|---|
| 1 | 8.852 | 478501 | 79669 | M | 100.000 | 100.000 |
| Total | | 478501 | 79669 | | 100.000 | 100.000 |

**44:** Exact mass: 962.60; observed $[M + H]^+$: 963.55

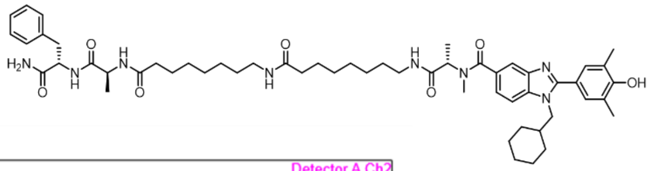

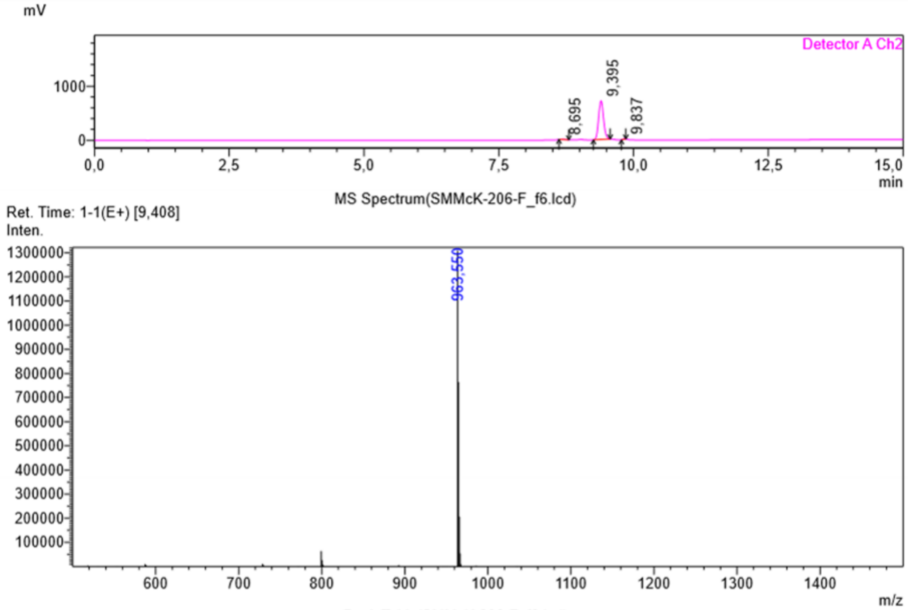

Peak Table(SMMcK-206-F_f6.lcd)

Detector A Channel 2 254nm

| Peak# | Ret. Time | Area | Height | Mark | Conc. | Area% |
|---|---|---|---|---|---|---|
| 1 | 8.695 | 44821 | 7645 | M | 0.993 | 0.993 |
| 2 | 9.395 | 4442784 | 716187 | M | 98.417 | 98.417 |
| 3 | 9.837 | 26654 | 6839 | M | 0.590 | 0.590 |
| Total | | 4514259 | 730671 | | 100.000 | 100.000 |

**45:** Exact mass: 970.52; observed $[M + H]^+$: 971.50

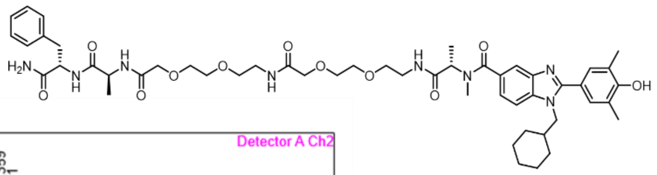

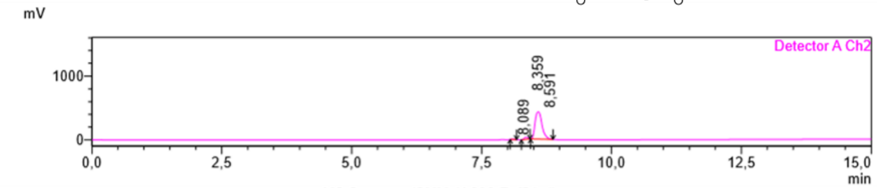

MS Spectrum(SMMcK-206-E_f5.lcd)

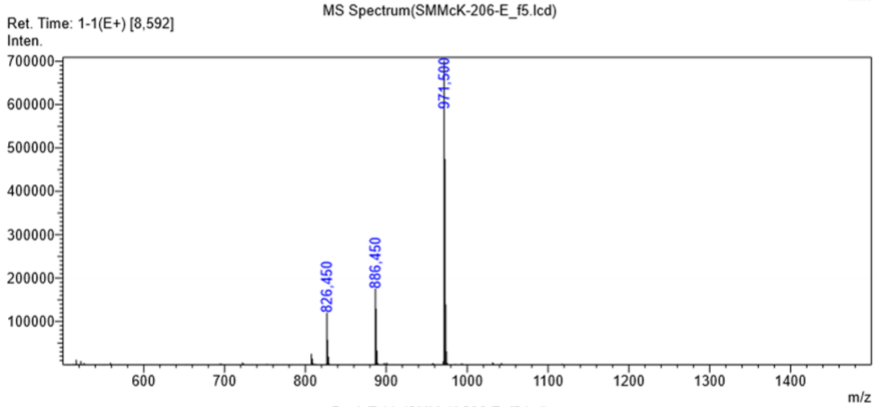

Peak Table(SMMcK-206-E_f5.lcd)

Detector A Channel 2 254nm

| Peak# | Ret. Time | Area | Height | Mark | Conc. | Area% |
|---|---|---|---|---|---|---|
| 1 | 8.089 | 23259 | 5796 | M | 0.557 | 0.557 |
| 2 | 8.359 | 158512 | 33602 | M | 3.798 | 3.798 |
| 3 | 8.591 | 3991757 | 425823 | M | 95.645 | 95.645 |
| Total | | 4173528 | 465222 | | 100.000 | 100.000 |

**46:** Exact mass: 834.44; observed $[M + H]^+$: 835.45

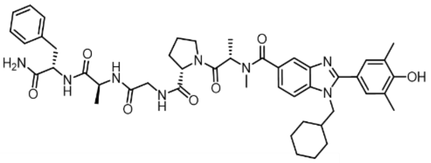

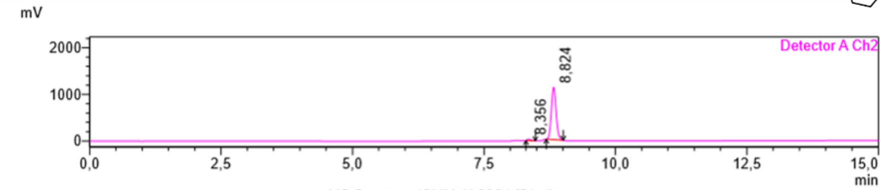

MS Spectrum(SMMcK-206d-f5.lcd)

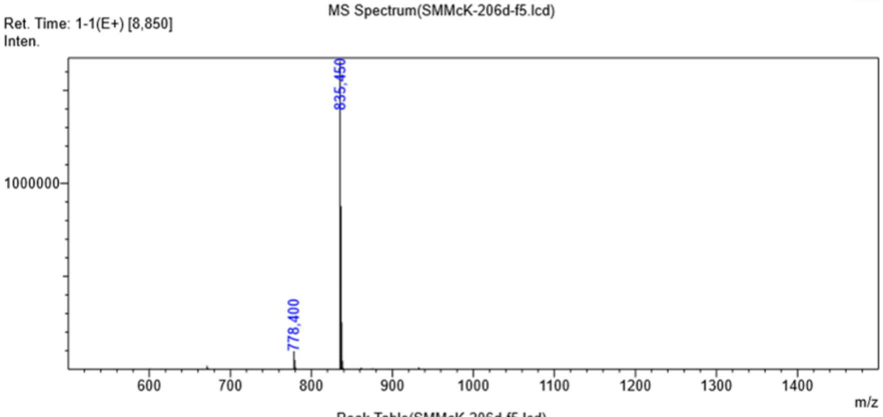

Peak Table(SMMcK-206d-f5.lcd)

Detector A Channel 2 254nm

| Peak# | Ret. Time | Area | Height | Mark | Conc. | Area% |
|---|---|---|---|---|---|---|
| 1 | 8.356 | 151033 | 32760 | M | 2.093 | 2.093 |
| 2 | 8.824 | 7065496 | 1122429 | M | 97.907 | 97.907 |
| Total | | 7216529 | 1155189 | | 100.000 | 100.000 |

**47:** Exact mass: 1008.54; observed $[M + H]^+$: 1009.55

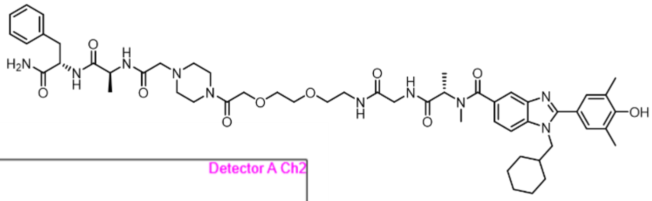

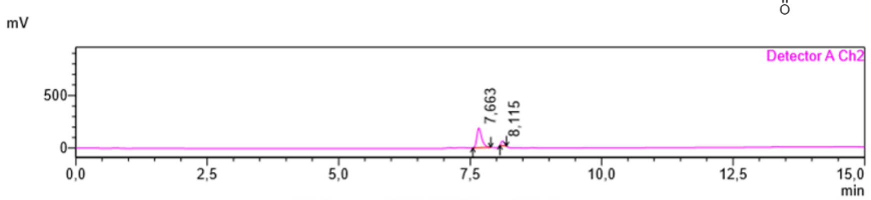

MS Spectrum(SMMcK-197-f-rep_f12.lcd)

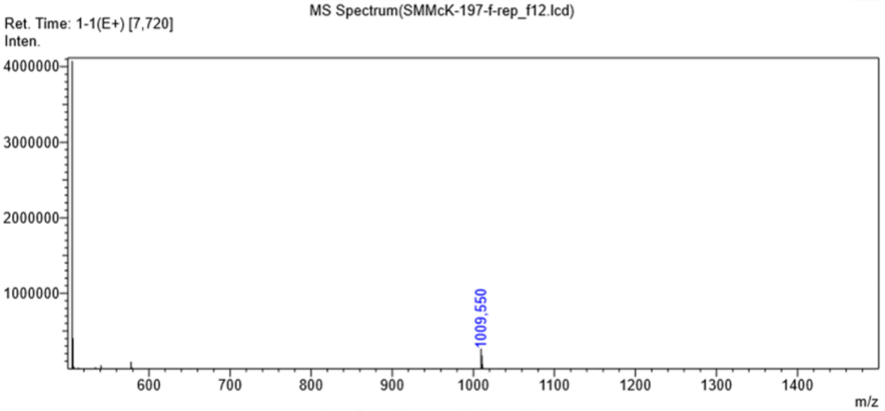

Peak Table(SMMcK-197-f-rep_f12.lcd)

Detector A Channel 2 254nm

| Peak# | Ret. Time | Area | Height | Mark | Conc. | Area% |
|---|---|---|---|---|---|---|
| 1 | 7.663 | 1204930 | 184828 | M | 86.914 | 86.914 |
| 2 | 8.115 | 181418 | 43099 | M | 13.086 | 13.086 |
| Total | | 1386347 | 227927 | | 100.000 | 100.000 |

**48:** Exact mass: 846.52; observed $[M + H]^+$: 847.50

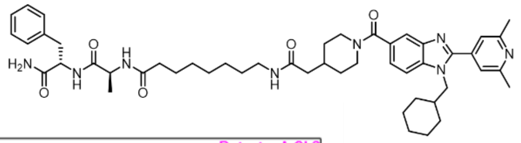

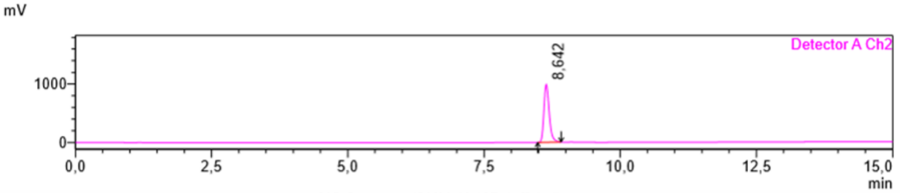

MS Spectrum(SMMcK-197-J_f2.lcd)

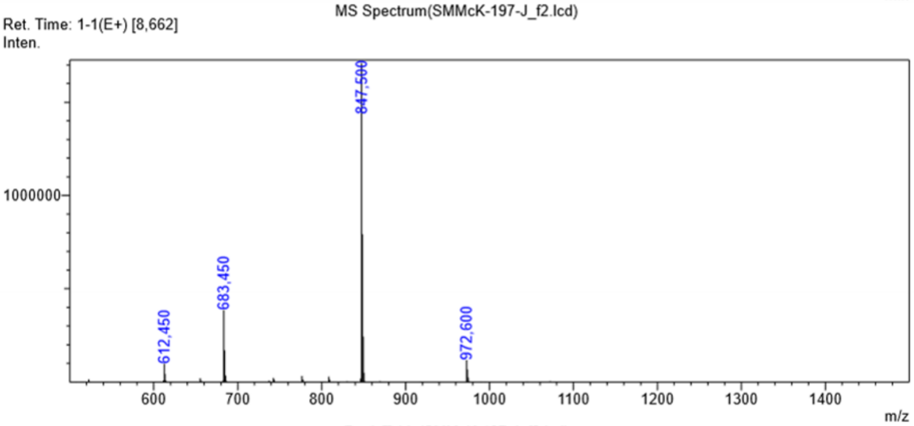

Peak Table(SMMcK-197-J_f2.lcd)

Detector A Channel 1 214nm

| Peak# | Ret. Time | Area | Height | Mark | Conc. | Area% |
|---|---|---|---|---|---|---|
| 1 | 8.645 | 13336751 | 1428029 | M | 100.000 | 100.000 |
| Total | | 13336751 | 1428029 | | 100.000 | 100.000 |

**49:** Exact mass: 862.51; observed $[M + H]^+$: 863.55

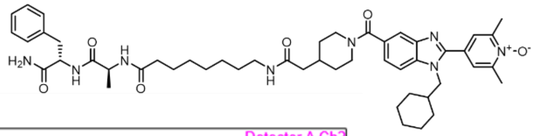

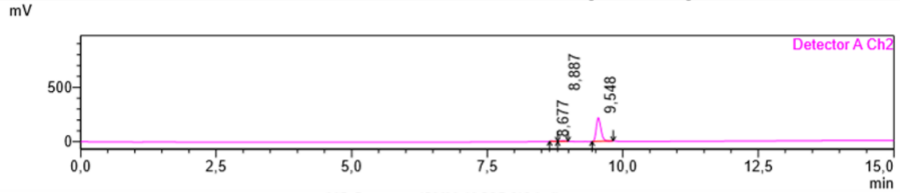

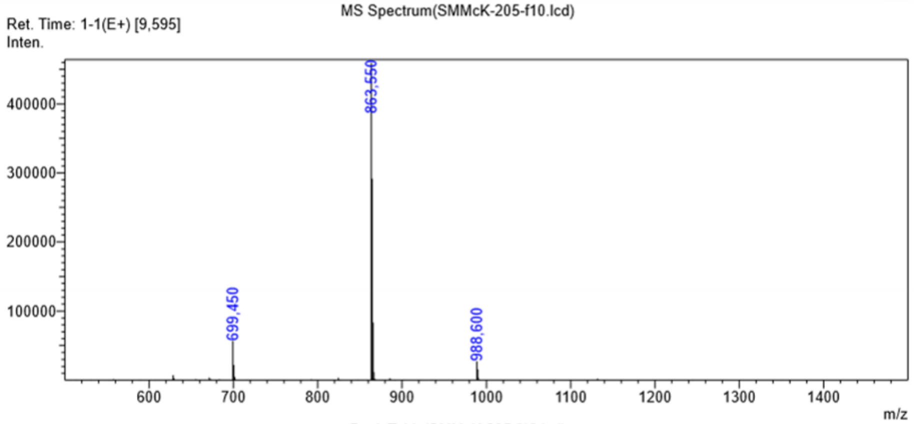

Peak Table(SMMcK-205-f10.lcd)

Detector A Channel 2 254nm

| Peak# | Ret. Time | Area | Height | Mark | Conc. | Area% |
|---|---|---|---|---|---|---|
| 1 | 8.677 | 5488 | 1007 | M | 0.416 | 0.416 |
| 2 | 8.887 | 28072 | 5250 | M | 2.130 | 2.130 |
| 3 | 9.548 | 1284114 | 216681 | M | 97.453 | 97.453 |
| Total | | 1317673 | 222938 | | 100.000 | 100.000 |

**50:** Exact mass: 639.31; observed $[M + H]^+$:

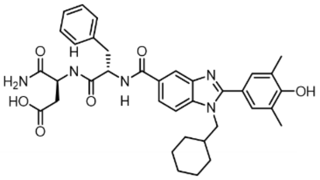

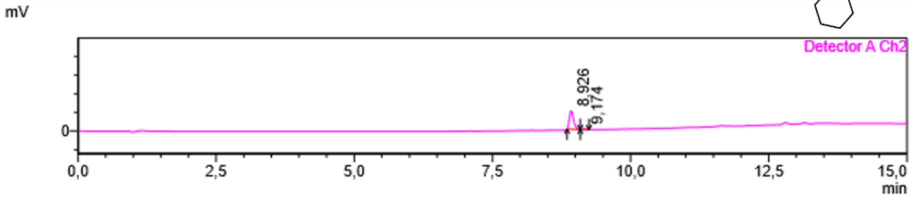

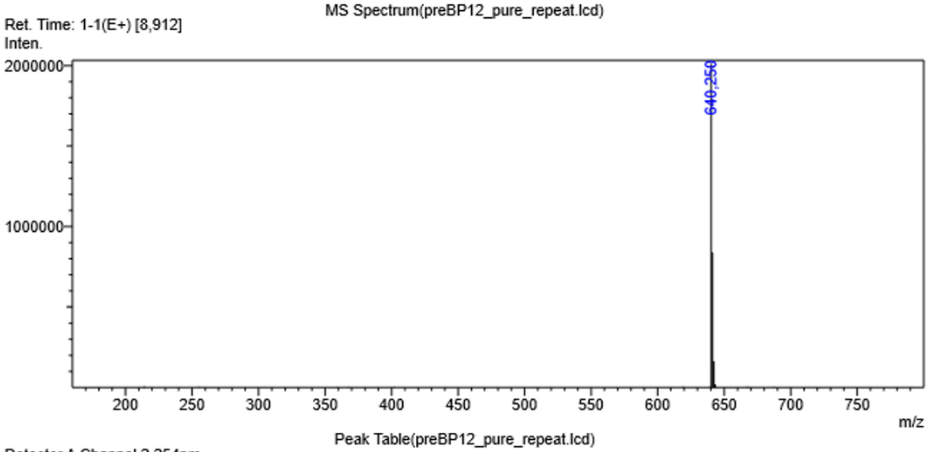

Peak Table(preBP12_pure_repeat.lcd)

Detector A Channel 2 254nm

| Peak# | Ret. Time | Area | Height | Mark | Conc. | Area% |
|---|---|---|---|---|---|---|
| 1 | 8.926 | 265861 | 49142 | M | 97.077 | 97.077 |
| 2 | 9.174 | 8004 | 1779 | M | 2.923 | 2.923 |
| Total | | 273865 | 50921 | | 100.000 | 100.000 |

**51:** Exact mass: 621.30; observed $[M + H]^{+}$: 622.30

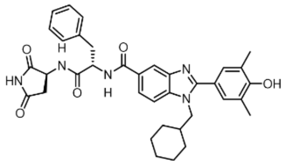

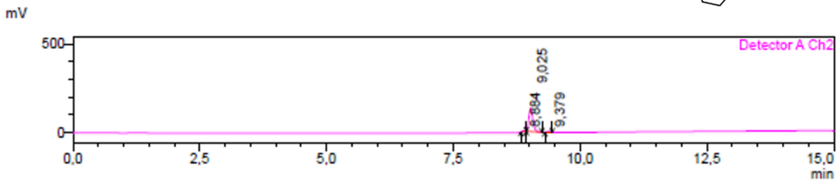

MS Spectrum(FJB032_BP12a_pure.lcd)

Ret. Time: 1-1(E+) [8,963->9,125]

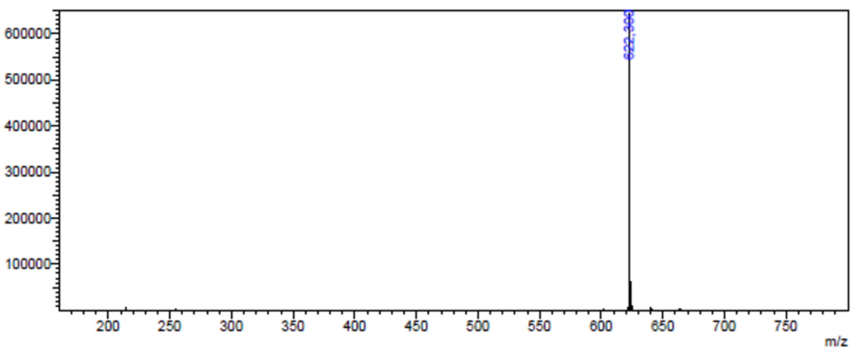

Peak Table(FJB032_BP12a_pure.lcd)

Detector A Channel 2 254nm

| Peak# | Ret. Time | Area | Height | Mark | Conc. | Area% |
|---|---|---|---|---|---|---|
| 1 | 8,884 | 14685 | 4366 | M | 1,934 | 1,934 |
| 2 | 9,025 | 725640 | 130135 | M | 95,552 | 95,552 |
| 3 | 9,379 | 19092 | 4115 | M | 2,514 | 2,514 |
| Total | | 759418 | 138616 | | 100,000 | 100,000 |

**52:** Exact mass: 474.23; observed $[M + H]^{+}$: 475.15

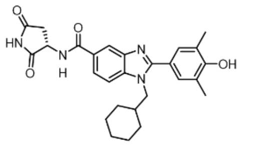

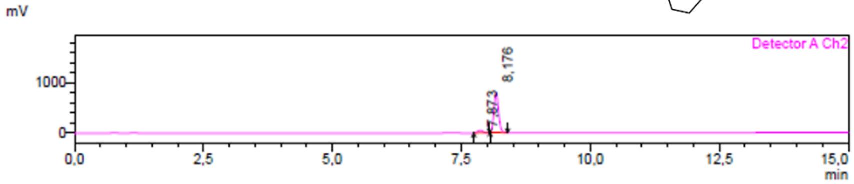

MS Spectrum(FJB026_BP16a_pure.lcd)

Ret. Time: 1-1(E+) [8,072->8,318]

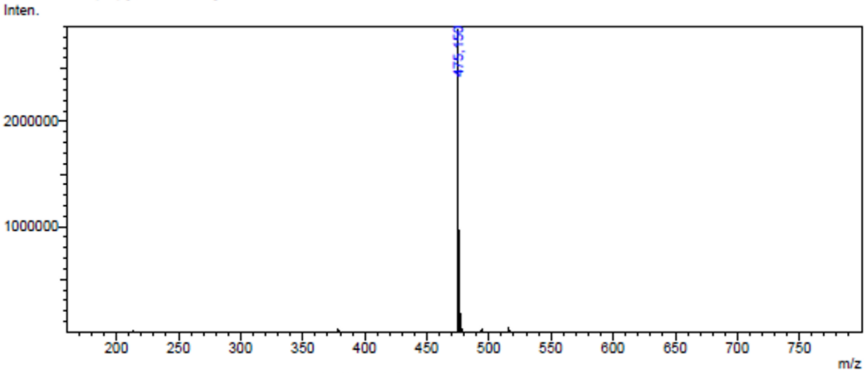

Peak Table(FJB026_BP16a_pure.lcd)

Detector A Channel 2 254nm

| Peak# | Ret. Time | Area | Height | Mark | Conc. | Area% |
|---|---|---|---|---|---|---|
| 1 | 7,873 | 356751 | 49761 | M | 6,673 | 6,673 |
| 2 | 8,176 | 4989226 | 790455 | M | 93,327 | 93,327 |
| Total | | 5345977 | 840216 | | 100,000 | 100,000 |

**53:** Exact mass: 635.31; observed $[M + H]^+$: 636.30

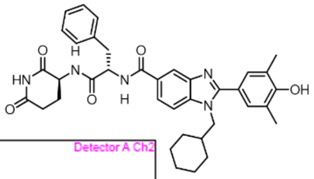

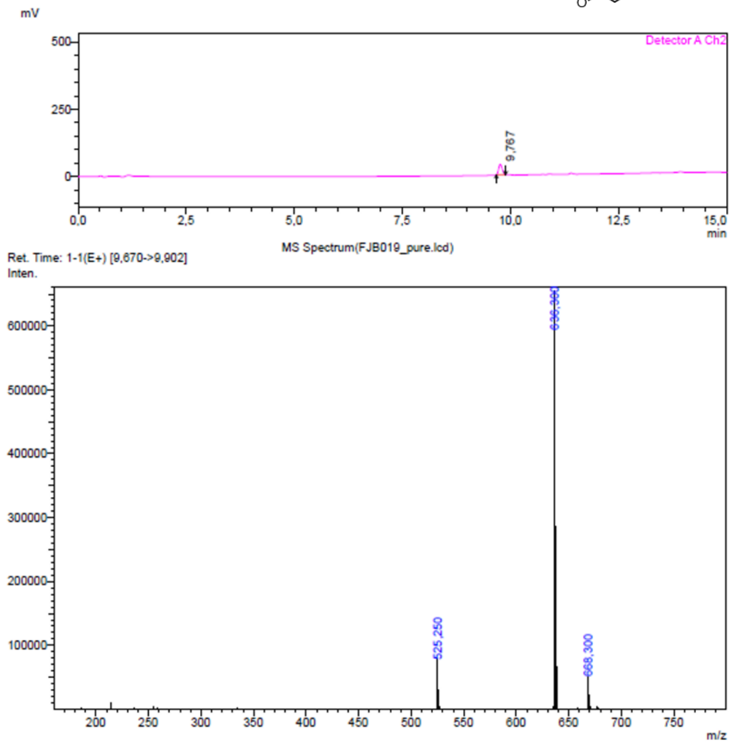

**54:** Exact mass: 746.38; observed $[M + H]^+$: 747.35

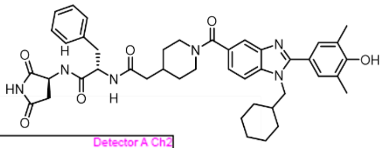

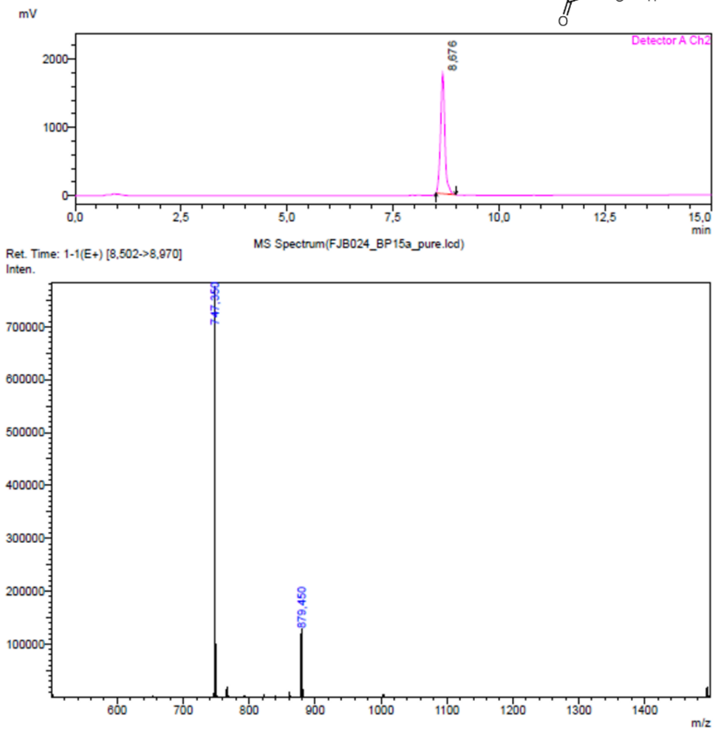

**55:** Exact mass: 859.46; observed $[M + H]^{+}$: 860.50

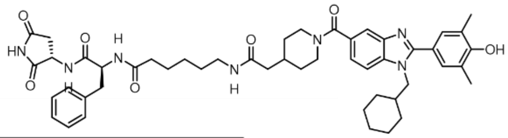

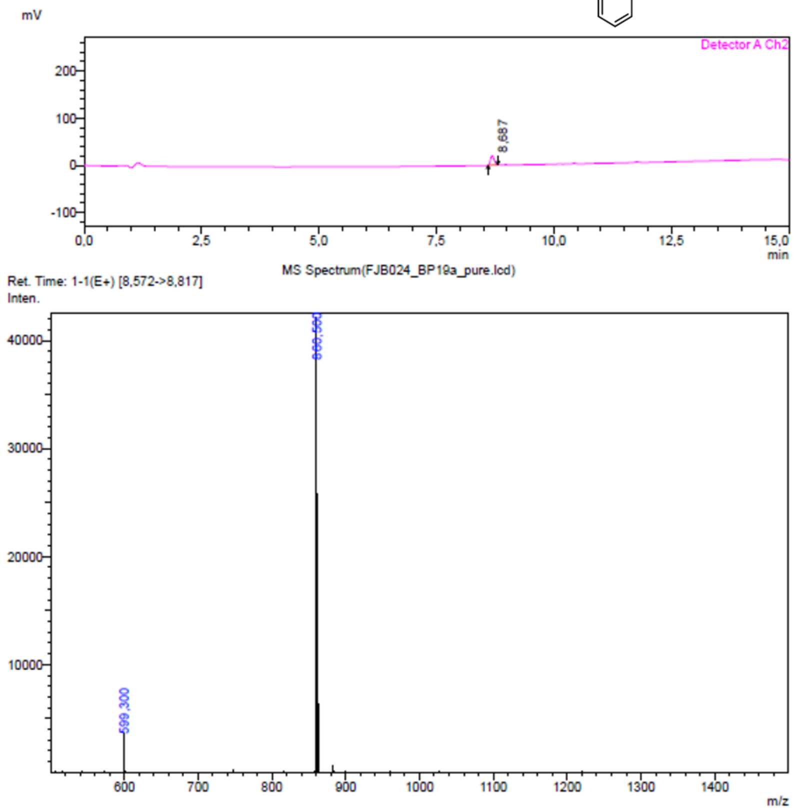

**56:** Exact mass: 749.35; observed $[M + H]^{+}$: 750.35

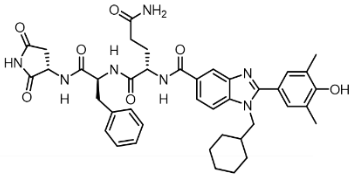

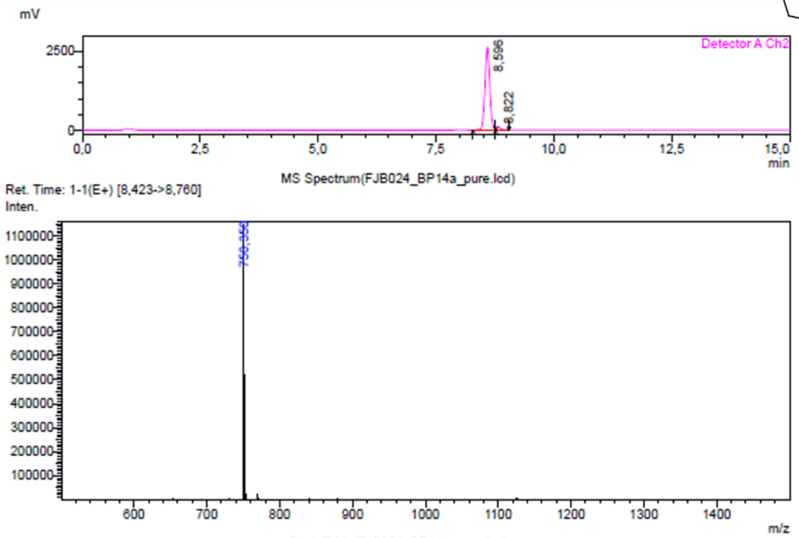

Peak Table(FJB024_BP14a_pure.lcd)

Detector A Channel 2 254nm

| Peak# | Ret. Time | Area | Height | Mark | Conc. | Area% |
|---|---|---|---|---|---|---|
| 1 | 8.596 | 19004883 | 2613249 | | 96.585 | 96.585 |
| 2 | 8.822 | 671866 | 95386 | V | 3.415 | 3.415 |
| Total | | 19676749 | 2708636 | | 100.000 | 100.000 |

**57:** Exact mass: 766.37; observed $[M + H]^{+}$: 767.40

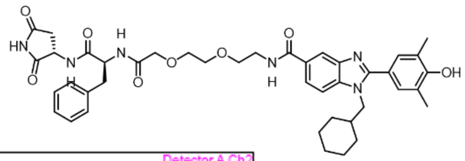

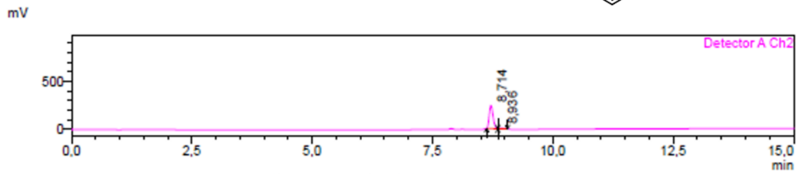

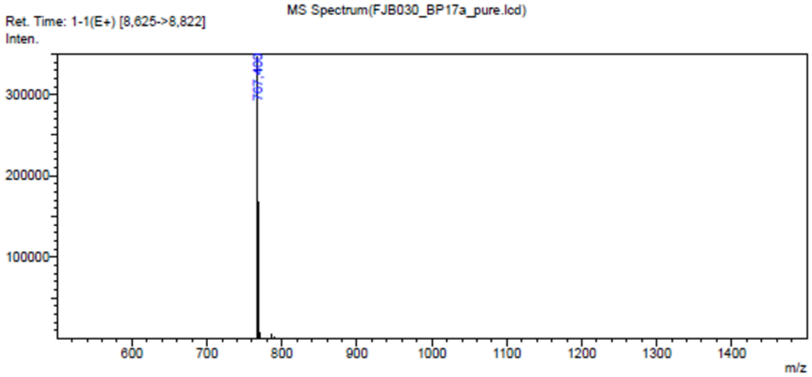

Peak Table(FJB030_BP17a_pure.lcd)

Detector A Channel 2 254nm

| Peak# | Ret. Time | Area | Height | Mark | Conc. | Area% |
|---|---|---|---|---|---|---|
| 1 | 8.714 | 1334332 | 242433 | M | 96.886 | 96.886 |
| 2 | 8.936 | 42888 | 6964 | M | 3.114 | 3.114 |
| Total | | 1377220 | 249397 | | 100.000 | 100.000 |

**58:** Exact mass: 891.45; observed $[M + H]^{+}$: 892.45

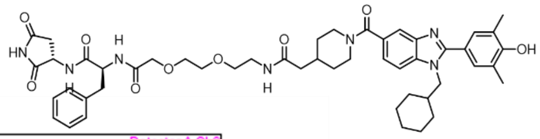

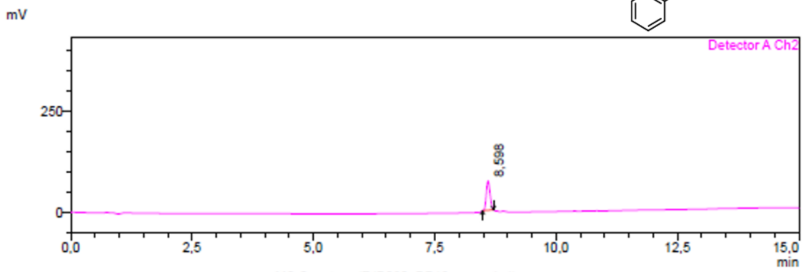

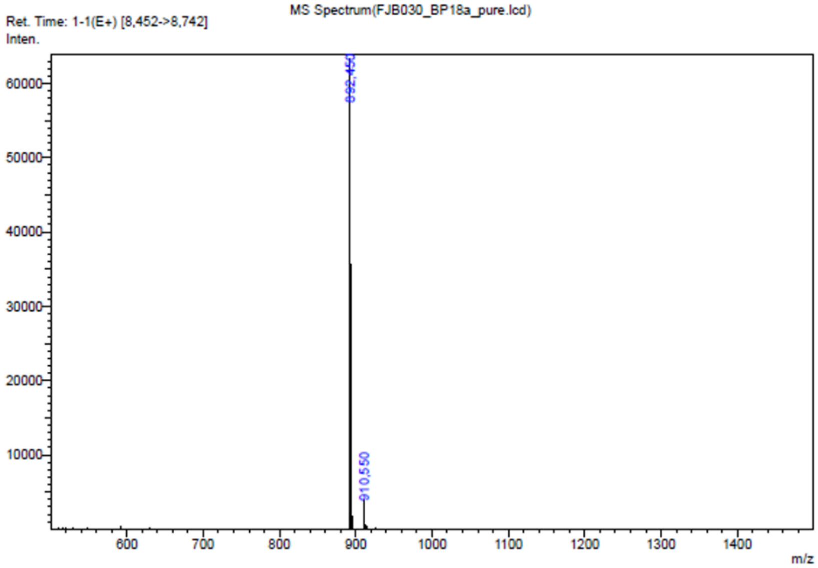

**59:** Exact mass: 803.40; observed [M + H]$^+$: 804.40

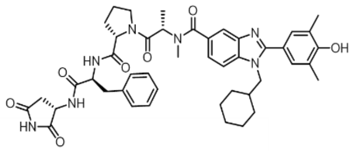

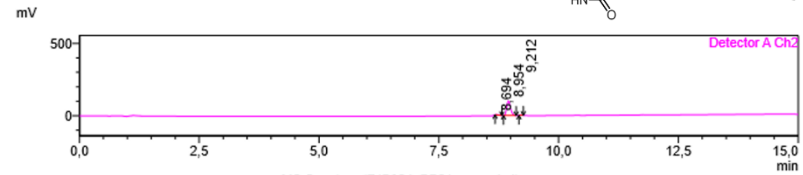

MS Spectrum(FJB024_BP21a_pure.lcd)

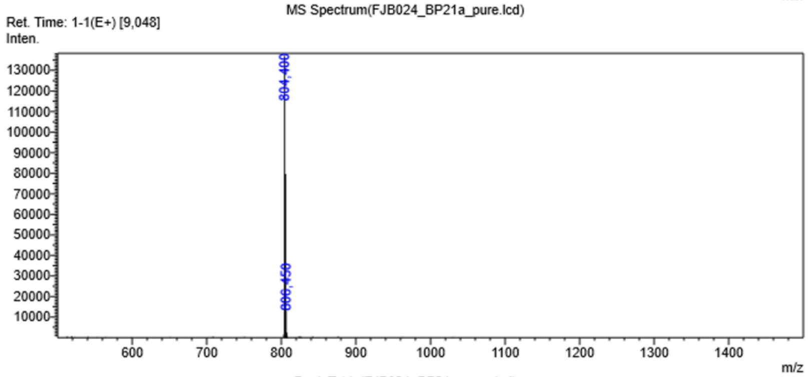

Peak Table(FJB024_BP21a_pure.lcd)

Detector A Channel 2 254nm

| Peak# | Ret. Time | Area | Height | Mark | Conc. | Area% |
|---|---|---|---|---|---|---|
| 1 | 8.694 | 8753 | 1492 | M | 1.512 | 1.512 |
| 2 | 8.954 | 562689 | 100459 | M | 97.221 | 97.221 |
| 3 | 9.212 | 7333 | 2238 | M | 1.267 | 1.267 |
| Total | | 578776 | 104189 | | 100.000 | 100.000 |

**60:** Exact mass: 859.46; observed [M + H]$^+$: 860.50

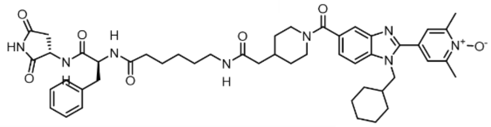

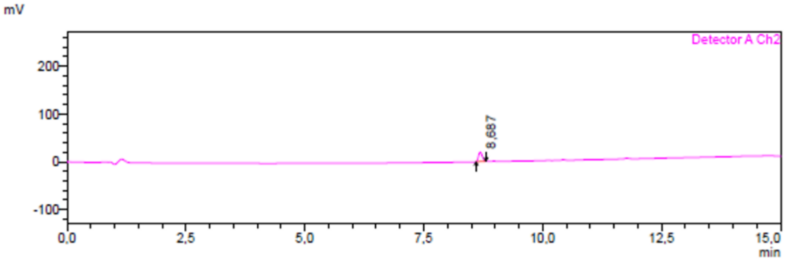

MS Spectrum(FJB024_BP19a_pure.lcd)

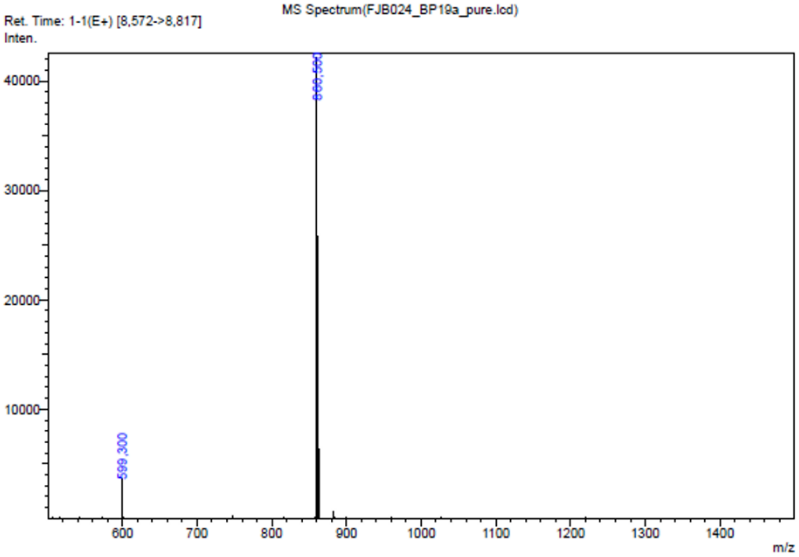

**61:** Exact mass: 804.40; observed $[M + H]^{+}$: 805.40

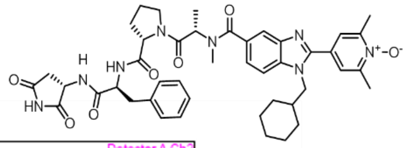

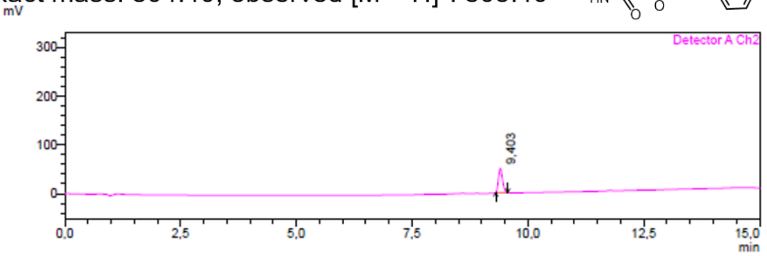

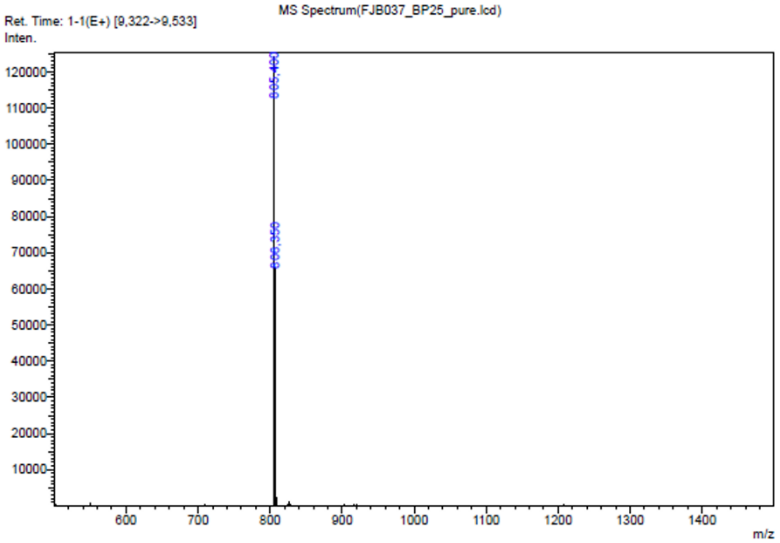

**62:** Exact mass: 845.45; observed $[M + H]^{+}$: 846.40

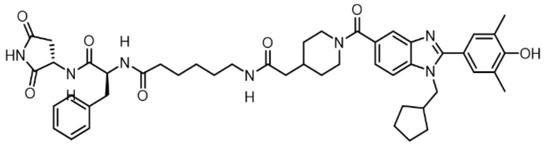

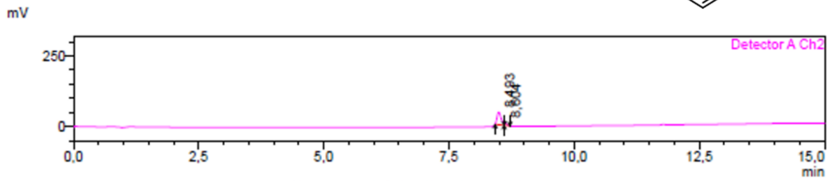

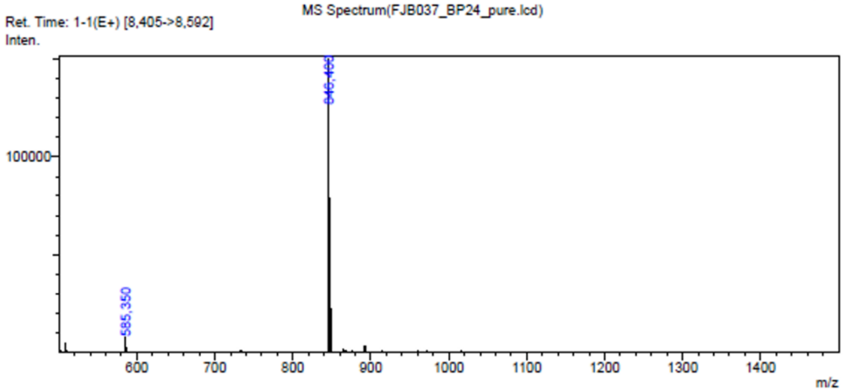

Peak Table(FJB037_BP24_pure.lcd)

Detector A Channel 2 254nm

| Peak# | Ret. Time | Area | Height | Mark | Conc. | Area% |
|---|---|---|---|---|---|---|
| 1 | 8.493 | 255896 | 44393 | M | 89.763 | 89.763 |
| 2 | 8.604 | 29184 | 9915 | V M | 10.237 | 10.237 |
| Total | | 285080 | 54308 | | 100.000 | 100.000 |

**63:** Exact mass: 789.39; observed $[M + H]^+$: 790.40

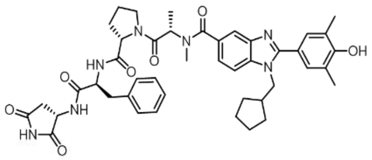

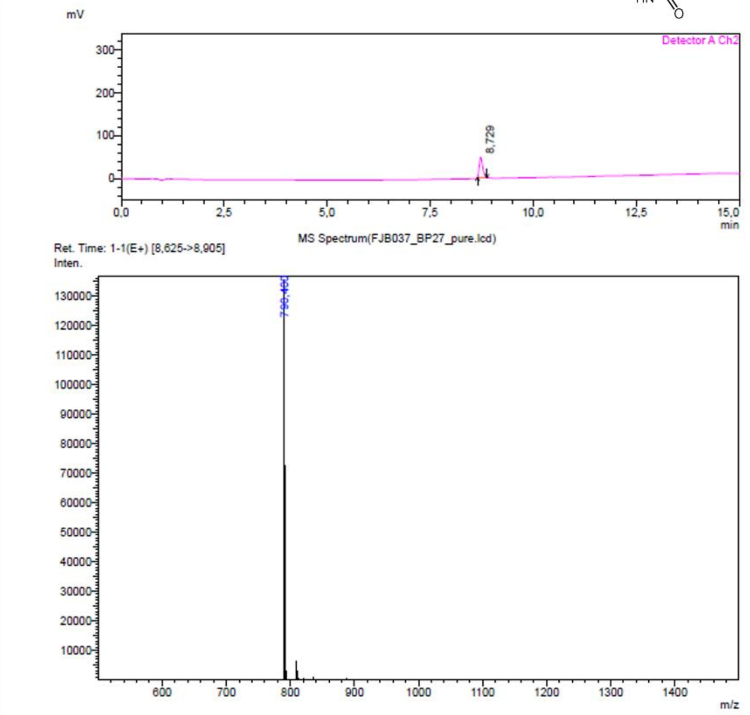

**64:** Exact mass: 873.48; observed $[M + H]^+$: 874.45

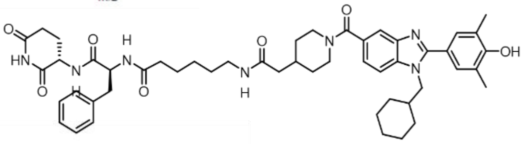

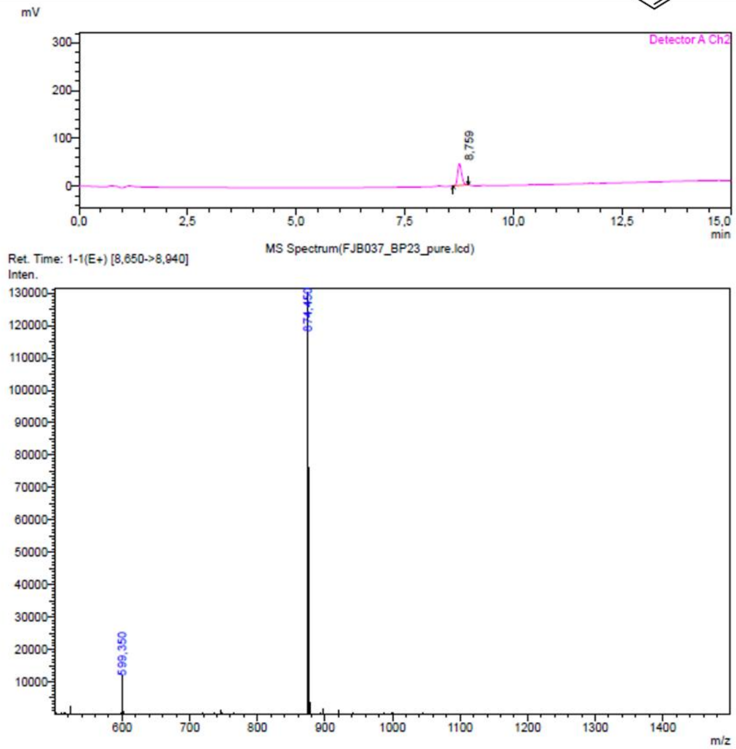

**65:** Exact mass: 817.42; observed $[M + H]^+$: 818.40

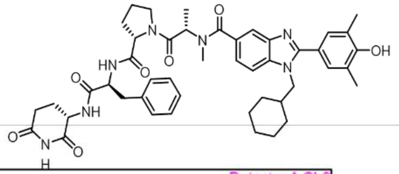

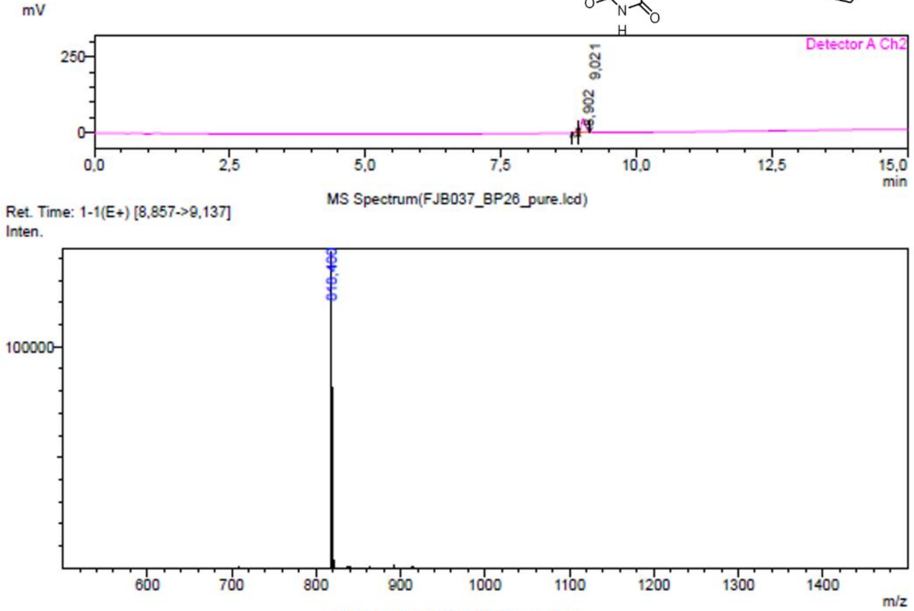

Peak Table(FJB037_BP26_pure.lcd)

Detector A Channel 2 254nm

| Peak# | Ret. Time | Area | Height | Mark | Conc. | Area% |
|---|---|---|---|---|---|---|
| 1 | 8.902 | 21784 | 4776 | M | 7.928 | 7.928 |
| 2 | 9.021 | 252984 | 43484 | V M | 92.072 | 92.072 |
| Total | | 274768 | 48260 | | 100.000 | 100.000 |

**66:** Exact mass: 802.42; observed $[M + H]^+$: 803.40

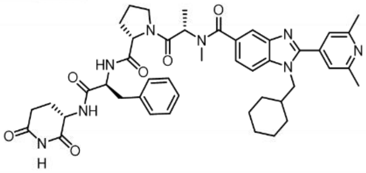

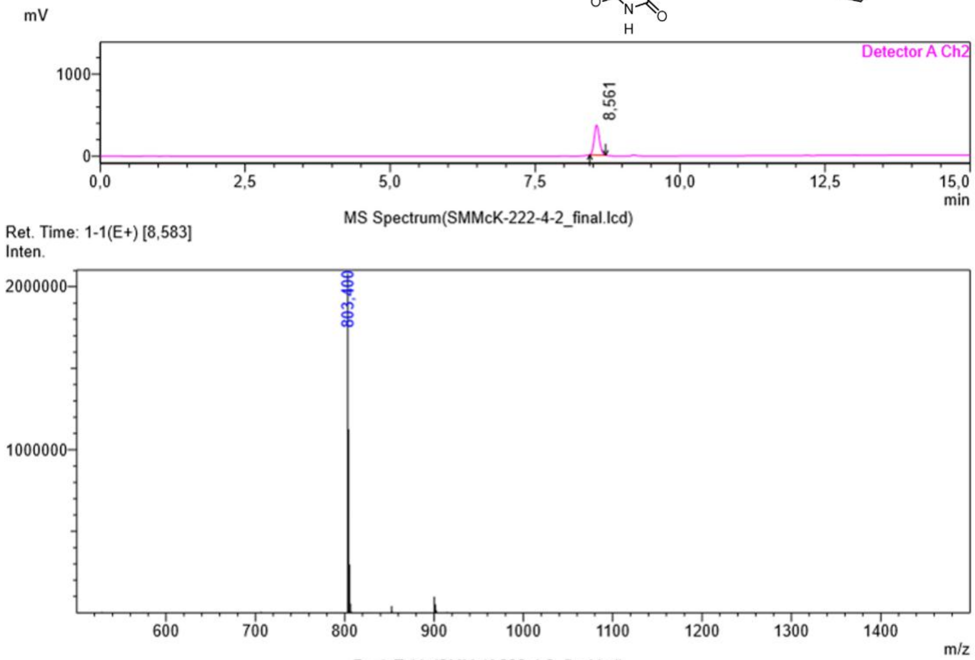

Peak Table(SMMcK-222-4-2_final.lcd)

Detector A Channel 2 254nm

| Peak# | Ret. Time | Area | Height | Mark | Conc. | Area% |
|---|---|---|---|---|---|---|
| 1 | 8.561 | 2289711 | 363177 | M | 100.000 | 100.000 |
| Total | | 2289711 | 363177 | | 100.000 | 100.000 |

**67:** Exact mass: 784.38; observed $[M + H]^+$: 785.35

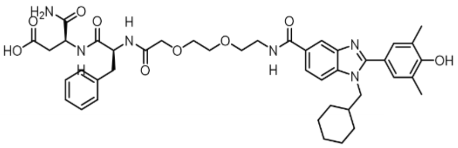

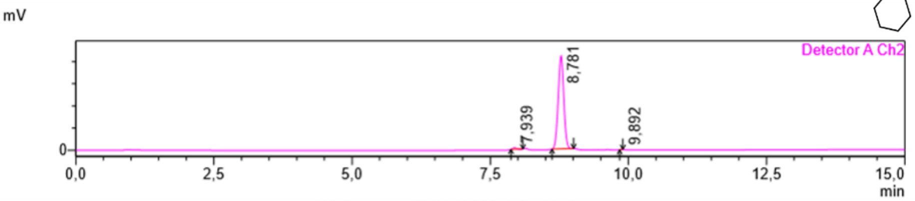

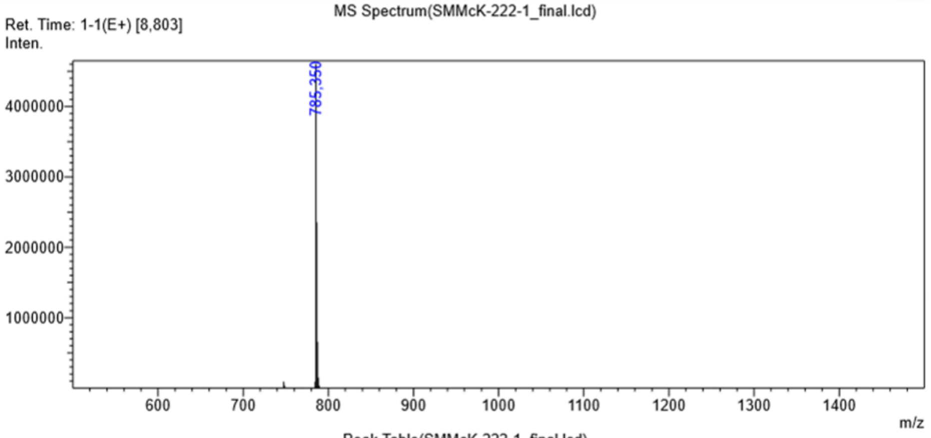

Peak Table(SMMcK-222-1_final.lcd)

Detector A Channel 2 254nm

| Peak# | Ret. Time | Area | Height | Mark | Conc. | Area% |
|---|---|---|---|---|---|---|
| 1 | 7.939 | 126870 | 32854 | M | 0.850 | 0.850 |
| 2 | 8.781 | 14785250 | 2093128 | M | 99.088 | 99.088 |
| 3 | 9.892 | 9144 | 1780 | M | 0.061 | 0.061 |
| Total | | 14921264 | 2127761 | | 100.000 | 100.000 |

**68:** Exact mass: 821.41; observed $[M + H]^+$: 822.35

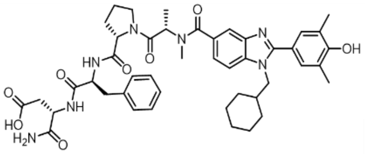

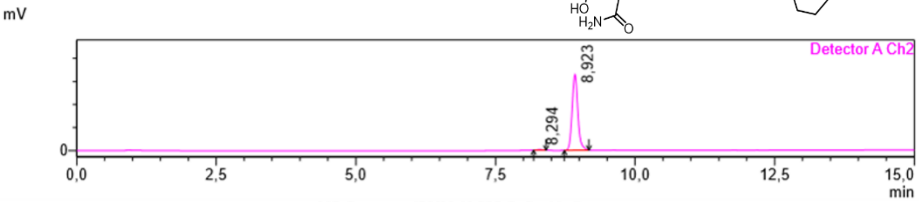

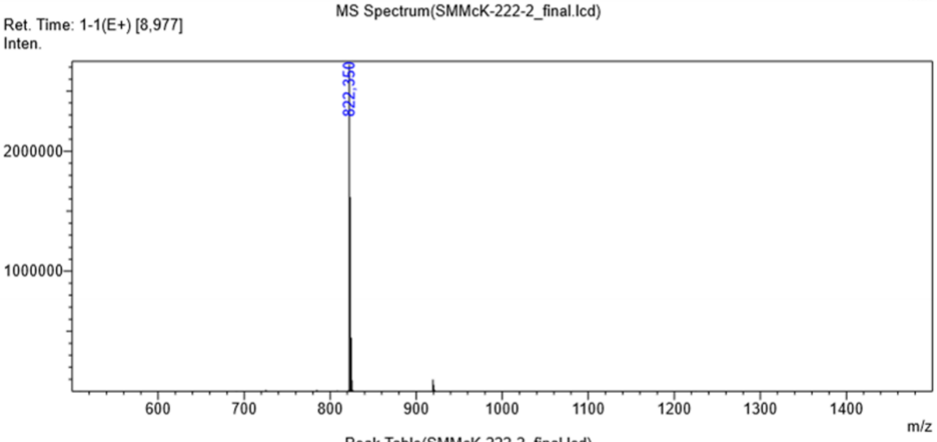

Peak Table(SMMcK-222-2_final.lcd)

Detector A Channel 2 254nm

| Peak# | Ret. Time | Area | Height | Mark | Conc. | Area% |
|---|---|---|---|---|---|---|
| 1 | 8.294 | 38193 | 9559 | M | 0.326 | 0.326 |
| 2 | 8.923 | 11669510 | 1638133 | M | 99.674 | 99.674 |
| Total | | 11707704 | 1647692 | | 100.000 | 100.000 |

**69:** Exact mass: 835.43; observed $[M + H]^+$: 836.40

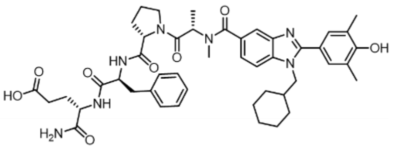

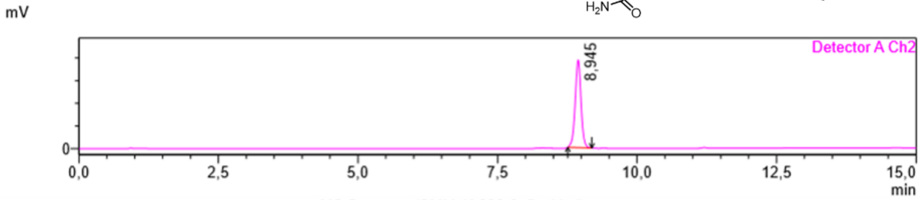

MS Spectrum(SMMcK-222-3_final.lcd)

Ret. Time: 1-1(E+) [8,952]

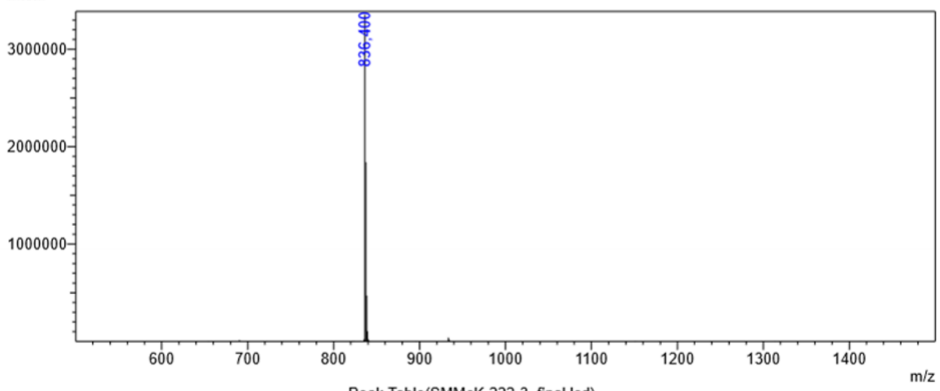

Peak Table(SMMcK-222-3_final.lcd)

Detector A Channel 2 254nm

| Peak# | Ret. Time | Area | Height | Mark | Conc. | Area% |
|---|---|---|---|---|---|---|
| 1 | 8.945 | 14083116 | 1926556 | M | 100.000 | 100.000 |
| Total | | 14083116 | 1926556 | | 100.000 | 100.000 |

**70:** Exact mass: 820.43; observed $[M + H]^+$: 821.40

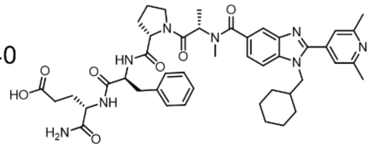

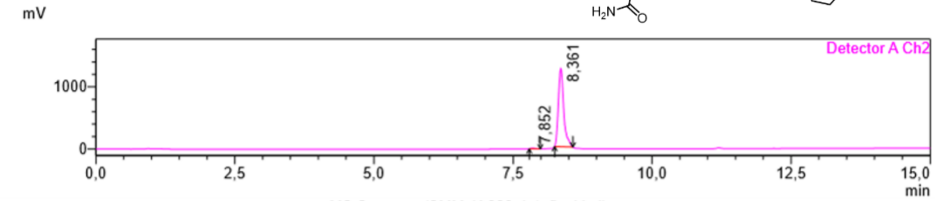

MS Spectrum(SMMcK-222-4-1_final.lcd)

Ret. Time: 1-1(E+) [8,383]

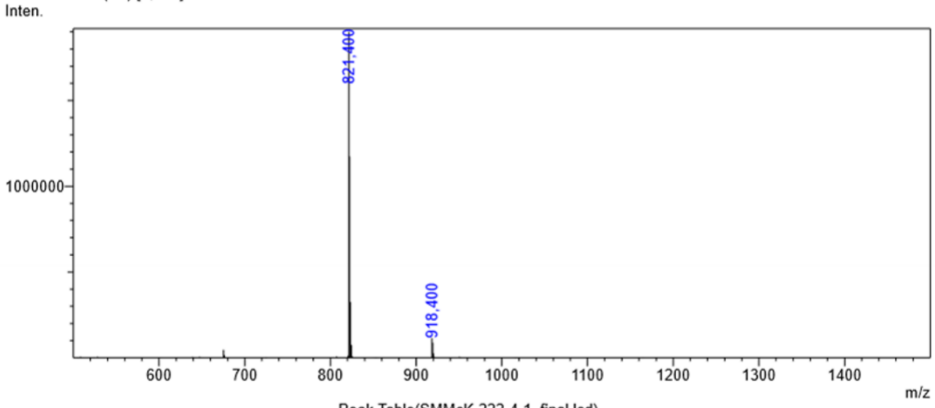

Peak Table(SMMcK-222-4-1_final.lcd)

Detector A Channel 2 254nm

| Peak# | Ret. Time | Area | Height | Mark | Conc. | Area% |
|---|---|---|---|---|---|---|
| 1 | 7.852 | 42991 | 9232 | M | 0.511 | 0.511 |
| 2 | 8.361 | 8363754 | 1253230 | M | 99.489 | 99.489 |
| Total | | 8406745 | 1262462 | | 100.000 | 100.000 |

### $^1$H-NMR Data

**Compound 3**: $^1$H NMR (400 MHz, DMSO-$d_6$) δ 8.76 (d, J = 7.6 Hz, 1H), 8.26 (s, 1H), 8.05 (s, 2H), 7.53 (s, 2H), 7.46 (s, 1H), 7.36 (s, 1H), 7.14 – 6.99 (m, 2H), 6.83 (s, 1H), 4.44 – 4.32 (m, 3H), 2.29 (s, 6H), 2.24 – 2.16 (m, 2H), 2.11 – 1.88 (m, 2H), 1.76 – 1.64 (m, 1H), 1.56 – 1.47 (m, 3H), 1.40 – 1.32 (m, 2H), 1.04 – 0.95 (m, 3H), 0.87 – 0.77 (m, 2H) ppm.

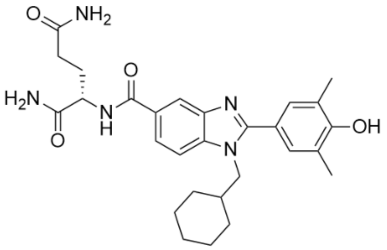

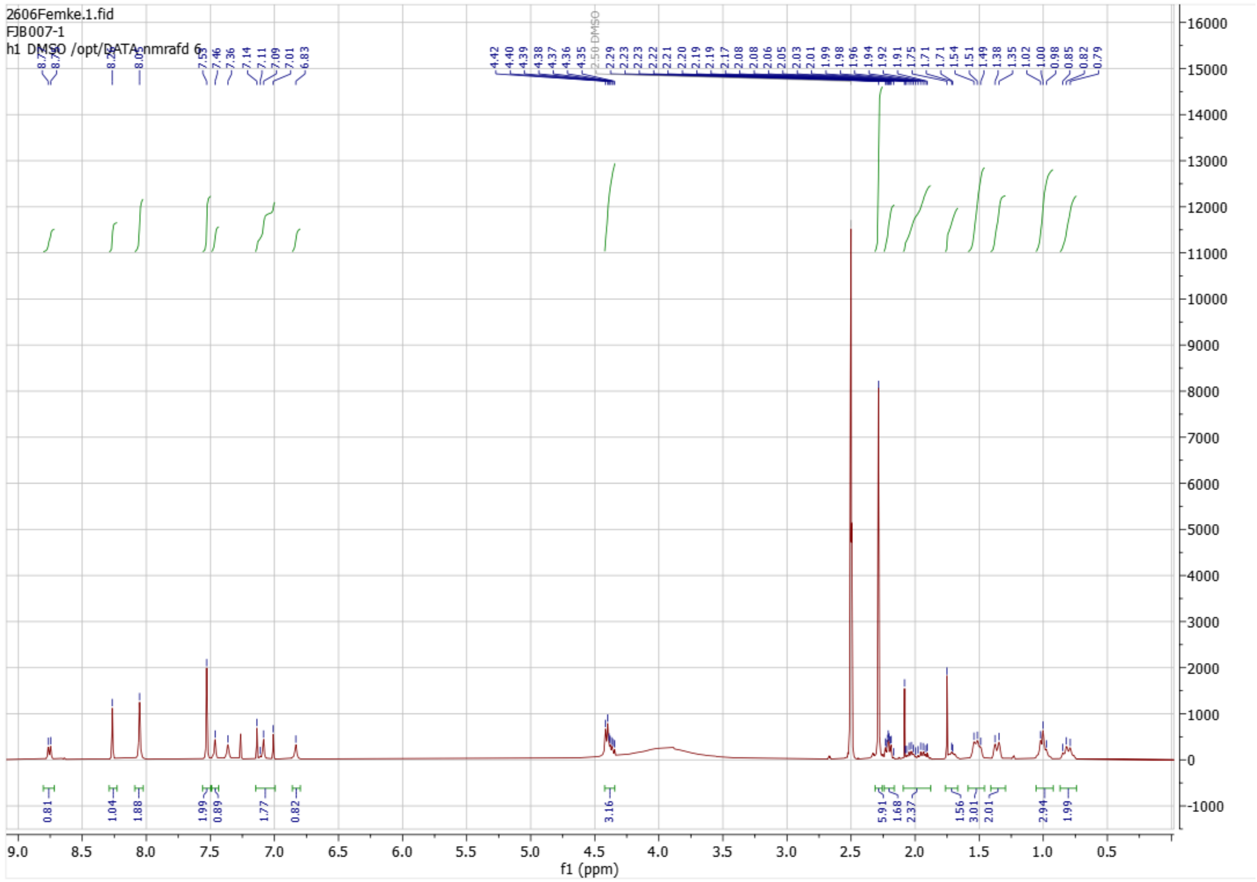

**Compound 26**: $^1$H NMR (600 MHz, DMSO-$d_6$) δ 8.55 (d, J = 7.6 Hz, 1H), 8.28 (s, 1H), 7.91 – 7.86 (m, 3H), 7.81 (d, J = 8.6 Hz, 1H), 7.38 (s, 1H), 7.34 (s, 1H), 7.03 (s, 1H), 6.82 (s, 1H), 4.43 – 4.32 (m, 3H), 2.47 (s, 6H), 2.27 – 2.15 (m, 2H), 2.07 – 1.88 (m, 2H), 1.69 – 1.59 (m, 1H), 1.55 – 1.46 (m, 3H), 1.31 – 1.21 (m, 2H), 1.07 – 0.93 (m, 3H), 0.87 – 0.77 (m, 2H) ppm.

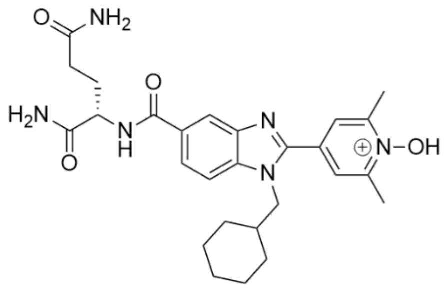

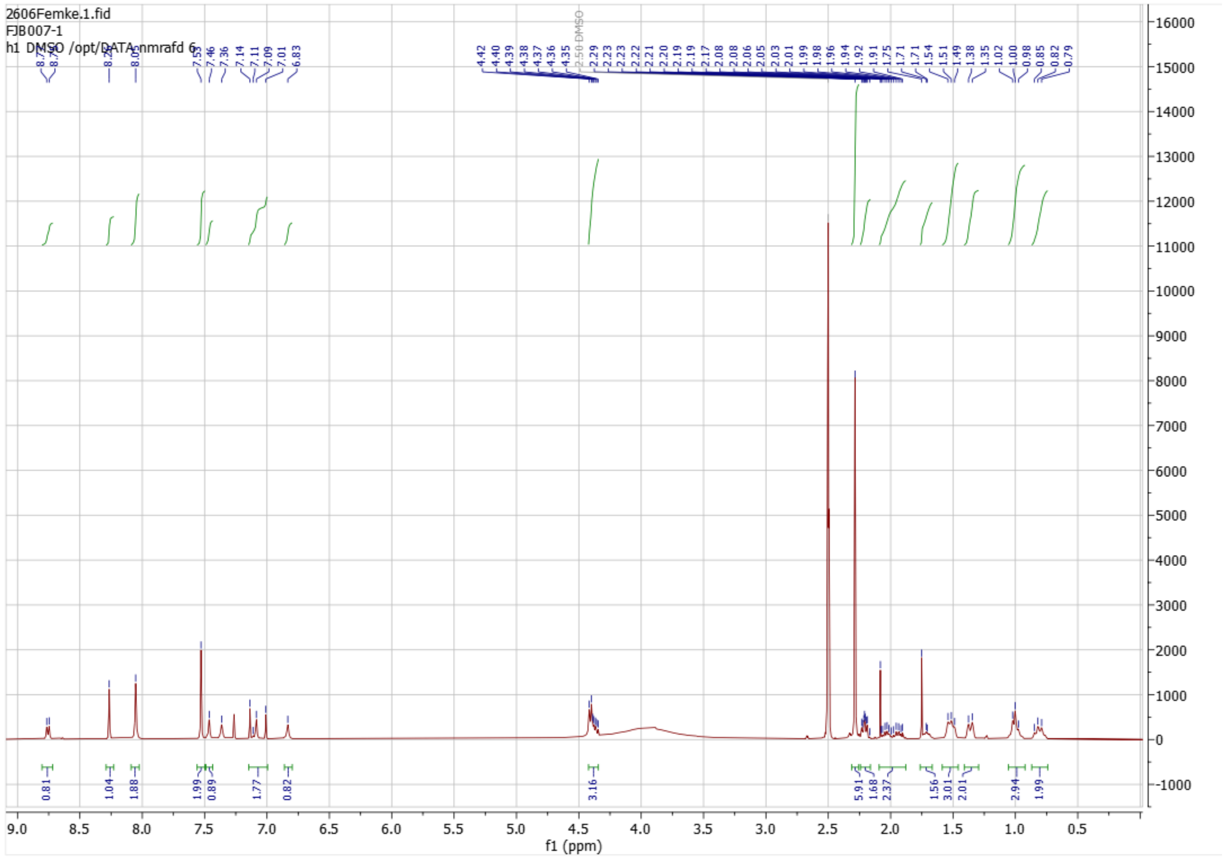

**Compound 51**: $^{1}$H NMR (600 MHz, DMSO-$d_6$) δ 11.24 (d, J = 10.7 Hz, 1H), 8.76 (s, 1H), 8.67 (dd, J = 8.0 Hz, 1H), 8.58 (dd, J = 8.4 Hz, 1H), 8.14 (s, 1H), 7.71 (d, J = 1.5 Hz, 1H), 7.65 (d, J = 8.6 Hz, 1H), 7.38 – 7.32 (m, 4H), 7.25 (d, J = 3.0 Hz, 2H), 7.21 – 7.12 (m, 1H), 4.77 – 4.49 (m, 2H), 4.22 (d, J = 7.4 Hz, 2H), 3.20 – 3.13 (m, 1H), 3.07 – 2.99 (m, 1H), 2.92 – 2.84 (m, 1H), 2.46 – 2.37 (m, 1H), 2.25 (s, 6H), 2.05 – 1.95 (m, 1H), 1.69 – 1.60 (m, 1H), 1.53 – 1.43 (m, 2H), 1.31 – 1.22 (m, 2H), 1.04 – 0.92 (m, 2H), 0.88 – 0.81 (m, 1H), 0.80 – 0.71 (m, 2H).

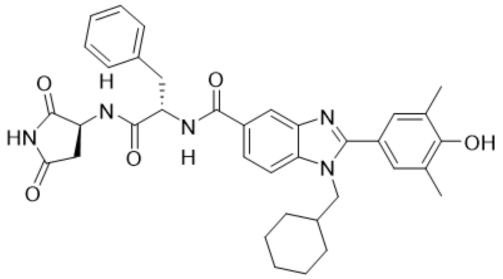

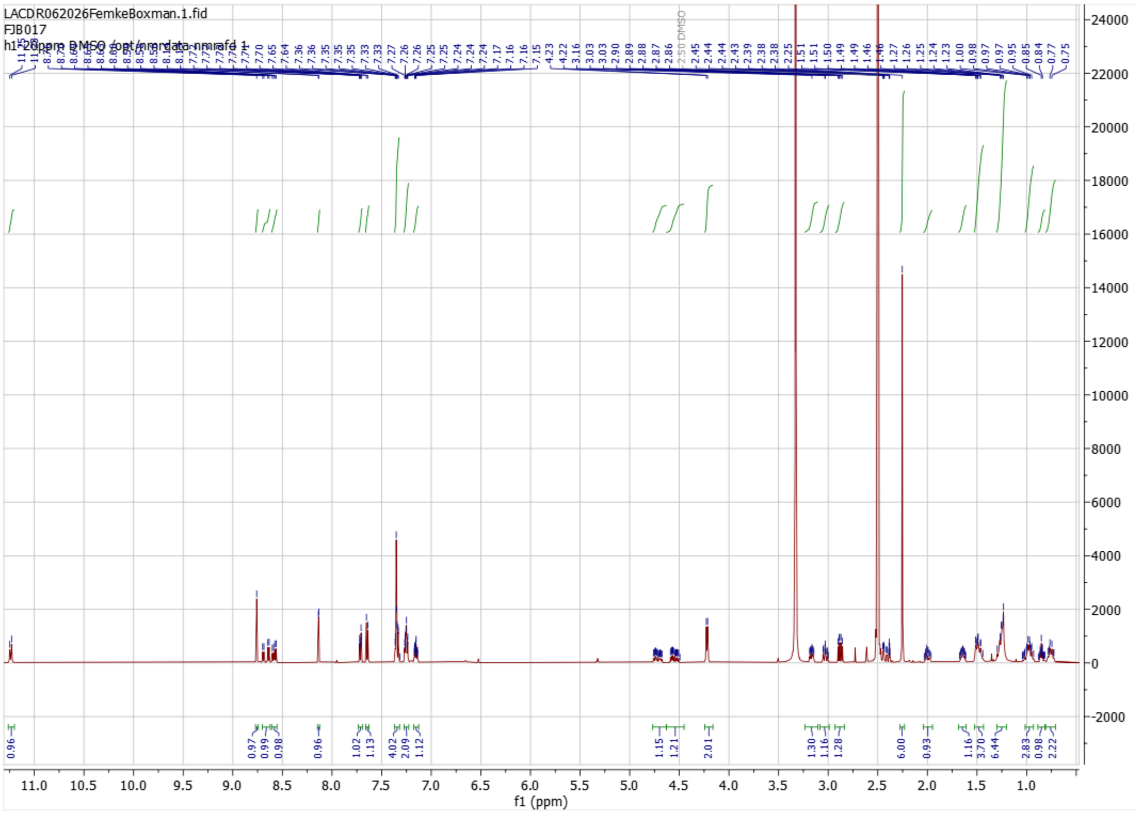

**Compound 53**: $^1$H NMR (600 MHz, DMSO-$d_6$) δ 10.83 (s, 1H), 8.76 (s, 1H), 8.56 (m, 1H), 8.45 (dd, J = 29.8, 8.1 Hz, 1H), 8.11 – 8.05 (m, 1H), 7.74 – 7.67 (m, 1H), 7.67 – 7.60 (m, 1H), 7.41 – 7.32 (m, 4H), 7.27 – 7.24 (m, 2H), 7.19 – 7.13 (m, 1H), 5.37 – 5.25 (m, 1H), 4.82 – 4.73 (m, 1H), 4.73 – 4.60 (m, 1H), 4.21 (d, J = 7.3 Hz, 2H), 4.05 – 3.93 (m, 1H), 3.50 (d, J = 1.2 Hz, 1H), 2.81 – 2.65 (m, 1H), 2.62 – 2.59 (m, 1H), 2.39 – 2.37 (m, 1H), 2.25 (s, 6H), 2.04 – 1.93 (m, 1H), 1.68 – 1.59 (m, 1H), 1.52 – 1.40 (m, 2H), 1.35 – 1.14 (m, 2H), 1.01 – 0.91 (m, 2H), 0.88 – 0.81 (m, 1H), 0.80 – 0.70 (m, 2H).

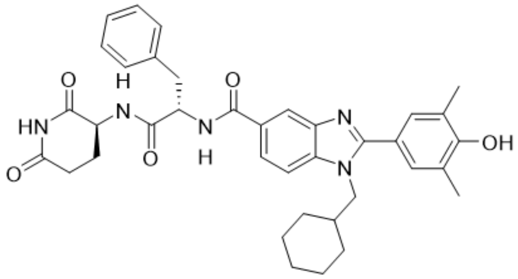

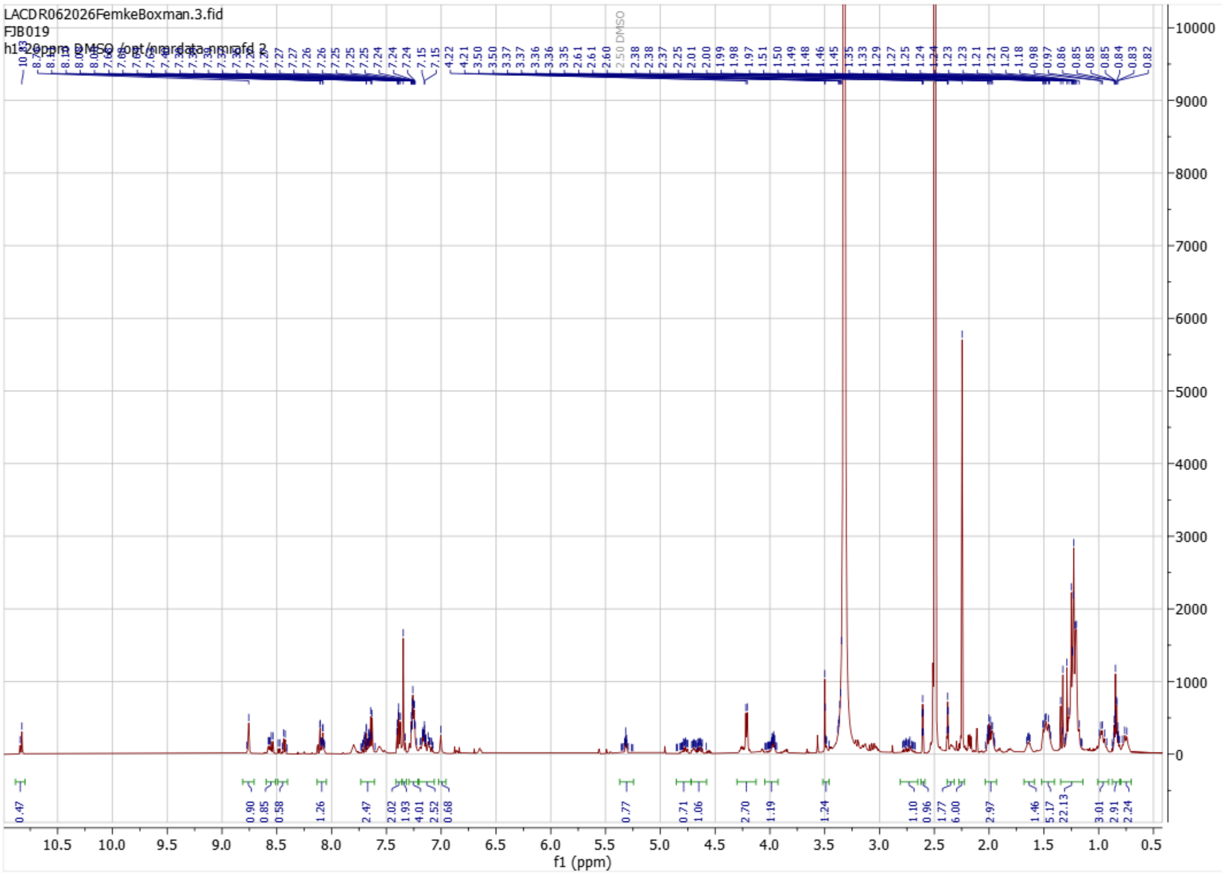

**Compound 59**: $^1$H NMR (600 MHz, DMSO-$d_6$) δ 11.26 (s, 1H), 9.09 (s, 1H), 8.93 (s, 1H), 8.17 (s, 1H), 8.00 (d, J = 8.7 Hz, 1H), 7.87 – 7.75 (m, 1H), 7.75 – 7.65 (m, 1H), 7.41 (s, 2H), 7.32 – 7.14 (m, 5H), 7.09 (s, 1H), 7.01 (s, 1H), 5.48 – 5.39 (m, 1H), 5.37 – 5.27 (m, 1H), 4.47 (d, J = 29.9 Hz, 2H), 4.27 (dd, J = 33.1, 8.3 Hz, 3H), 2.94 – 2.78 (m, 4H), 2.64 – 2.59 (m, 1H), 2.43 – 2.36 (m, 1H), 2.27 (s, 6H), 2.08 – 1.92 (m, 2H), 1.72 – 1.65 (m, 1H), 1.58 – 1.41 (m, 3H), 1.38 – 1.16 (m, 4H), 1.07 – 0.95 (m, 2H), 0.88 – 0.76 (m, 3H).

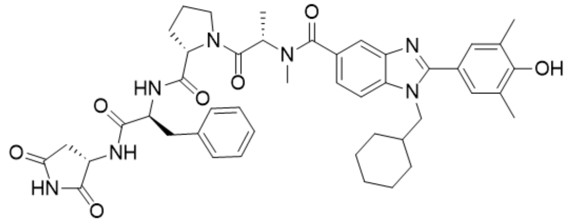

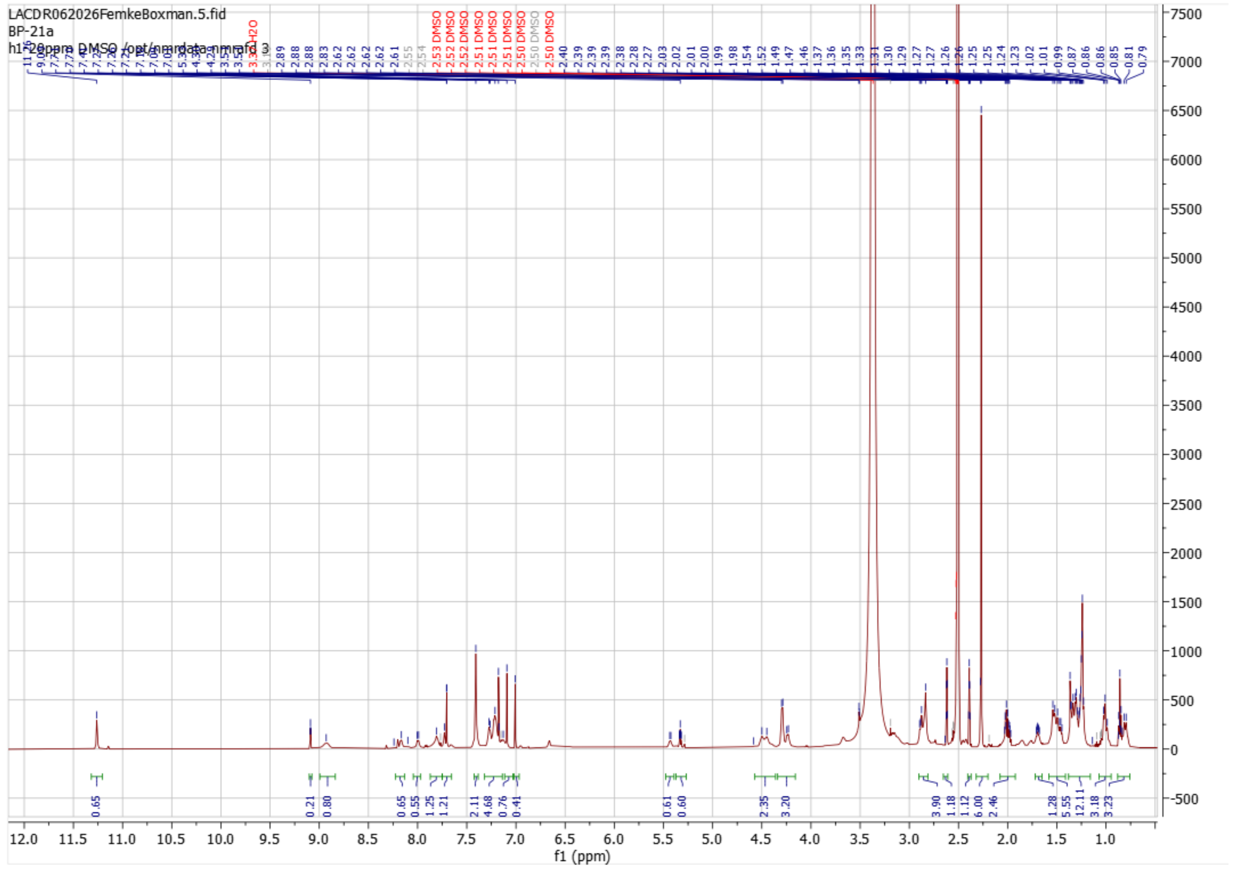

### Biological Data

Uncropped Western blots for compounds **37**- **39**, **48** and **49**

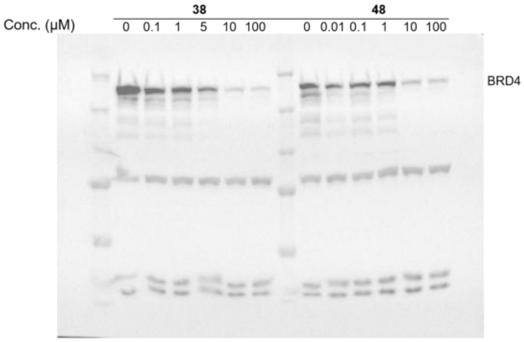

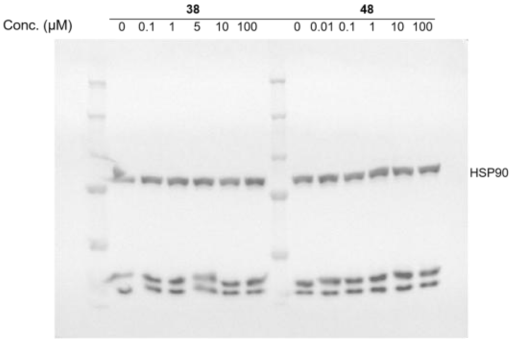

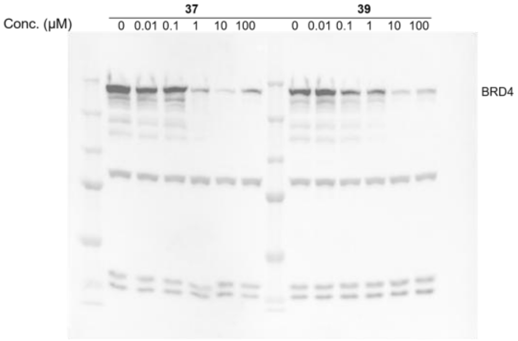

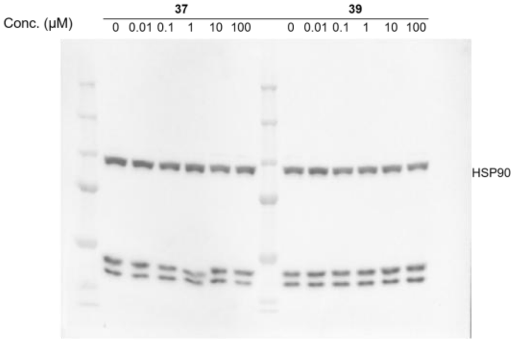

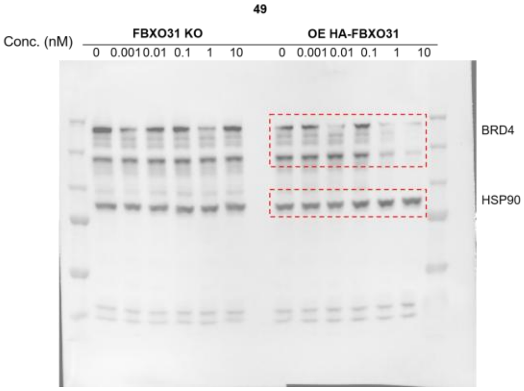

Uncropped Western blots for compounds **51**- **59**

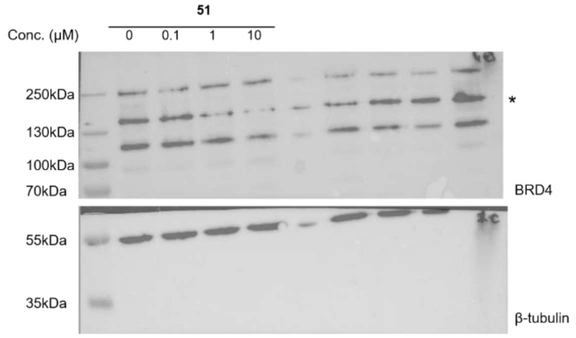

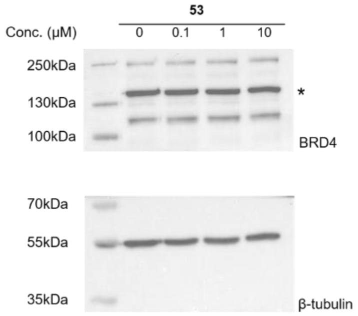

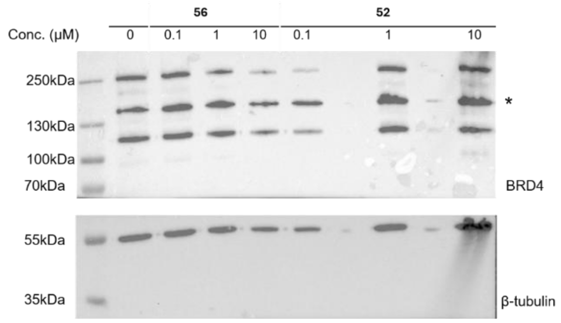

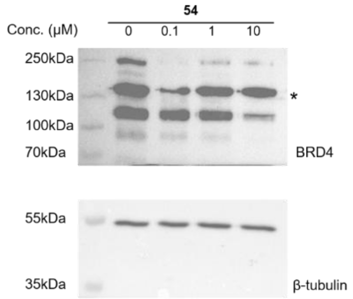

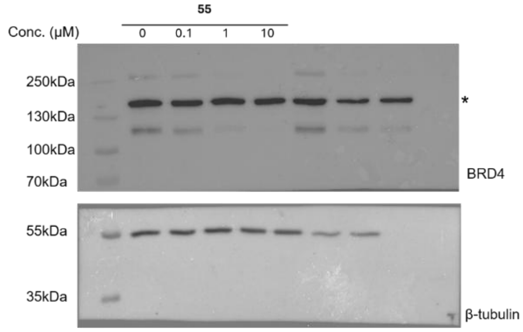

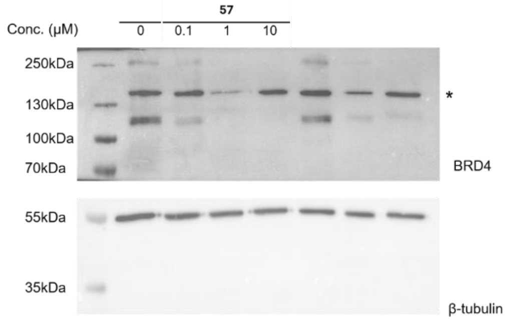

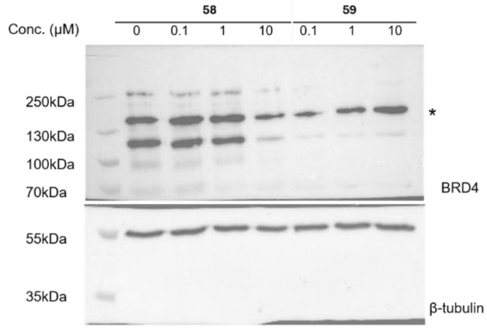

Uncropped Western blots for compounds **60**- **70**

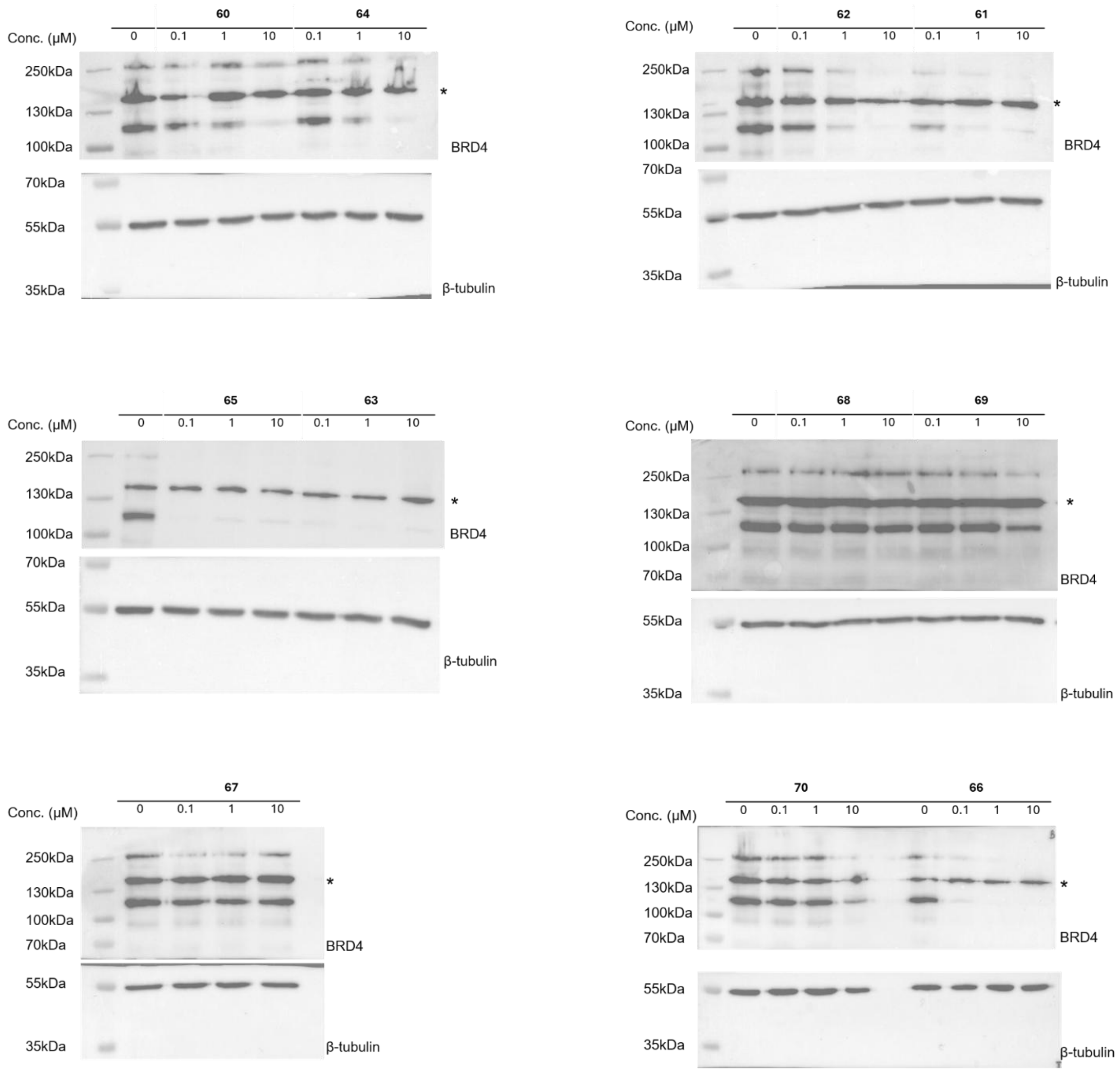

* Indicates unspecific bands
